# The conformational landscape of human transthyretin revealed by cryo-EM

**DOI:** 10.1101/2024.01.23.576879

**Authors:** Benjamin Basanta, Karina Nugroho, Nicholas L. Yan, Gabriel M. Kline, Evan T. Powers, Felix J. Tsai, Mengyu Wu, Althea Hansel-Harris, Jason S. Chen, Stefano Forli, Jeffrey W. Kelly, Gabriel C. Lander

## Abstract

Transthyretin (TTR) is a natively tetrameric thyroxine transporter found in blood and cerebrospinal fluid whose misfolding and aggregation causes transthyretin amyloidosis. A rational drug design campaign identified the small molecule tafamidis (Vyndaqel/Vyndamax) as an effective stabilizer of the native TTR fold, and this aggregation inhibitor is regulatory agency-approved for the treatment of TTR amyloidosis. Despite 50 years of structural studies on TTR and this triumph of structure-based drug design, there remains a notable dearth of structural information available to understand ligand binding allostery and amyloidogenic TTR unfolding intermediates. We used single-particle cryo-electron microscopy (cryo-EM) to investigate the conformational landscape of this 55 kiloDalton tetramer in the absence and presence of one or two ligands, revealing inherent asymmetries in the tetrameric architecture and previously unobserved conformational states. These findings provide critical mechanistic insights into negatively cooperative ligand binding and the structural pathways responsible for TTR amyloidogenesis. This study underscores the capacity of cryo-EM to provide new insights into protein structures that have been historically considered too small to visualize and to identify pharmacological targets suppressed by the confines of the crystal lattice, opening uncharted territory in structure-based drug design.

## Introduction

Transthyretin amyloidosis (ATTR) is a life-threatening degenerative disease associated with the misfolding and aggregation of the transthyretin (TTR) protein, leading to accumulation of aggregates, including amyloid fibrils, in the heart, peripheral and autonomic nerves, brain, and gastrointestinal tract. Hereditary forms of ATTR (ATTRv), linked to mutations in one TTR allele, destabilize the resulting heterotetramers comprising mutant and WT subunits and hasten dissociation, the rate-limiting step of TTR aggregation. Additionally, the aggregation of WT TTR upon aging leads to a life-threatening cardiomyopathy that affects many more patients than ATTRv (ATTRwt), making ATTRwt the third most common amyloid disease behind Alzheimer’s and Parkinson’s^1,2^.

TTR is a transporter of thyroid hormone (T4) and holoretinol-binding protein in blood and cerebrospinal fluid, and there is evidence that TTR may also be involved in nerve regeneration and protein homeostasis^3^. It was previously shown that TTR amyloid formation is initiated by dissociation of native tetrameric TTR, followed by monomer misfolding into a misassembly-prone species, although the details of this process remain the subject of debate^4^. The small molecule tafamidis was found to stabilize the native TTR tetramer by occupying the T4 binding pocket, thereby preventing dissociation, misfolding, and aggregation^5^. Tafamidis-mediated stabilization of TTR was shown to effectively inhibit the amyloidogenesis cascade and stop or slow disease progression, and in 2019 the U.S. Food and Drug Administration approved tafamidis (Vyndaqel/Vyndamax) for treatment of ATTR-associated heart failure.

Despite this successful structure-based drug design campaign, there remains a notable dearth of structural insights into TTR ligand-binding allostery. Moreover, there is not very much structural information on amyloidogenic TTR unfolding intermediates. In this study, we used single-particle cryo-electron microscopy to resolve different conformations that TTR adopted in solution. The TTR protomer natively adopts a β-sandwich fold that oligomerizes as a “dimer of dimers” to form a D2-symmetric homotetramer. Two TTR subunits form a dimer via pairing of the “H” β-strands, which connects the hydrophobic cores of the neighboring β-sandwiches (**Fig. 1A, Supplementary Fig. 1**). The peripheral strands of each β-sandwich fold are “capped” by a structured loop connecting strands D and E (**Fig. 1A**). A pair of dimers then form the tetramer through four contact points between the A-B and G-H loops of opposing subunits, comprising the two T4 binding sites (**Fig 1A, Supplementary Fig. 1**). Tafamidis binds at the two T4 sites to stabilize this weaker dimer-dimer interface (**Supplementary Fig. 1**), thereby inhibiting tetramer dissociation, which is rate-limiting for the misfolding and misassembly of TTR. Despite the binding pockets being symmetrically equivalent according to prior crystallographic structure determinations, the association constant of tafamidis, T4, and a host of other small molecules is substantially higher (as much as 100-fold) for the first binding event compared to the second^5,6^. This has led to decades of speculation and dispute regarding the mechanistic source of this negative cooperativity^7^. The single-particle cryo-electron microscopy (cryo-EM) reconstructions presented in this work shed light on two paramount aspects of TTR biochemistry: the basis of negatively cooperative small molecule binding, and the propensity for amyloidogenic unfolding.

**Figure 1.**
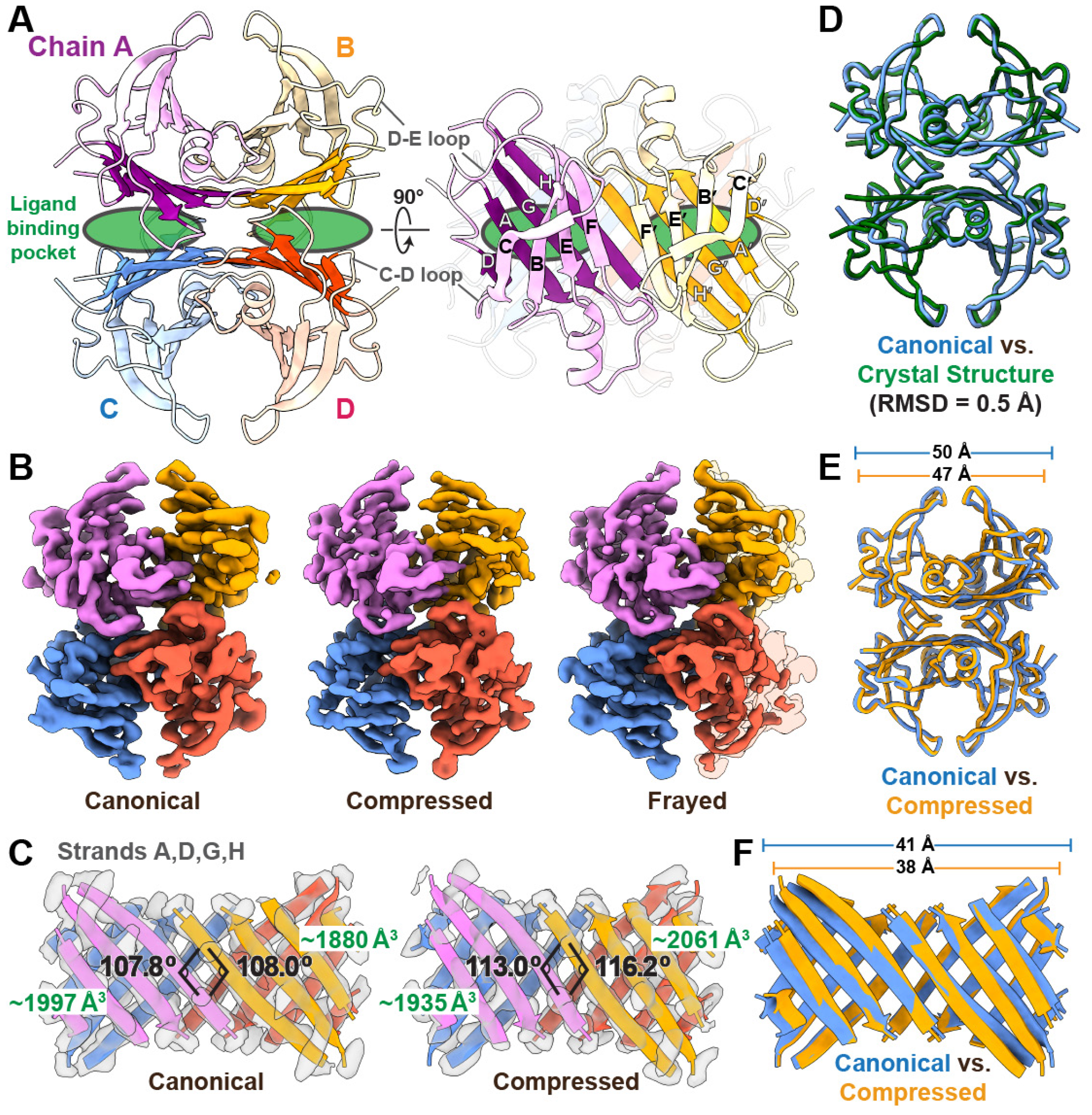
Unliganded tetrameric TTR adopts multiple asymmetric conformations, as determined by cryo-EM. (**A**) A cartoon ribbon representation of TTR (PDB ID: 1ICT) is colored and labeled by protomer. Regions of the β-sheets that form the ligand binding sites (green ovals) are denoted with more saturated coloring. On the right is an orthogonal view where the individual strands of the A-B dimer are labeled. (**B**) Cryo-EM reconstructions of the canonical, compressed, and frayed states of tetrameric TTR (all resolved to *∼*3.3 Å resolution), colored by subunit. To emphasize the disordering of the B and D subunits in the frayed state, a two-fold symmetrized semi-transparent TTR density is underlaid beneath the reconstruction. (**C**) Ribbon representations of β-strands A, D, G, and H, which comprise the binding pocket, are shown with transparent cryo-EM density to highlight the quality of the reconstruction in this region. The different dihedral angles between opposing H β-strand angles in the canonical and compressed states are denoted in black text, and the different pocket volumes corresponding to each ligand binding site are labeled in green text. (**D**) Licorice representations of the peptide backbones of the canonical TTR conformation determined by cryo-EM (blue) fit to the closest matching crystallography-based atomic model identified in the PDB (ID: 4PVN). (**E**) Licorice representation of the peptide backbones of the canonical (blue) and compressed (gold) TTR conformations determined by cryo-EM are superimposed to emphasize the compaction of the structure in the compressed state. (**F**) Ribbon representation of the β-strands shown in (C) are superimposed to emphasize the differences in β-sheet architecture at the dimer-dimer interface.

## Results

### The TTR tetramer is asymmetric, adopting multiple conformations with distinct binding pockets

At 55 kDa, the TTR tetramer is a small target for single-particle cryo-EM analysis, given the theoretical limit of approximately 35 kDa. Initial cryo-EM imaging studies revealed a strong interaction between TTR and the air-water interface, which prevented structure determination of TTR using traditional open-hole TEM grids (**Supplementary Fig. 2A**). To overcome this issue, we used TEM grids modified with a monolayer of crystalline graphene, a technique that has been successful for high-resolution structure determination of other *∼*50 kDa proteins^8^. Notably, these cryo-EM studies provided an opportunity to examine the three-dimensional TTR structure free of crystal packing and enforced symmetry, and thus we did not impose any symmetry constraints to the final reconstructions (**Supplementary Figs. 2-4**).

Unexpectedly, and in stark contrast to prior crystallographic studies, our cryo-EM analyses revealed that unliganded TTR is an asymmetric tetramer in solution. Further, we observed distinct asymmetric conformational states of unliganded TTR, characterized by a shearing motion of the β-sheets along the axis of the weaker dimer-dimer interface bisecting the T4 sites (**Fig. 1B,C**). One of these asymmetric states, resolved to a global resolution of *∼*3.3 μ, most closely resembles previously determined crystal structures of TTR, as indicated by the H β-strand dihedral angles (**Fig. 1D, Supplementary Fig. 5A**), and thus will be referred to as the canonical state. We also resolved a previously unobserved conformation, characterized by an accordion-like compression of the tetramer along the axis bisecting the T4 binding pockets (hereafter referred to as the compressed state, **Fig. 1B,C, Supplementary Movie 1**). The β-strands accommodate this compression by undergoing a shearing rearrangement that results in an increase in the dihedral angle between opposingly directed β-strands (**Fig. 1C,E**).

Intriguingly, we identified a subset of the particles within the compressed state data that lacked density at periphery of the β-sandwiches. We refer to this structure as “frayed,” since its appearance evokes frayed fabric. The fraying of the density in this state is asymmetric, with the density for the C-terminus of strand C and loops flanking strand D being particularly weaker on one side of the tetramer than the other, as evidenced by visual inspection and reflected in the B-factors (**Fig. 1B** and **Supplementary Fig. 6**). We considered the possibility that the loss of density could be arising from partial denaturation at the air-water interface or by graphene interactions, but analysis of 2D class averages and particle orientation distribution suggested otherwise (**Supplementary Text 1**). Thus, we attribute the weakened density to greater mobility and entropy of the structural elements in this region. The observed disordering of strands C, D, and their flanking loops in our structures is consistent with a partially unfolded amyloidogenic state previously hypothesized to be adopted by a TTR monomer upon tetramer dissociation^9^. The involvement of these disordered structural elements in the unfolding and aggregation process is supported by the fact that the most common ATTR-associated mutation, V30M, positioned near the N-terminal end of strand “B”, is located proximal to one of these disordered regions in the frayed state (**Supplementary Fig. 7**). This mutation of a small amino acid with a high β-strand propensity to a bulky one with lower β-strand propensity likely^10^ destabilizes the pairing of strands B and C in a region that we observe to already be labile in wild-type TTR. In the context of a dissociated monomer, a destabilizing V30M mutation would likely further promote unfolding, exacerbating irreversible aggregation, including amyloid formation. Another ATTR-associated mutation, L55P, is likely to abolish pairing between strands D and A to drive unfolding and aggregation through a similar mechanism (**Supplementary Fig. 7**). Observation of this proposed amyloidogenic intermediate in our ligand-free TTR tetramer strongly supports this mechanism of amyloidogenesis, which was initially proposed over 30 years ago^11^.

The asymmetric structural arrangement of the frayed TTR tetramer prompted us to further investigate the asymmetry within the canonical and compressed TTR conformations. Nearly all of the more than 200 available crystallographically-derived TTR structures were based on crystallization in the P2_1_2_1_2 space group, with the dimer as the asymmetric unit and a crystallographic symmetry axis running through the T4 binding pockets (green ovals in **Fig. 1A**), giving rise to binding pockets of equivalent size and shape. In contrast to these crystal structures, the calculated volume of the binding cavities in each of our unliganded TTR tetramer structures are notably different from one another. While the binding pockets within each tetramer differ by *∼*6%, the compressed and frayed states each contain binding pockets that are up to 5% larger in volume than those in the canonical state (**Fig. 1C, Supplementary Fig. 8, Supplementary Text 2**). Together, these data confirm an inherent asymmetry of the native TTR tetramer assembly that, combined with conformational variation, gives rise to a range of structurally and chemically distinct T4 binding pockets, which may explain the previously established differences in binding affinity of ligands for the two TTR binding sites, as well as the capacity of TTR to bind diverse small molecule structures.

### Asymmetric cryo-EM structures of TTR with ligand in both binding pockets

We sought to further investigate by cryo-EM how ligand binding to both T4 binding sites influences the TTR tetramer. For this, we examined TTR tetramers that were stabilized by co-valent attachment of a stilbene substructure to one of the two Lys15 ε-amino groups within both T4 binding pockets (see **Supplementary Fig. 9** for a line drawing of the (Stilbene)_2_-TTR conjugate and **Supplementary Text 3**)^12^. Crystallography has historically prevented visualization of asymmetry between the two different TTR binding sites, given that the crystallographic symmetry axis runs through the T4 binding pockets. This is compounded by degeneracy that arises from each binding site (**Fig. 1A, Supplementary Fig. 1**, showing one site formed by chain A and C, and the other by chains B and D) due to the inherent two-fold symmetry of the binding pockets. Our cryo-EM image analysis workflow enabled us to examine any asymmetry in the organization of the subunits that comprise the TTR tetramer, as well as in the positioning of the tethered stilbene substructures within the two T4 binding sites (**Fig. 2 and Supplementary Figs. 10-12**).

**Figure 2.**
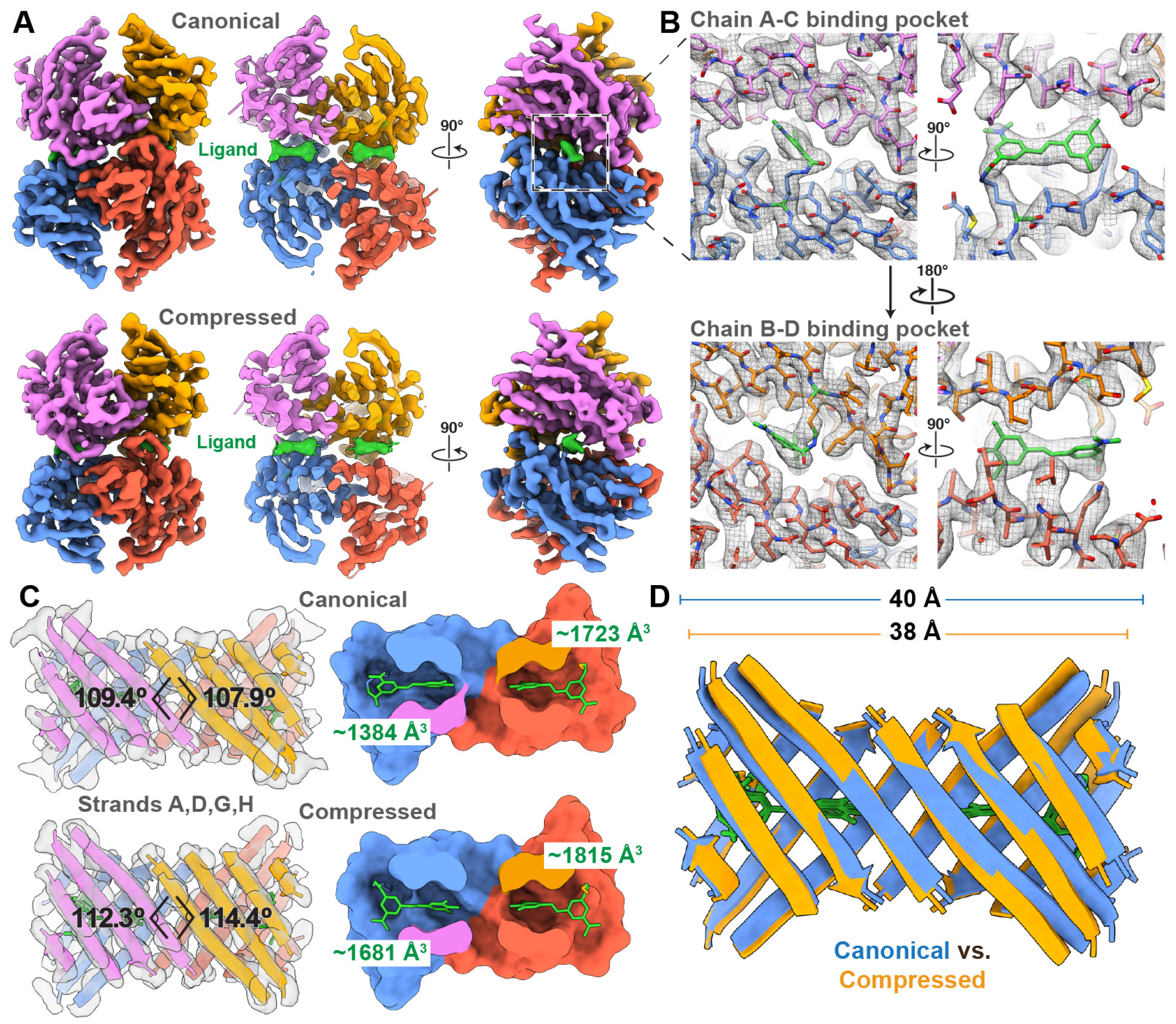
Asymmetric binding pose of stilbene tethered to both TTR binding pockets. (**A**) Orthogonal views of the cryo-EM reconstructions of the canonical (above) and compressed (below) conformations of TTR, resolved to *∼*2.7 and *∼*3.0 Å resolution, respectively, colored by subunit as in Fig. 1. The center image is a cross-sectional view of the cryo-EM density showing the density corresponding to the covalently bound ligand (green). (**B**) The cryo-EM density of each stilbene moiety in the binding pockets of the canonical state reconstruction is shown as a mesh with the atomic model shown as a stick model. The stilbene covalently attached to Lys15 of chain C in the A-C binding pocket (top panel) is better resolved as a single conformation than the stilbene in the B-D binding pocket (bottom panel). (**C**) Ribbon representations of β-strands A, D, G, and H along with the stick representation of the bound stilbene ligand as a green stick representation are shown with transparent cryo-EM density to highlight the quality of the reconstruction in this region. The different H β-strand dihedral angles in the canonical and compressed states are denoted. To the right, a cutaway view of the surface representation of the ligand binding pockets with the stilbene ligand rendered a green stick representation is shown, with the different pocket volumes denoted in green text. (**D**) Ribbon representation of the β-strands shown in (C) are superimposed to emphasize the differences in β-sheet architecture at the dimer-dimer interface.

We expected that covalent tethering of the stilbene ligand within both binding pockets would stabilize a single symmetric conformation, structurally consistent with prior TTR crystallographic studies. Surprisingly, our cryo-EM data revealed that the (Stilbene)_2_-TTR conjugate similarly adopts the asymmetric canonical and compressed conformations as previously observed in the unliganded TTR dataset (**Supplementary Fig. 5B**). However, we were unable to identify the presence of the frayed state in the (Stilbene)_2_-TTR dataset, indicating that ligand binding in both T4 binding sites stabilizes the C and D strands of the tetramer. Notably, we were able to unambiguously distinguish the precise asymmetric pose of the stilbene ligand covalently linked to one of the two Lys15 ε-amino side chains in the smaller binding pocket (comprising subunits A and C) in the canonical state (**Fig. 2A-B, Supplementary Fig. 13**). We also noted that the quality of the covalently bound stilbene density worsens as the size of the binding pocket increases, indicating that the ligand is more flexible in larger binding pocket made up by the B and D subunits (**Fig. 2B, Supplementary Fig. 14**). This observation is supported by molecular dynamics simulations showing greater mobility of the ligand and nearby backbone in the larger T4 pocket of the canonical conformer (**Supplementary Fig. 15, Supplementary Text 4**). Although the pockets in the (Stilbene)_2_-TTR tetramers had a smaller volume than in the corresponding unliganded TTR conformers, the difference is likely due to distinct side chain positions in the presence or absence of ligand, as the inter-chain distances around the pocket are preserved whether liganded or not (**Supplementary Figs. 8 and 14**). These cryo-EM structures confirm that even in the presence of a covalently bound stabilizing ligand at both of the T4 sites, the TTR tetramer is an asymmetric entity that adopts different conformations, and that variability in the size of the T4 binding pockets correlates with the flexibility of the bound ligand.

### Cryo-EM structures of TTR bound to a single ligand

One of TTR’s most prominent characteristics is the negative cooperativity with which it binds small molecule ligands. With the unliganded and (Stilbene)_2_-TTR conjugate structures in hand, we next aimed to shed light on the mechanism of negative cooperativity by imaging a TTR complex engineered to produce a homogeneous population of TTR tetramers disulfide-tethered to a biarylamine ligand in only one of the two T4 binding sites^6^. Thus we synthesized ((biarylamine-FT_2_-WT)_1_(C10A)_3_) TTR (see **Supplementary Fig. 16)**, hereafter referred to as single-liganded TTR (FT_2_ = double FLAG-tag.) Briefly, we synthesized and purified a (FT_2_-WT)_1_(C10A)_3_ TTR tetramer, wherein only the single FT_2_-WT subunit harbors a Cys-10 residue that was quantitatively labeled with 2-((3,5-dichloro-4-(2-(2-(2-(pyridin-2-yldisulfaneyl)ethoxy)ethoxy)ethoxy)phenyl)amino)benzoate to covalently position a 2-(3,5-dichlorophenylamino)benzoic acid stabilizer in only one of the two TTR T4 binding sites. Covalent occupancy of one T4 site with a stabilizing ligand prevents subsequent scrambling of subunits by tetramer dissociation followed by subunit exchange^6^. Accordingly, our cryo-EM reconstructions of this sample contained ligand density exclusively present in one binding pocket (**Fig. 3, Supplementary Figs. 17-21**). Surprisingly, we did not observe the canonical state in the single-liganded TTR sample, instead only observing the compressed and frayed states (**Fig. 3**). As observed in the unliganded TTR dataset, fraying is only seen in the presence of a compressed sheet state, indicating fraying is dependent on sheet compression (see **Supplementary Text 5**). This relationship between disordering of certain regions and the tertiary organization of the sheets is further supported by previously published NMR studies, where the mutants V30M and L55P (**Supplementary Fig. 7**), located at the peripheral edge of the β-sandwich, affect the chemical environment of the rest of the β-sheet^3^. In this context, the region around β-sheet “D” could be thought of as a linchpin that, once disengaged, leads to sheet compression and the observed compressed state (see **Supplementary Text 5**).

**Figure 3.**
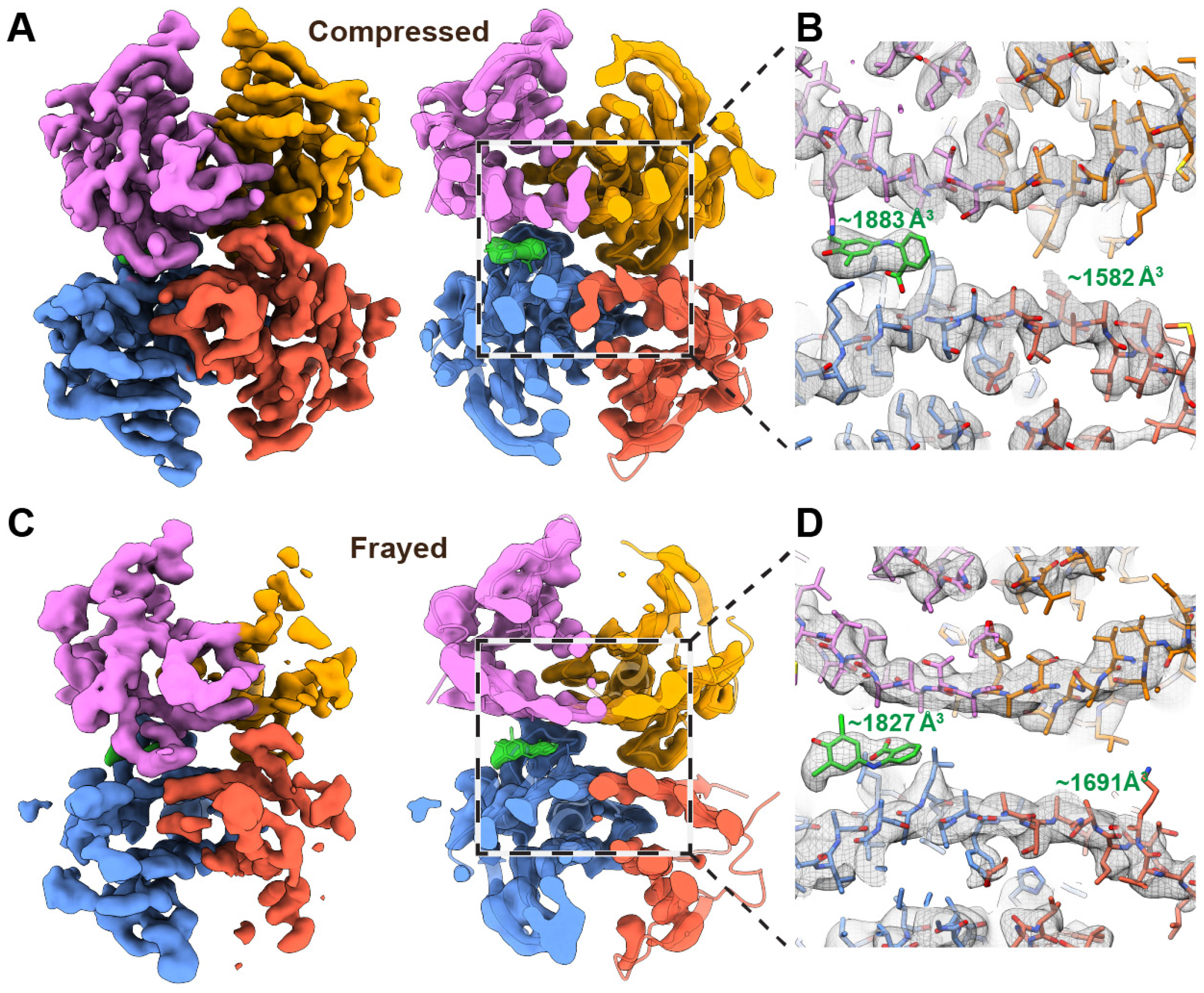
Asymmetric single-liganded TTR structures determined by cryo-EM. (**A**) Cryo-EM reconstructions of the (biarylamine-FT_2_-WT)1(C10A)3 TTR compressed state (determined to *∼*3.4 Å resolution), colored by subunit. On the right is a cross-sectional view of the cryo-EM density showing the density corresponding to the single bound ligand (green) in the A-C binding pocket, while the B-D binding pocket is empty. (**B**) The cryo-EM density corresponding to the area denoted by the dashed box in (A) is shown as a mesh with the atomic model shown as a stick model. The pocket volume of each binding site is denoted in green text. (**C**) Cryo-EM reconstruction of the (biarylamine-FT2-WT)1(C10A)3 TTR compressed frayed state (determined to *∼*4.1 Å resolution), colored by subunit. On the right is a cross-sectional view of the cryo-EM density showing the density corresponding to the single-bound ligand (green) in the A-C binding pocket, while the B-D binding pocket, which is associated with the less-ordered protomers, is empty.

The previously published crystal structure of this (biarylamine-FT_2_-WT)_1_(C10A)_3_ TTR contains ligand density in both pockets due to averaging across the crystal lattice, and TTR is in an intermediate conformation between canonical and compressed, with opposing H strand dihedral angles of 105^*°*^ and 110^*°*6^. Considering the asymmetry distinguishing the two T4 binding sites observed across the seven reconstructions of TTR in this study, it is evident that the TTR pockets are inherently asymmetric and malleable. Subtle changes in the positions of the β-strands within the TTR tetramer are linked to the varied volume and chemical environments of the T4 binding sites, explaining TTR’s ability to bind a host of distinct small molecule structures with varying affinities (see **Supplementary Text 6**). In this context, we speculate that the higher overall symmetry of the canonical state, as quantified by the similarity of H-strand cross angles across the dimers (**Supplementary Fig. 5**), is responsible for its prevalence in crystallographic studies.

## Discussion

Many homo-oligomeric proteins bind their ligands with negative cooperativity, a phenomenon that has been proposed to endow these proteins with the capacity to respond to a range of ligands that are present at widely varying concentrations. Accordingly, TTR is known to bind a diverse array of chemicals, a characteristic that has been proposed to enable the tetramer to function as a broadly functioning detoxification agent^13^. Negative cooperativity is generally understood in terms of the Koshland-Némethy-Filmer sequential binding model, in which the first ligand binding event allosterically induces asymmetry in the complex that reduces compatibility for a second binding event. While it is generally assumed, however, that unliganded homooligomers such as TTR are symmetric in nature, we show that structural asymmetry with distinct ligand binding pockets is inherent to the TTR tetramer, with or without bound ligands. Thus, the entropic penalty associated with ligand binding is likely to be different for each binding pocket, both within a single tetramer and across conformational states.

The TTR protomers comprising the ligand-bound pocket of our single-liganded TTR reconstructions are better ordered than the protomers forming the unoccupied pocket, as evidenced by B-factors in both the compressed and frayed conformations (**Supplementary Fig. 21**). Given that a frayed state is observed in both the single-bound TTR and unliganded TTR datasets, but not in the (Stilbene)_2_-TTR conjugate dataset, we reason that the second ligand binding event has a generally less negative Δ G due to a greater loss of chain entropy upon binding. Our unliganded and (biarylamine-FT_2_-WT)_1_(C10A)_3_ TTR datasets suggest that the disordered regions associated with the frayed state can become ordered, but we expect this ordering would incur an entropic cost. As outlined in **Fig. 4**, we propose that the initial ligand binding event would occur in the ordered half of TTR without incurring this entropic penalty, while the second binding event must be coupled to substantial ordering, especially in the region around the “D” β-strand. Prior thermodynamic experiments support this hypothesis, in that the entropic contribution to the binding Δ G is generally less favorable for the second ligand binding event^3,14,15^. The strongly negatively cooperative binding of ligands to TTR in blood is of unknown physiological benefit, but this feature is a crucial aspect of tafamidis’ role as an aggregation inhibitor to ameliorate TTR amyloidosis. Negative cooperativity allows a single equivalent of tafamidis to nearly fully occupy one of TTR’s binding sites, instead of yielding a mixture of unbound, single-bound, and double-bound complexes. This is notable, since the binding of one site is sufficient to stabilize the TTR tetramer against dissociation, the rate-limiting step of TTR aggregation^6^.

**Figure 4.**
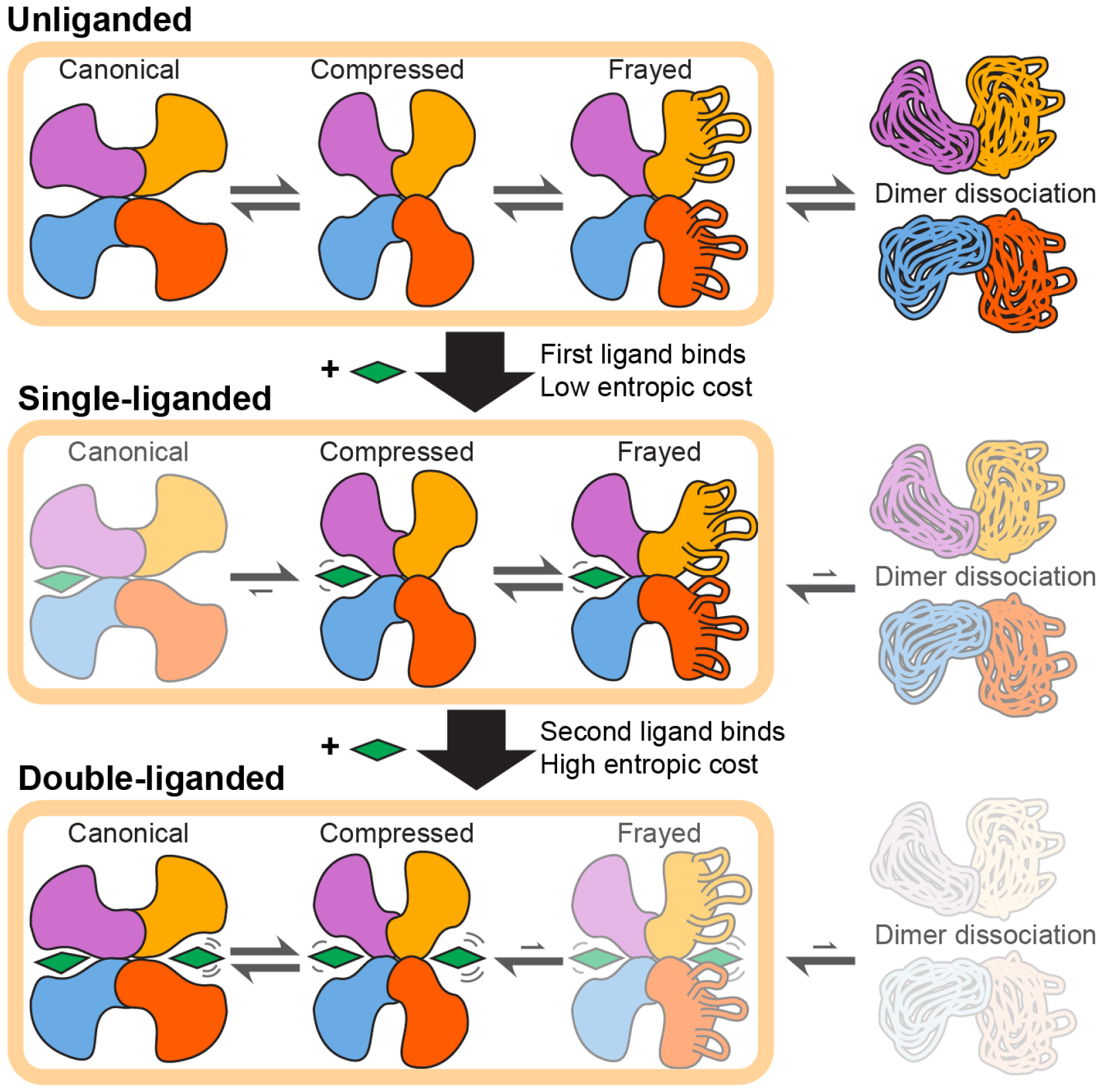
Schematic outlining the conformational landscape of TTR and how it is influenced by ligand binding. A cartoon representation of the TTR tetramer is colored by subunit as in other figures, with the ligand represented as a diamond. In the absence of ligand, the TTR tetramer adopts an equilibrium of conformational states described in this work, with some fraction of the tetramer having a propensity to dissociate into dimers, which could subsequently further dissociate and aggregate. A small molecule ligand can bind one of the two available binding pockets in the TTR tetramer. This binding occurs without incurring a substantial entropic cost, as the ligand binds to the distinct binding pocket that more readily accommodates the interaction. This single ligand binding event is sufficient to increase the stability of the dimer-dimer interactions, lowering the propensity for dimer dissociation and aggregation. The next binding event by the same ligand will incur a much larger entropic cost, as the remaining binding pocket requires structural rearrangements to accommodate the ligand. Furthermore, stabilization of the dimer-dimer interfaces induces an ordering of the loops associated with the frayed state, which comes with an entropic cost.

It is not yet possible to explain why WT TTR dissociates, misfolds, and aggregates in about 10% of the elderly, leading to cardiomyopathy and peripheral neuropathy. We can now explore whether an aging-associated change in the distribution of the canonical, compressed, and frayed states, or the population of a new TTR state, could render a sub-population of WT TTR amyloidogenic.

Our findings highlight the role of the inherent asymmetry in the quaternary structure of TTR and the asymmetric rearrangements occurring upon ligand binding to the second T4 binding site (properties either averaged or suppressed in prior crystallographic studies) to explain the basis of TTR’s negatively cooperative ligand binding. We present previously unobserved TTR tetramer conformations, including a frayed state that could be prone to aberrant proteolysis and aggregation, particularly if the population of this state increased upon aging, for example due to proteostasis network deficiencies. Our observations suggest that proteins exhibiting apparently symmetrical quaternary structures could benefit from being reexamined by single-particle cryo-EM to discern whether less-populated or asymmetric states exist natively. Given that transthyretin was used to exemplify structure-based drug design, the potential of single-particle cryo-EM studies to upend our understanding of structural biology should be considered, even for proteins that have been historically considered too small to visualize using these methods or that have been studied for decades and where fundamentally new insights were not expected.

## Materials and Methods

### Fabrication of graphene grids for cryo-EM

#### Batch preparation of gold-substrate grids for coating with graphene

We prepared graphene grids using a previously established protocol^16^. 12 to 24 R1.2/1.3 or R0.6/1.0 UltrAuFoil holey gold grids were washed by submersion in *∼*100 mL of chloroform while slowly stirring on an orbital shaker for 30 minutes. This washing step was carried out in a crystallizing dish covered with tin foil, with grids placed at the bottom, on a PELCO Clamping TEM aluminum grid holder block (prod. 16830-45) without the lid on. Two more washes were carried out in the same manner, first with acetone and subsequently with isopropanol. Between washes, grids were quickly transferred by initially placing the holder lid on the submerged lower half of the holder and scooping the assembly out with a metal fork bent to form an upright scoop, and then submerging the assembly into the next organic solvent bath at an angle of *∼*45^*°*^ to avoid bubbles forming at the bottom. Finally, the lid was removed by pulling it up by the central holes with an 8-inch stainless steel tweezers with curved pointed serrated tip. After the final isopropanol bath, grids were picked up one by one and left to dry on air on filter paper for 15 minutes. Grids were then immediately coated with monolayer graphene.

#### Preparation of ∼1.5 x 1.5 cm graphene film for coating of gold-substrate grids

A 15x15 cm CVD graphene film on Cu was purchased from Graphenea. In order the transfer a single layer of graphene to gold-support TEM grids, the purchased film was coated with methyl methacrylate (MMA, “MMA EL 6” from Kayaku Advanced Materials, M310006 0500L1GL) using a homemade spin coater, as described previously^8^. The large CVD film on Cu was cut in small squares that fit the spin coater glass slide (*∼*1.5x1.5 cm). Before MMA coating, filter paper was placed between the bottom of the Cu-graphene stack and the spin coater glass slide to prevent MMA from seeping under and coating the Cu from the bottom. Once coated, the MMA-graphene-Cu stack was placed Cu-side up on a glass slide and plasma cleaned for 20 seconds at 25 Watts (75% Argon/25% oxygen atmosphere) using a Solarus plasma cleaner (Gatan, Inc.). To etch the Cu layer, the MMA-coated and plasma-cleaned stack was floated MMA-side up on 200 mL of freshly made 200mM ammonium persulfate in a crystallizing dish for 3 hr. For rinsing, the graphene-MMA film was carefully picked up with a glass slide and transferred to a new crystallization dish with 200 mL of MilliQ water for 1 hour. Immediately after that, the film was used to coated grids. The leftover uncoated graphene was stored in the dark and under vacuum.

#### Batch coating of gold-substrate grids with monolayer graphene

Right after cleaning, the grids were submerged in water inside a grid coating trough (LADD Research Industries, Smith Grid Coating Trough 10840), on a small filter paper square that was cut to fit the metal tray of the trough. Grids were placed very closely to one another, forming as tight a square array as possible. The rinsed MMA-graphene film was then floated onto the trough’s water surface. Using a pair of glass Pasteur pipettes whose tips had been closed by melting, we aligned the MMA-graphene film and grids. By extracting water with the syringe attached to the trough and guiding the MMA-graphene film as the water level went down, we coated 12-24 grids at the same time. The coated grids were left to dry for 30 minutes, and then further dried at 65 ^*°*^C for another 30 minutes. After letting the grids cool down at room temperature for 5 minutes, we proceeded to remove the MMA layer by rinsing grids in a series of organic solvents, using the same setup as for the initial grid cleaning. We used two baths of acetone for 30 minutes each, then a final isopropanol bath for 30 minutes. Grids were left to dry at room temperature on filter paper for 10 minutes, and then stored in dark under vacuum until used. Grids were used up to one month later without noticeable issues.

#### UV-ozone treatment of monolayer graphene grids

To make monolayer graphene able to take aqueous sample, we treated the graphene grids with UV/ozone using the UVOCS T10x10 system, as described previously^8,16^. To achieve consistent performance, we followed the manufacturers’ instructions and carried out a 10-minute “warmup” run, before immediately inserting and treating grids for 4 minutes.

### Expression and purification of WT, C10A, and FT_2_-WTTetrameric TTR

#### Expression of WT tetrameric TTR

BL21 (DE3) competent *E. coli* (New England BioLabs, Ref #C2527H) were transformed with the pMMHa vector encoding the WT TTR gene. The transformed cells were selected by growing them on agar plates containing 100 μg/mL ampicillin at 37 ^*°*^C. *E. coli* starter cultures were initiated by taking a single colony from the agar plate and culturing it in 50 mL LB media in a 250 mL shake flask containing 100 μg/mL ampicillin at 37 ^*°*^C. These cultures were grown at 37 ^*°*^C with shaking until bacterial growth indicated by turbidity in the media were observed. The expression cultures were prepared by inoculating 1 L shake flask containing 100 μg/mL ampicillin with a 1:20 dilution of the starter cultures. The expression cultures were grown at 37 ^*°*^C until OD_600_ were approximately 0.5, at which point they were induced with 1mM IPTG and incubated overnight at 30 ^*°*^C with shaking. These cultures were then harvested by centrifugation at 6,000 rpm for 30 minutes at 4 ^*°*^C. The supernatants were removed, and the pellets were stored at –80 ° C until ready for subsequent purification steps.

#### Expression of C10A tetrameric TTR

BL21 (DE3) competent *E. coli* (New England BioLabs, Ref #C2527H) were transformed with the pMMHa vector encoding the C10A TTR gene. The transformed cells were selected by growing them on agar plates containing 100 μg/mL ampicillin at 37 ^*°*^C. *E. coli* starter cultures were initiated by taking a single colony from the agar plate and culturing it in 50 mL LB media in a 250 mL shake flask containing 100 μg/mL ampicillin at 37 ^*°*^C. These cultures were grown at 37 ^*°*^C with shaking until bacterial growth indicated by turbidity in the media were observed. The expression cultures were prepared by inoculating 1 L shake flask containing 100 μg/mL ampicillin with a 1:20 dilution of the starter cultures. The expression cultures were grown at 37 ^*°*^C until OD_600_ were approximately 0.5, at which point they were induced with 1mM IPTG and incubated overnight at 30 ^*°*^C with shaking. These cultures were then harvested by centrifugation at 6,000 rpm for 30 minutes at 4 ^*°*^C. The supernatants were removed, and the pellets were stored at –80 ° C until ready for subsequent purification steps.

#### Expression of FT_2_-WT tetrameric TTR

BL21 (DE3) competent *E. coli* (New England BioLabs, Ref #C2527H) were transformed with the pET-29a vector encoding the N-terminal dual-FLAG-tagged WT TTR gene. The transformed cells were selected by growing them on agar plates containing 50 μg/mL kanamycin at 37 ^*°*^C. *E. coli* starter cultures were initiated by taking a single colony from the agar plate and culturing it in 50 mL LB media in a 250 mL shake flask containing 50 μg/mL kanamycin. These cultures were grown at 37 ^*°*^C with shaking until bacterial growth indicated by turbidity in the media were observed.

The expression cultures were prepared by inoculating a 1 L shake flask containing 50 μg/mL kanamycin with a 1:20 dilution of the starter cultures. The expression cultures were grown at 37 ^*°*^C until OD_600_ were approximately 0.5, at which point they were induced with 1mM IPTG and incubated overnight at 30 ^*°*^C with shaking. These cultures were then harvested by centrifugation at 6,000 rpm for 30 minutes at 4 ^*°*^C. The supernatants were removed, and the pellets were stored at –80 ^*°*^C until ready for subsequent purification steps.

#### Purification of tetrameric TTR

The pellets were thawed and resuspended with 50 mL Tris-Buffered Saline (TBS) with an added protease inhibitor tablet (Thermo Scientific Pierce Protease Inhibitor Tablets EDTA-Free, Ref #A32965), followed by 3 cycles of probe sonication (Qsonica Q125; 3 minutes sonication / 3 minutes rest at 4 ° C). The supernatants were collected with centrifugation at 15,000 rpm for 30 minutes, subjected to 50% ammonium sulfate precipitation (w/v), and stirred for 45 minutes at room temperature; then, supernatant was collected via centrifugation at 15,000 rpm for 30 minutes at 4 ^*°*^C. The supernatants were further precipitated with 90% ammonium sulfate (w/v) and stirred for 45 minutes at 25 ^*°*^C (room temperature). The solutions were then centrifuged, and the pellets were collected and dialyzed with 3,500 MWCO dialysis tubing (SnakeSkin Dialysis Tubing, Ref #68035) overnight against 25mM Tris pH 8.0 buffer in a 4 ^*°*^C cold room.

The protein mixtures were filtered using a low protein binding filter (Millipore Sigma) and purified on a SourceQ15 anion exchange column (Cytiva) at a flow rate of 2 mL/min. Protein mixtures were injected at 5% Buffer A (25mM Tris pH 8.0) and eluted over a 60-minute gradient up to 100% Buffer B (25mM Tris/1.0 M NaCl) at room temperature. The eluates were then subjected to further purification step using a Superdex 75 gel filtration column. TTR proteins were eluted with 10mM sodium phosphate pH 7.6 / 100mM KCl, confirmed for purity and mass by LC-MS, aliquoted, and stored in –80 ^*°*^C.

### Subunit exchange to generate (FT_2_-WT)_1_(C10A)_3_ TTR tetramer

Subunit exchange between recombinant C10A TTR (40μM) and recombinant dual-FLAG-tagged WT (20 μM) was carried out by adding stocks of each TTR variant to the same buffer sample and incubating the sample for two days. Subsequently, the sample was injected into a 4.6 x 100 mm Waters Protein-Pak Hi-Res Q ion exchange column (5 μm particle size) using Buffer A (25 mM Tris pH 8.0) as mobile phase in Waters UPLC system. The sample was eluted using a linear gradient 24-39% Buffer B (25 mM Tris pH 8.0/1.0 M NaCl) with a flow rate of 0.5 mL/min, in which separation of the five peaks resulting from subunit exchange was observed. The second peak, which corresponds to (FT_2_-WT)_1_(C10A)_3_ TTR, was isolated and confirmed for purity via HPLC. The concentration of (FT_2_-WT)_1_(C10A)_3_ TTR was determined using NanoDrop based on the molar absorptivity (ε = 75829 M^−1^ cm^−1^). The purified heterotetramer was stored at –80 ^*°*^C.

### Modification of (FT_2_-WT)_1_(C10A)_3_ TTR tetramer with *Biarylamine 7*

Previously purified (FT_2_-WT)_1_(C10A)_3_ (40μM) was reacted with a 10-fold molar excess of Biarylamine 7 for 2 hours at room temperature (25 ^*°*^C) with occasional vortexing. The samples were then purified using a Microcon 3,000 MWCO centrifugal filter column (Milipore Sigma) spun at 12,000 rpm (4 ^*°*^C). The identity of the conjugate eluting ((biarylamine-FT_2_-WT)_1_(C10A)_3_ TTR, also referred to as single-bound TTR) was then confirmed using a liquid chromatography coupled to Time-of-Flight mass spectrometry (Agilent LC/MS TOF) employing an Agilent Zorbax 300SB-C8 4.6x50mm column, confirming that only the dual-flag-tagged WT TTR peak was modified by Biarylamine 7. (C10A TTR = 13860 Da, FT_2_-WT TTR = 15882, FT_2_-WT + Biarylamine 7 minus the 2-thiopyridine leaving group = 16325)

### Synthesis of small molecule covalent modifier 2-((3,5-dichloro-4-(2-(2-(2-(pyridin-2-yldisulfaneyl)ethoxy)ethoxy)ethoxy)phenyl)amino) benzoate “Biarylamine 7”

Synthesis of the covalent small molecule 7 was adapted from the synthetic route previously published^6^.

#### General synthetic procedures

All starting materials and solvents were purchased commercially from Sigma-Aldrich, Acros, Alfa Aesar, Combi-blocks, and EMD Millipore and were used without further purification. Thin-layer chromatography was carried out using Merck silica plates (60-F254), using UV light for visualization to monitor reaction progress. Liquid chromatography mass spectrometry (LC-MS) was performed using a ZORBAX RRHT StableBond C18, 2.1 x 50 mm, 1.8 μm column (Agilent Infinity, mobile phase A = 0.1% Formic Acid in H_2_O, mobile phase B = 0.1% Formic Acid in MeCN) coupled to an Agilent G6125B single quadrupole mass spectrometer (Agilent Infinity) was used to monitor reaction progress. Flash column chromatography was carried out using a Teledyne Isco Combiflash Nextgen 300+ machine using Luknova SuperSep columns (SiO2,25 μm) with ethyl acetate and hexanes employed as eluents. ^1^H NMR spectra were recorded on Bruker 400 or 500 MHz spectrometer. Characterization data are reported as follows: chemical shift, multiplicity (s=singlet, d=doublet, t=triplet, q=quartet, br=broad, m=multiplet), coupling constants, number of protons.

#### 4-Benzyloxy-3,5-dichloroaniline (Compound 3 in **Supplementary Fig. 22**)

Di-tert-butyl dicarbonate (11 mmol, 2.4 g, 1.1 eq) was added dropwise to a stirring solution of 4-amino-2,6-dichlorophenol (10 mmol, 1.96 g, 1.0 eq) in 50 mL THF. The resulting solution was refluxed for 12 hours before concentration in vacuo. The crude solid was dissolved in DMF before addition of potassium carbonate (20 mmol, 2.76 g, 2.0 eq) and tetra-n-butylammonium iodide (1 mmol, 369 mg, 0.1 eq). Benzyl bromide (1 mmol, 1.78 mL, 1.5 eq) was added dropwise to the stirring solution and stirred at room temperature for 1 hour. Upon reaction completion as discerned by TLC, the solution was diluted with NH_4_Cl and water and extracted with EtOAc (3 x 100 mL). The combined organics were concentrated, and in the same flask, 30% TFA in DCM was added dropwise. After stirring for 1 hour, the solution was neutralized with NaCO_3_ and saturated NaHCO_3_, extracted with EtOAc (3 x 50 mL), and concentrated. Flash column chromatography purification (SiO_2_, 4:1 Hexanes:EtAOc) afforded 4-Benzyloxy-3,5-dichloroaniline as a white powder (1.1 g, 41% yield)^1^H NMR (500 MHz, CDCl_3_) δ 7.58 (d, *J* = 7.4 Hz, 2H), 7.42 (t, *J* = 7.4 Hz, 2H), 7.38 (d, *J* = 7.3 Hz, 1H), 6.66 (s, 2H), 4.97 (s, 2H), 3.56 (s, 2H) (**Supplementary Fig. 23**).

#### tert-Butyl 2-((4-(benzyloxy)-3,5-dichlorophenyl)amino) benzoate (Compound 4 in **Supplementary Fig. 22**)

Compound 3 (0.5 mmol, 134 mg, 1.0 eq), *tert*-butyl 2-bromobenzoate (i) (0.6 mmol, 116 μL,1.2 eq), cesium carbonate (0.7 mmol, 228 mg, 1.4 eq), RuPhos Pd G3 (0.05 mmol, 41.8 mg, 0.1 eq) were dissolved in toluene under argon. After refluxing under inert atmosphere for 24 hours, the reaction was filtered through celite and concentrated in vacuo. Flash column chromatography purification (SiO_2_, 95:5 Hexanes:EtAOc) afforded *tert*-Butyl 2-((4-(benzyloxy)-3,5-dichlorophenyl)amino)benzoate as a yellow solid (96 mg, 43%).^1^H NMR (500 MHz, CDCl_3_) δ 9.57 (s, 1H), 7.93 (dd, *J* = 8.0, 1.7 Hz, 1H), 7.60 – 7.55 (m, 2H), 7.44 – 7.40 (m, 2H), 7.36 (ddt, *J* = 10.2, 4.9, 1.5 Hz, 3H), 7.24 (dd, *J* = 8.5, 1.1 Hz, 1H), 7.22 (s, 2H), 6.80 (ddd, *J* = 8.1, 7.1, 1.1 Hz, 1H), 5.03 (s, 2H), 1.61 (s, 9H) **(Supplementary Fig. 24)**.

#### tert-Butyl 2-((3,5-dichloro-4-hydroxyphenyl)amino) benzoate (Compound 5 in **Supplementary Fig. 22**)

Compound 4 (0.2 mmol, 90 mg, 1 eq) and 10% palladium on carbon (0.017 mmol, 18 mg, 0.09 eq) and Compound **4** (0.2 mmol, 90 mg, 1 eq) were dissolved in a 1:1 MeOH:EtOAc mixture and stirred under an H_2_ atmosphere balloon for 5 hours. Upon observation of complete consumption of the starting material by TLC, the reaction was diluted with DCM, filtered through celite, and concentrated in vacuo. Flash column chromatography purification (SiO_2_, 5:1 Hexanes:EtOAc) afforded *tert*-butyl 2-((3,5-dichloro-4-hydroxyphenyl)amino)benzoate as a yellow solid (70 mg, 98% yield). The physical and spectral data are consistent with those reported previously^6^.

#### S-(2-(2-(2-hydroxyethoxy)ethoxy)ethyl) ethanethioate (Compound iii in **Supplementary Fig. 22**)

Thioacetic acid (5 mmol, 350 μL, 1 eq) and potassium carbonate (5 mmol, 690 mg, 1 eq) were dissolved in anhydrous DMF (10 mL) and stirred for 10 minutes. To the resulting solution was added 2-(2-(2-bromoethoxy)ethoxy)ethan-1-ol (4.7 mmol,1.065 g, 0.95 eq) dropwise before stirring at 40^*°*^ C for 2 hours. The solvent was removed under reduced pressure and crude residue purified by flash column chromatography (SiO_2_, 1:1 hexanes:EtOAc) to give a clear yellow-clear oil (755 mg, 77% yield).^1^H NMR (400 MHz, CDCl3) δ 3.67 (dd, J = 5.4, 3.8 Hz, 2H), 3.64 – 3.51 (m, 8H), 3.04 (t, J = 6.5 Hz, 2H), 2.28 (s, 3H) (**Supplementary Fig. 25**).

#### tert-butyl 2-((4-(2-(2-(2-(acetylthio)ethoxy)ethoxy)ethoxy)-3,5-dichlorophenyl)amino) benzoate (Compound 6 in **Supplementary Fig. 22**)

Diisopropyl azodicarboxylate (80.8 μL, 0.4 mmol, 2 eq) and triphenylphosphine (104.8 mg, 0.4 mmol, 2 eq) were dissolved in 4 mL anhydrous THF and stirred for 30 minutes at room temperature. After 30 minutes, **5** (70 mg, 0.2 mmol, 1 eq) and **iii** (83.2 mg, 0.4 mmol, 2 eq), each dissolved in 0.2 mL THF, were added dropwise and stirred at room temperature for 2 hours. The reaction was concentrated in vacuo and the crude residue purified by column chromatography (SiO_2_, 4:1 Hex:EtOAc) to afford yellow syrup (35.4 mg, 32.6% yield).^1^H NMR (400 MHz, ACETONE-D6) δ 9.47 (s, 1H), 7.91 (dd, J = 8.0, 1.7 Hz, 1H), 7.41 (ddd, J = 8.7, 7.1, 1.7 Hz, 1H), 7.32 – 7.23 (m, 3H), 6.83 (ddd, J = 8.1, 7.0, 1.1 Hz, 1H), 4.15 (dd, J = 5.6, 4.1 Hz, 2H), 3.86 – 3.80 (m, 2H), 3.68 – 3.61 (m, 2H), 3.61 – 3.49 (m, 4H), 3.02 (t, J = 6.5 Hz, 2H), 2.27 (s, 3H), 1.58 (s, 9H) (**Supplementary Fig. 26**).

#### 2-((3,5-dichloro-4-(2-(2-(2-(pyridin-2-yldisulfaneyl)ethoxy)ethoxy)ethoxy)phenyl)amino)benzoate (Biarylamine 7) (Step 7 in **Supplementary Fig. 22**)

Compound 6 (22.5 mg, 0.0414 mmol, 1 eq), and 2,2’-dithiopyridine (11 mg, 1.2 eq, 0.05 mmol) were dissolved in 1 mL anhydrous THF under an argon atmosphere. To the resulting solution was added freshly prepared NaOMe in THF (110 μL, 1.2 eq, 0.05 mmol) dropwise at 0 ^*°*^C. The reaction was allowed to warm to room temperature over 2 hours before diluting with EtOAc and sequential brine and water washes. The combined organic extracts were removed under reduced pressure before addition of 1 mL TFA. After stirring at room temperature for 2 hours, the solvent was removed under reduced pressure and final compound Biarylamine 7 was purified via preparative HPLC (6.2 mg, 27% yield over two steps).^1^H NMR (500 MHz, DMSO) δ 9.49 (s, 1H), 8.44 (dd, J = 4.8, 1.5 Hz, 1H), 7.92 (dd, J = 8.0, 1.7 Hz, 1H), 7.87 – 7.80 (m, 2H), 7.46 (ddd, J = 8.7, 7.1, 1.7 Hz, 1H), 7.34 (s, 2H), 7.27 – 7.19 (m, 2H), 6.89 (t, J = 7.5 Hz, 1H), 4.12 – 4.07 (m, 2H), 3.81 – 3.75 (m, 2H), 3.67 – 3.58 (m, 4H), 3.51 (dd, J = 5.8, 3.6 Hz, 2H), 3.03 (d, J = 12.1 Hz, 2H) (**Supplementary Fig. 27**).

### Crystallization and structure determination of (Stilbene)_2_-TTR

Crystals of (Stilbene)_2_-TTR were grown analogously as previously described for TTR co-crystals with the covalent kinetic stabilizer 4-fluorophenyl 3-[(*E*)-2-(4-hydroxy-3,5-dimethylphenyl)ethenyl]benzoate (A93, PDB: 3HJ0).^18^ Data were collected at beamline 5.0.1 at the Advanced Light Source (Berkeley, CA) at a wavelength of 0.97741 Å and a temperature of 100 K. Frames were indexed and integrated using XDS^19^, the space group was assigned as P2_1_2_1_2 using Pointless, and data were scaled using Scala.^20^ Five percent of reflections (randomly distributed) were flagged for model cross-validation using R_free_. ^21^ The (Stilbene)_2_-TTR crystal structure was solved by molecular replacement with Phaser^22^, using one monomer of human WT TTR as a search model.

This model was generated from the co-crystal structure of tafamidis-bound WT TTR (PDB code: 3TCT^23^) with the ligands removed. Two subunits (one dimer) were found in the asymmetric unit. A small molecule stabilizer CIF dictionary was generated using ACEDRG^24^. The model was refined with iterative cycles of manual adjustment in Coot^25^ and refinement in Refmac5 using isotropic thermal parameters and hydrogen atoms at calculated positions^26^. Final adjustments were made after analysis with MolProbity and the wwPDB Validation System^27^. Due to the poor electron density surrounding the Lys15-stilbene linkage, the Lys15 sidechain and amide linkage could not be modeled with certainty. Data acquisition and modeling metrics are reported in **Supplementary Table 1**. The crystal structure was deposited to the Protein Data Bank under accession code 8U52.

### Mass spectrometry analysis of the (Stilbene)_2_-TTR conjugate for cryo-EM sample preparation

Before sample freezing, to verify that the major species in the (Stilbene)_2_-TTR sample is conjugated at both TTR binding sites, we carried out liquid chromatography followed by mass spectrometry (LC-MS, with LC equipment Agilent 1260 and mass spectrometer Agilent 6230 ESI-TOF) on a 3X diluted aliquot taken 3 hours after starting the reaction. For this, 5 μL of sample was injected in a reverse phase column (Agilent PLRP-S 2.1 x 50 mm 100A 5 μm) held at 60 ^*°*^C and eluted with a water-to-acetonitrile gradient from 95% to 50% water, and a flow of 0.5 mL/min, 95% for 2 min then 50% water for 6 min.

To probe for the existence of A2 hydrolysis products, we carried out a time series of the A2+TTR reaction (same concentrations as used for sample preparation), injecting sample every 30 minutes for 5 hours. We monitored UV absorbance at 280 and 305 nm, as well as total ion count. We also analyzed samples containing only A2 (150μM A2 in TTR buffer + 10% DMSO) and only TTR (21μM TTR in TTR buffer). The baseline was removed using the ZhangFit method from the BaselineRemoval python package^28^. The exponential labeling temporal curves were generated by normalizing the area under the curve dividing by the maximum, after baseline removal. See results in **Supplementary Fig. 31** and **Supplementary Text 4**.

### Unliganded human transthyretin cryo-EM sample preparation

A 50 μL solution of 1.7 mg/mL TTR in 100 mM KCl, 10 mM Phosphate buffer pH 7.6 (1 mM EDTA) was thawed at room temperature and centrifuged for 10 minutes at 16,000 x g, then diluted to 0.85 mg/mL. To make 20 μL of sample, we added 5 μL of TTR, 5 μL of TTR buffer, and 10 μL of 0.1% w/v beta-octyl glucoside (BOG) in TTR buffer, resulting in a final concentration of 7.31μM TTR and 0.01% w/v BOG. For grid preparation, R1.2/1.3 UltrAuFoil Holey Gold grids covered with monolayer graphene were oxidized using UV/ozone immediately before use. A manual plunger in a cold room at 6 ^*°*^C and 90% humidity was used for flash freezing samples: 3 μL of sample were applied onto the graphene side of the grid after it was mounted on the plunger (always at the same height), and immediately blotted for 6 seconds by holding a *∼*1 x 6 cm piece of Whatman #1 filter paper parallel to the grid, in full contact. Timing was kept with the help of a metronome. The blotting countdown was started after the blotted liquid spot on the filter paper stopped spreading and the grid was plunged in liquid ethane at the same time the blotting paper was pulled back, in a single motion. Four grids were prepared this way, two pairs of identical grids blotted for 4 and 6 seconds, which were later clipped right before the imaging session. Imaging was performed on one of the grids blotted for 4 seconds.

### (Stilbene)_2_-TTR sample preparation for cryo-EM

A 50 μL solution of 1.7 mg/mL TTR in 100 mM KCl, 10 mM phosphate buffer pH 7.6 and 1 mM EDTA was thawed at room temperature and spun down for 10 minutes at 16000 x g. The TTR+A2 reaction was set up by adding 1 μL of a 1.5 mM solution of A2 in DMSO to 19 μL of the TTR stock solution specified immediately above. The A2+TTR reaction was incubated for 4 hours in the dark at room temperature.

To make 20 μL of sample, we added 5 μL of TTR+A2 incubation reaction, 5 μL of TTR buffer, and 10 μL of 0.1% w/v beta-octyl glucoside (BOG) in TTR buffer, resulting in final concentrations: 7.31μM TTR, 18.75μM A2, 0.01% w/v BOG, and 1.25% v/v DMSO. Grids were prepared in the same way as the unliganded TTR conjugate sample, except that sample was blotted for 6 and 8 seconds.

### Single-bound (biarylamine-FT_2_-WT)_1_(C10A)_3_ TTR cryo-EM sample preparation

A frozen (FT_2_-WT)_1_(C10A)_3_ TTR Tetramer sample at 0.9 mg/mL was thawed at room temperature for 10 minutes, and then used to prepare grids. Grids were prepared in the same way as the (Stilbene)_2_-TTR conjugate sample, except that sample was blotted for 3 seconds.

### Cryo-EM data acquisition for single-particle analysis

Movies of all frozen-hydrated TTR samples were collected using a Talos Arctica TEM (Thermo Fisher Scientific) with a field emission gun operating at 200 keV, equipped with a K2 Summit direct electron detector (Gatan, Inc.). Images were acquired using the Leginon automated data collection software^29^. Alignments were performed as previously described to minimize coma and establish parallel illumination^30,31^, and stage movement was used to center image holes in a 4 x 4 array for (Stilbene)_2_-TTR and unliganded TTR samples (graphene-coated R1.2/1.3 UltrAuFoil grids) and 8x8 array (graphene-coated 0.6/1 UltrAuFoil grids) for all other samples. Image shift with beam tilt compensation was used to acquire images within each hole in the centered array. Movies were collected in counting mode (0.562 Å/pixel) with an exposure rate of 2.3 e^-^/pix/s, with 50 frames (150 ms each) over 7.5 s, totaling an exposure of 55 e^-^/Å^2^. A nominal defocus between –1.0 μm and –1.5 μm was used. The Appion image analysis environment^32^ was used to carry out real-time frame alignment and estimation of the contrast transfer function (CTF) to assess quality of ongoing data acquisition.

### Data analysis of unliganded TTR

Frames from 4,634 movies were aligned and combined applying a dose-weighting scheme using MotionCor2^33^ from within the RELION 3.1 interface^34^. The aligned micrographs were imported to cryoSPARC 3.1.0^35^ and CTF was estimated using CTFFind4^36^, from within the cryoSPARC interface. Micrographs with estimated defocus values higher than 1.5 μm and reported CTF resolutions lower than 4 Å were discarded. Micrographs displaying no particles and torn or crumpled and stacked graphene were also discarded, resulting in 3,739 micrographs. For template-based particle picking, a density model of PDB 1TTR chains A, B, A’ and B’ (full native homotetramer) was generated using UCSF Chimera^37^ and low-pass filtered to 20 Å resolution in cryoSPARC. A maximum of 2000 picks was allowed per micrograph. Picks were inspected and curated using a normalized cross correlation (NCC) and local power (LP) thresholds. Based on visual inspection, using an NCC = *∼*0.5 provided the most accurate particle selection results, and lowering the local power threshold diminished the number of false positive picks on crumpled graphene areas. 1.95 million particle picks were extracted in 256-pixel boxes, and 2D averages were obtained using the 2D classification node in cryoSPARC, changing default settings to 120 classes, 1000 particles per class, 80 iterations and turning off “force Max over poses/shifts,” an inner mask radius of 75 Å, and outer mask radius of 90 Å. Using a lower number of particles per class or keeping “force Max over poses/shifts” selected resulted in a much lower number of different 2D classes with secondary structure features. We were not particularly stringent at the 2D cleanup stage, opting to maintain all classes remotely resembling TTR, which corresponded to 1.49 million particles. We then carried out a second round of 2D classification with the same parameters, which resulted in most classes being identifiable as TTR and with secondary structure features (**Supplementary Fig. 2**). The particles from these classes (1.3 million) were pooled and exported to RELION 3.1 using scripts from the pyem Github repository^38^.

In RELION 3.1, duplicates were removed and the particles re-extracted with a binning factor of 2. The rest of the workflow was carried out in RELION 3.2 (**Supplementary Fig. 3**). We performed a 3D classification imposing C2 symmetry, with the symmetry axis going through the TTR binding pockets, with K=6, tau=4 for 100 iterations. From the six resulting classes, four contained features clearly resembling secondary structure elements. We pooled these classes (870,899 particles) and carried out further 3D classification with C2 symmetry, K=4, tau=4, and 52 iterations. From the resulting four distinct classes, three had well-defined secondary structure elements, which we subsequently 3D refined separately. We used class 1 particles (335,266 particles) as input for 3D auto-refine without imposing symmetry and obtained a 3.5 Å resolution reconstruction. Using K-means clustering, we grouped micrographs by image shift using associated values stored in the Leginon database and assigned them to distinct optic groups. CTF refinement of per-particle, per-micrograph astigmatism, and per-optic group beam tilt using the K-means clustering and reported CTF resolutions lower than 4 μwere discarded. Micrographs displaying no particles and torn or crumpled and stacked graphene were also discarded. For template-based particle picking, a density model of PDB 1TTR chains A, B, A’ and B’ (full native homotetramer) made generated using UCSF Chimera^37^ and low-pass filtered to 20 Å resolution of image shifts resulted in a 3.5 Å resolution reconstruction. Re-extraction of particles without binning, Bayesian polishing (model parameters s vel=1.2855, s div=4755, s acc=2.175, used for all polishing jobs in this dataset) and further CTF refinement improved resolution to 3.3 Å. Based on atomic structure modeling and comparison to previously determined TTR structures in the Protein Data Bank, we termed class 1 as “canonical,” based on its overall similarity to the most common TTR structures determined by crystallography (**Fig. 1 and Supplementary Figs. 4-5**). Class 3 and 4 particles represented similarly compacted conformers of TTR relative to the canonical conformation, although Class 4 particles appeared to have better overall structural integrity, particularly at the periphery of the loops. We used class 4 particles (130,181 particles) as input for 3D auto-refine without imposing symmetry and obtained a 3.5 Å resolution reconstruction. CTF refinement of per-particle, per-micrograph astigmatism and per-optic group (see image shift clustering above) beam tilt resulted in a 3.5 μresolution reconstruction. Re-extraction of particles without binning, Bayesian polishing and further CTF refinement improved resolution to 3.3 Å. We refer to this reconstruction as representative of a “compressed” TTR structural state (**Fig. 1 and Supplementary Fig. 4**). We finally further analyzed the class 3 particles (202,700 particles) with 3D auto-refine without imposing symmetry, and obtained a 3.5 Å resolution reconstruction. CTF refinement of per-particle, per-micrograph astigmatism and per-optic group (see image shift clustering above) beam tilt resulted in a 3.5 Å in cryoSPARC. A maximum of 2000 picks was allowed per micrograph. Picks were inspected and curated using a normalized cross correlation (NCC) and local power (LP) thresholds. Based on visual inspection, using an NCC = *∼*0.5 provided the most accurate particle selection results, and lowering the local power threshold diminished the number of false positive picks on crumpled graphene areas. Picks were extracted in 256-pixel boxes, and 2D averages were obtained using the 2D classification node in cryoSPARC, changing default settings to 120 classes, 1000 particles per class, 80 iterations and turning off “force Max over poses/shifts,” an inner mask radius of 75 Å, and outer mask radius of 90 Å . Using a lower number of particles per class or keeping “force Max over poses/shifts” selected resulted in a much lower number of different 2D classes with secondary structure features. We were not particularly stringent at the 2D cleanup stage, opting to maintain all classes remotely resembling TTR, which contained approximately half of the particles. We then carried out a second round of 2D classification with the same parameters, which resulted in most classes being identifiable as TTR and with secondary structure features (**Supplementary Fig. 10**). The particles from these classes (1.03 million) were pooled and exported to RELION 3.1 using scripts from the pyem Github repository^38^.

Duplicate particles were removed, remaining particles were re-centered and extracted to the same box size using RELION 3.1. The rest of the workflow was carried out in RELION 3.2 (**Supplementary Fig. 11**). Initial 3D auto-resolution reconstruction. Re-extraction of particles without refinement without imposing symmetry led to a 4 Å reconbinning, Bayesian polishing and further CTF refinement improved resolution to 3.3 Å . While this subset of particles were similar in conformation to the compressed Class 4 reconstruction, the disordering of the peripheral loops led us to name this conformation as “frayed” (**Fig. 1 and Supplementary Fig. 4**). The three resulting maps were sharpened by applying an overall B-factor being roughly *∼*1/2 of the sharpening B-factor suggested by RELION 3.1 post-processing node, to avoid excessive noise in the form of “dust” in the final density. The B-factor applied to the canonical, compressed, and frayed reconstructions was –60, –55, and –55, respectively. See **Supplementary Table 2** for further details.

### Cryo-EM data analysis of (Stilbene)_2_-TTR conjugate

Frames from 4,536 movies were aligned and combined applying a dose-weighting scheme using MotionCor2^33^ from within the RELION 3.1 interface^34^. The aligned micrographs were imported to cryoSPARC 3.1.0^35^ and CTF was estimated using CTFFind4^36^, from within the cryoSPARC interface. Micrographs with estimated defocus values higher than 1.5 Å m struction according to a Fourier Shell Correlation (FSC) cut-off at 0.143. CTF refinement of per-particle defocus, per-micrograph astigmatism, and per-group beam tilt using K-means clustering of image shifts (see unliganded TTR image processing methods) led to a reconstruction at 3.4 Å resolution. Further CTF refinement including third-order aberrations improved resolution to 3.1 Å, and Bayesian polishing (model parameters: s vel=0.96, s div=10020, s acc=5.88) improved the resolution further to 2.9 Å . While the preceding steps improved resolution, the resulting density seemed “stretched” or “smeared” in one direction, indicating anisotropic resolution. To solve this, we attempted several 3D-classification schemes, hoping to identify a subset of particles that might produce a more isotropic-resolution reconstruction. 3D classification with 6 classes, a tau value of 4, a circular mask of 90 Å, and no limitation in the E-step resolution resulted in two seemingly isotropic classes after 86 iterations (containing a total of 289,981 particles). Particles from these classes were combined, and 3D auto-refinement led to a 2.7 Å resolution reconstruction where a single stilbene rotamer could be identfied in the pocket formed by chains A and C. We categorized this reconstruction as having a canonical conformation due to its similarity to our unliganded TTR canonical structure (**Fig. 2, Supplementary Figs. 5 and 12**). All other classes resembling TTR (736,525 particles) were pooled and used for 3D classification with 6 classes, tau value=4, and enforcing C2 symmetry with the symmetry axis (Z axis) going through the binding sites. From this classification we were able to isolate 2 more states via 3D auto-refinement without imposed symmetry, one of which was a lower resolution version of the canonical reconstruction (133,817 particles), while the other (105,961 particles, 3.1 Å resolution) we categorized as compressed based on its similarity to the unliganded compressed conformer (**Fig. 2 and Supplementary Fig. 12**). The two resulting maps were sharpened by applying an overall B-factor being roughly *∼*1/2 of the sharpening B-factor suggested by RELION 3.1 post-processing node, to avoid excessive noise in the form of “dust” in the final density. The B-factor applied to the canonical and compressed reconstructions was –60 in both cases. See **Supplementary Table 3** for further details.

### Data analysis of single-bound (biarylamine-FT_2_-WT)_1_(C10A)_3_ TTR

Two datasets consisting of 6,017 and 6,088 movies were acquired using two different grids prepared in the same freezing session and processed separately using the same processing strategies as for the prior two datasets, resulting in 4,002 / 3,937 micrographs yielding 1.23 / 1.42 million particle picks, and 1.17 / 1.26 million particles after 2D cleanup (**Supplementary Fig. 17**). The particles were combined for 3D processing in RELION 3.1 (**Supplementary Fig. 18**), where duplicates were removed and particles re-extracted with a binning factor of 2. 3D classification into 6 classes was performed with tau=4 for 58 iterations imposing C2 symmetry (symmetry axis going through the binding sites). From the six resulting classes, four had discernible secondary structure elements. We refined (3D auto-refine) each of them separately, but only class 3 (233,376 particles, corresponding to the compressed state) and class 5 (432,655 particles corresponding to the frayed state) had isotropic directional resolution, and yielded ligand density in one binding site (**Fig. 3, Supplementary Fig. 18**).

3D auto-refinement of class 3 imposing C2 symmetry yielded a 4 Å reconstruction. CTF refinement of per-particle defocus, per-micrograph astigmatism, and per-optic group beam tilt (see image shift clustering above) led to an improved reconstruction at 3.7 Å resolution. Re-extraction without binning and further CTF refinement improved resolution to 3.6 Å (C2 imposed). Bayesian polishing improved resolution to 3.5 Å (C2 imposed). Refinement of per-particle defocus, per-micrograph astigmatism, beam tilt, and third and fourth order aberrations, as well as 3D auto-refinement without imposing symmetry yielded a reconstruction at 3.5 Å resolution. How-ever, this reconstruction did not contain clear ligand density in either of the two binding sites. To separate different states with a ligand in each binding site, we created two spherical masks encompassing one binding site each and subtracted the signal outside of it from each of them in each particle, using the “Particle Subtraction” node in RELION 3.1. From this process we obtained two new particle stacks in which only the signal from one of the binding sites remained. We carried out 3D classification without alignment with two classes and a tau value of 2 for 100 iterations and obtained two classes for each of them. One of those classification runs yielded a class with apparent ligand density (class 1, 168,443 particles) and one without (class 2, 64,933 particles). 3D auto-refinement without imposed symmetry of these classes after reconstitution of subtracted signal yielded a 3.4 Å reconstruction with clear ligand density in the binding site (168,443 particles from class 1, (biarylamine-FT_2_-WT)_1_(C10A)_3_ TTR compressed state, **Supplementary Table 4**) and one at 3.8 Å without any density in the pocket (64,933 particles from class 2) (**Fig. 3, Supplementary Figs. 18 and 19**).

In contrast to class 3, class 5 displayed a small bi-lobed body of density in one of the binding pockets, which we attributed to the biarylamine moiety, even before refinement. Since per-optic group beam tilt was already calculated for class 3, they were imparted on class 5 particles before carrying out any further analysis. 3D auto-refinement did not improve the resolution or interpretability of class 5, so 3D classification was performed with 6 classes, tau=4, and 70 iterations, which yielded a class (class 6, 110,253 particles) that when refined with 3D auto-refinement imposing C2 symmetry, yielded a 4.1 Å reconstruction. 3D auto-refinement with C2 symmetry after re-extraction without binning, CTF refinement (per-particle defocus, per-micrograph astigmatism and per-optic group beam tilt) and Bayesian polishing yielded a reconstruction at 3.9 Å . Further CTF refinement (per-particle defocus, per-micrograph astigmatism, and per-optic group beam tilt as well as third and fourth-order aberrations) yielded a 4.1 Å reconstruction after 3D auto-refinement without imposing symmetry (biarylamine-FT_2_-WT)_1_(C10A)_3_ TTR frayed state **Supplementary Figs. 18 and 19, Supplementary Table 4**).

### Calculation of TTR H-strand dihedral angles and elongation distance

The dihedral angle between H strands of monomers on opposing sides of a binding pocket is defined by the Cα atoms of residues 114A^→^17A^→^117C^→^114C and 114B^→^117B^→^117D^→^114D. The overall TTR extension lengths, as labeled in the cartoons in figures 1-3, are the averages of the distances between Cα atoms pairs from Leu58 in chains A and B and chains C and D.

### Calculation of Cα dihedral changes between conformations

The change in Cα dihedral angles, as initially presented in^39^, is used to describe hinge-like molecular motions. The Cα dihedral angle at a given position X is calculated from the position of the Cα of positions X+2, X+1, X and X–1. This angle is defined in the –180^*°*^ to 180^*°*^ space, and changes are expressed in absolute values to facilitate visualization.

### Modeling of atomic coordinates in cryo-EM-derived density for all analyzed samples

All atomic coordinate modeling used to create models for cryo-EM datasets presented in this work followed a similar procedure as described below. We began with a model from human wild-type transthyretin, PDB 2QGB. Symmetry expansion was applied in pymol^40^ to generate a tetramer, and chains A’ and B’ were changed to C and D respectively. Non-protein atoms and alternative rotamers were removed. The initial model was docked and aligned to density in UCSF Chimera. The saved and translated copy of the starting model was docked in the density using the “Dock in map” node in Phenix^41^. The resulting model was refined in real space using the “Real-space refinement” node in Phenix, with options “minimization global”, “rigid body”, “local grid search” and “adp” turned on. Secondary structure or initial model restraints were not used. The resulting model was inspected in Coot^42^, where side-chain atoms without density were deleted, and “real-space refine zone” was applied to 8-10 overlapping residue stretches throughout the whole protein. Attempts to remove Ramachandran and side-chain geometry outliers were made by real-space and manual refinement, as well as peptide bond flipping. When considered adequate by local resolution, water molecules were modeled considering their location and likely coordination, as well as their modeling in other TTR structures. For data derived from the (Stilbene)_2_-TTR sample, lysine 15 covalently modified with the stilbene was modeled as a non-canonical amino-acid with the three-letter code A2K. Parameters for A2K were generated using eLBOW and the Lys-stilbene amide bond was modeled in cis as suggested by the density in the highest resolution binding site (chains A and C of the canonical state density). For data derived from the (biarylamine-FT_2_-WT)_1_(C10A)_3_ TTR sample, parameter files and models for the small molecule moiety were obtained from the Protein Data Bank, as crystal structures of this complex have already been published^6^. In (biarylamine-FT_2_-WT)_1_(C10A)_3_ TTR position B10 was modeled as cysteine, while A10, C10 and D10 were modeled as alanine. This decision was based on the local density features in the compressed state, given its higher resolution.

In all models, after the initial manual refinement in coot the model was refined again using the “Real-space refinement” node in Phenix^41^. This process was repeated until no changes were considered by manually inspecting the model in Coot; i.e., “Real-space refinement” in Phenix was the last step in all modeling. Data modeling statistics produced with Phenix^41^ are reported in **Supplementary Tables 2-4**. Map-to-model Fourier correlation plots are presented in **Supplementary Figs. 4, 12, and 19**.

### Figure generation

The DeepEMHancer^43^ software package was used to sharpen the density maps for generating figure panels depicting the TTR reconstructions. These DeepEMHancer-generated maps were not used for modeling or interpretation. UCSF ChimeraX^44^ was used to create the surface representations of cryo-EM densities and ribbon/licorice representations of atomic models. UCSF Chimera^37^ was used to generate the wire-mesh representations that include the atomic models.

### Ligand-binding pocket volume measurement with Fpocket

Measuring pocket volume in a consistent and comparable way can be challenging, as automatic detection may produce very different results in similar structures depending on the area accessible to a solvent-like probe, methods for defining the limits of the pocket, and slight structural differences such as sidechain conformation. For these reasons, we decided to use the Fpocket functionality^45^ that defines pockets by the presence of a ligand, and modeled missing sidechains in our seven TTR structures using Rosetta^46^ (see below). To account for ambiguities in the density, we generated 10 models per cryo-EM density (70 in total), which enabled us to estimate the error associated with the measurements (**Supplementary Fig. 32**). To minimize noise coming from ligand placement and chemistry, and to enable ligand-based pocket volume measurement in unliganded TTR, we took the canonical (Stilbene)_2_-TTR conjugate structure, aligned it to each of the other six species, and grafted the atomic coordinates of the stilbene moiety in each of them. We then ran Fpocket in the newly generated artificial (Stilbene)_2_-TTR complexes.

### Full-atom model refinement with Rosetta

To obtain reasonable estimates of pocket volume, we used Rosetta to generate full-atom models from the partial (missing side chains) models made in Coot. We followed the protocol described in^47^. Briefly: rotamer positions were optimized by Monte Carlo optimization of the Rosetta score function^47^ and density values, followed by gradient minimization of all atom positions, allowing covalent interactions to deviate lightly from their theoretical optima. These steps were carried for four iterations going from low-repulsion (”soft ball” atoms) to full-repulsion van der Waals potential, in a routine called “FastRelax”. The necessary scripts and command-line options are provided in the supplementary materials. Ten structures were produced for each initial Coot model to account for uncertainty derived from the lack of density. It should be emphasized that, given the conservative nature of the FastRelax protocol, the resulting full atom models only differ by small backbone and side-chain motions.

### MD Simulations of the (Stilbene)_2_-TTR Conjugate

TTR structures with covalent stilbene were prepared with pdb4amber within AmberTools21^48^, including residue protonation-state determination with REDUCE^49^. Structurally determined water molecules were also removed. Antechamber^50^ was used to calculate AM1-BCC partial-charges for the ligand-residue adduct via the SQM method. Lysine 15 with attached stilbene ligand was given N-methyl and C-acetyl caps prior to Antechamber calculations. The ff19SB^51^ forcefield was used for protein residues, while GAFF2^52^ was used for the stilbene ligand. To maintain planarity of the amide linkage between the K15 residue and the stilbene ligand, the N_z_ was explicitly parameterized with the GAFF2 sp^2^ amide N atom type (”n”) instead of the N3 ff19SB atom type. An OPC^53^ water box was placed around the protein with at least a 15 Å buffer; then Na^+^ and Cl^-^ ions were added to neutralize the system. Minimization and simulation were performed with OpenMM v7.7.0^54^. Following an energy minimization, the simulation was equilibrated for 0.1 nanoseconds to prior to the production run of 500 nanoseconds. The simulation was run with an NPT ensemble at 310 K and 1.0 ATM with periodic boundary conditions. A Monte Carlo barostat was used with an interval of 25 steps along with a Langevin Middle integrator/thermostat with friction coefficient of 1.0/ps. A 2 fs timestep was used. The Particle Mesh Ewald method^55^ was used for electrostatic interactions with a nonbonded cutoff of 1.2 nm and a tolerance of 0.0005. The length of bonds involving hydrogen atoms were constrained and water molecules were kept rigid with a constraint tolerance of 0.000001. The resulting trajectory was autoimaged with CPPTRAJ^56^ and analyzed with MDAnalysis in Python^57^.

## Acknowledgements

We thank Bill Anderson and Charles Bowman at the Scripps Research Electron Microscopy Facility for microscopy and technical support. We thank Jean-Christophe Ducom at Scripps Research High Performance Computing core for computational support. This work was supported by the National Institutes of Health (NIH) grants GM142196 and AG067594 to G.C.L. and DK046335 to J.W.K. B.B. is supported by a Postdoctoral Fellowship from the George E. Hewitt Foundation for Medical Research. Beamline 5.0.1 of the Advanced Light Source, a DOE Office of Science User Facility under Contract No. DE-AC02-05CH11231, is supported in part by the ALS-ENABLE program funded by NIH P30 GM124169-01. Editing support was provided by Emily P. Bentley.

## Data availability

Atomic coordinates and structure factors for crystal structure of the (Stilbene)_2_-TTR conjugate was deposited to the PDB with accession code 8U52. Cryo-EM maps and associated atomic models were deposited to the Electron Microscopy Databank (EMDB) and the Protein Databank (PDB), respectively, with the following EMDB and PDB IDs: unliganded canonical TTR – EMD-43162, 8VE2; unliganded compressed TTR - EMD-43163, 8VE3; unliganded frayed TTR - EMD- 43164, 8VE4; double-bound canonical TTR - EMD-43161, 8VE1; double-bound compressed TTR - EMD-43160, 8VE0; single-bound compressed TTR - EMD-43165, 8VE5; single-bound frayed TTR - EMD-43166, 8VE6.

## Supplementary Text

### Supplementary Text 1. Evidence supporting the “frayed” conformations is not artifactual

Adsorption of protein particles to the air-water interface or other hydrophobic surfaces has been shown to induce denaturation^58^. For this reason, we considered the possibility that the frayed state, which displays disordered regions, may be a product of the imaging support chemistry rather than a physiologically relevant structural state. If the frayed state was generated by adsorption, we expect the disordered region to be in contact with either the air-water interface or graphene, which would result in particle views being concentrated along that axis. If views were sufficiently localized, the sphericity of the frayed states would be significantly below ideal values 0.9-1.0, and the density would display the characteristic “smear” from preferred orientation. We find this is not the case, as sphericity for the unliganded frayed state is 0.91, and 0.8 for the (biarylamine-FT_2_-WT)_1_(C10A)_3_ TTR frayed state, and visual inspection indicates little to no “smearing” (**Supplementary Figs. 4, 19**).

Assuming the frayed state was not rigidly stuck to a hydrophobic surface, but rather was able to nutate around the point(s) of adsorption, one would expect less dramatic effects on sphericity and density quality, but views would still originate mostly at or near the perpendicular axis to the attachment surface. Such a phenomenon would be reflected on the distribution of 2D class averages, which would mostly reflect small tilts from a focal, or series of focal points. However, 2D class averages generated without realignment from the 3D-aligned particles display a wide range of views for both unliganded and (biarylamine-FT_2_-WT)_1_(C10A)_3_ TTR frayed states (**Supplementary Figs. 4, 19**). Moreover, the angular distribution of views is similar between frayed states and other states generated from the same samples (**Supplementary Figs. 4, 19**), suggesting the source of the disorder is not differential attachment to other objects in the sample. Further evidence that this frayed state is not an artifact of partial denaturation of the complex at the air-water-interface is the absence of this frayed conformation in the double-liganded sample. We expect that structural perturbations due to hydrophobic interactions would be present in all TTR datasets regardless of ligand occupancy.

### Supplementary Text 2. Transition between canonical and compressed states

Here we identify a previously unobserved conformation of TTR, which we refer to as the “compressed” state. The transition between canonical and compressed states is not dramatic in terms of its magnitude (0.91 and 0.98 Å RMSD in unliganded TTR and (Stilbene)_2_-TTR, respectively), but it involves rearrangement of the entirety of the structure. Sheet compression can be followed by the crossing (dihedral) angle between H strands of monomers on opposing sides of a binding pocket, defined by the Cα atoms of residues 114A^*→*^117A^*→*^117C^*→*^114C and 114B^*→*^117B^*→*^117D^*→*^114D.

This angle becomes smaller as the sheet extends. In the stilbene-conjugated structures, compression leads to a widening of the pocket, which is reflected in a volume change, and is captured by the distance between Ala19 alpha carbons from opposing pocket chains increasing by *∼*2 Å (**Supplementary Fig. 14**). This pocket widening, however, is not as substantial in the unliganded TTR canonical to compressed transition, in which Ala19 alpha carbon distances remain *∼*19 Å (**Supplementary Fig. 8**), closer to the value observed in the double-bound compressed state. The lack of substantial pocket shape change is also reflected in the pocket volume being relatively similar between unliganded TTR canonical and compressed states (**Fig. 1C**). The structural change that appears to compensate for sheet compaction in the compressed unliganded TTR state is the sliding of the residues that form the smaller dimer interface (formed by the strands A-B loop), shortening the distance between Val20A and Val20D by *∼*1 Å (**Supplementary Fig. 8**).

In both the unliganded TTR and (Stilbene)_2_-TTR samples, sheet compression is accompanied by the outward elongation of the EF helix, which is pulled from its C-terminus by the rigid loop connecting it with strand F, when the start of strand F and of the EF helix are brought closer by compression (**Supplementary Fig. 28**). The H-bonds of the helix become strained when the helix structure is elongated outwards, with O-N distances increasing beyond 3 Å .

Another measure of the canonical to compressed transition, alternative to individual pairwise distances, is the change of alpha carbon dihedral angles. This measure, not to be confused with the well-known Phi and Psi backbone angles, focuses on the dihedral angles described by sets of four consecutive backbone alpha carbons, and their change between two protein conformations^59^. There are significant (>30 °) changes in alpha carbon dihedrals around the F-G loop (position *∼*100) and near position 60 between canonical and compressed conformations of the unliganded TTR and (Stilbene)_2_-TTR samples (**Supplementary Figs. 29, 30**). In the transition between the unliganded forms, chain B presents a large change near position 67 (D-E loop, near C-terminus) and chain C near position 100 (F-G loop). Something similar is observed in the transition between double-bound forms: a large change is present in chain A near both positions 100 and 58 (D-E loop, near N-terminus), and in chain B near position 100. It is worth mentioning that the F-G and D-E loops are spatially close in each monomer, contributing to the same large hydrophobic cluster, which means these sequence regions could influence each other in 3D space. The backbone changes that mediate the canonical-compressed transition in unliganded TTR and (Stilbene)_2_-TTR states are variations of a same theme: the rearrangement of the core near the outer edge of each monomer β-sandwich. There are, however, small variations in the canonical-compressed transition in unliganded TTR and (Stilbene)_2_-TTR states: one or both loops can be involved in each subunit, with not all subunits necessarily undergoing changes. These small variations may underlie how the canonical-compressed transition affects pocket shape differently in the unliganded TTR and (Stilbene)_2_-TTR.

### Supplementary Text 3. Covalently modified (Stilbene)_2_-TTR

To streamline image analysis, we sought to produce a homogeneous double-covalently-liganded TTR sample, which would mitigate issues arising from sub-stoichiometric labeling or less likely artifacts arising from labeling of lysines residues in addition to 2 of 4 of the Lys15 ε-amino groups. We confirmed near-stoichiometric labeling (two ligands per tetramer, see methods) using mass spectrometry (LC-MS, see methods), which showed peaks corresponding to two stilbene-labeled and two unlabeled TTR subunits (**Supplementary Fig. 31**). Using the area under the curve from the peaks isolated by the majority ion (**Supplementary Fig. 31B**), we calculated that 43% of all subunits were labeled by stilbene, although this is likely an underestimate, as attaching a hydrophobic structural element typically lowers ionization efficiency. Assuming the worst-case scenario, where the single-stilbene-labeled TTR tetramer population is maximized, this would result in 60% double-stilbene-labeled TTR. As described in the main text, only reconstructions with both TTR pockets occupied were produced from this sample, suggesting the level of labeling achieved was higher than the ion counts would suggest. This is consistent with the results from the single-particle analysis, where only double-labeled densities were obtained, despite multiple steps of classification and refinement without imposing symmetry.

### Supplementary Text 4. Assessment of possible non-covalent binding to A2 hydrolysis products

The difficulty in fitting a covalently bound stilbene moiety in the BD side of our highest-resolution TTR reconstruction suggested averaging of multiple conformations. This is expected, based on the nature of the reconstruction process and the fact that the stilbene can conjugate with TTR by lysine residues on one or both sides of each pocket. However, it is also possible that products of spontaneous A2 hydrolysis could covalently bind to TTR. To explore this possibility, we carried out a labeling time course as described in the Methods section. The results indicated no A2 hydrolysis products were present before or after reaction with TTR (**Supplementary Fig. 33**). Most of A2 reacts with TTR in the first 30 minutes of incubation at room temperature, as no A2 hydrolysis product appears at early elution times. If present, we would expect A2 hydrolysis product to elute at earlier times, yet early peaks that would suggest hydrolysis are not observed, even after five and a half hours incubation.

### Supplementary Text 5. Accordion-like compression motion and dislodging of the 58-65 region

The two frayed structures presented in this work have a “compressed” sheet conformation, suggesting that disordering of the 58-65 region is contingent on the sheet being in a compressed conformation. We propose the coupling between sheet compression and increase in D-E loop disorder is mediated by the aromatic sidechain cluster around Phe64 (Phe44, Tyr69, and Tyr105), which becomes deeper and narrower to better accommodate Phe64 upon sheet extension, effectively pinning down the 58-65 region (**Supplementary Fig. 34**). This mechanism can be appreciated in the compressed-canonical transition for both unliganded TTR and (Stilbene)_2_-TTR states. Further supporting this idea, loops D-E and F-G, which display the largest changes in Cα dihedral angles between compressed and canonical conformations, as measured in^39^ (**Supplementary Figs. 29, 30**), seem to pull from the Tyr69 and Tyr105 side chains by the hydrogen bonds these make with the loops backbone (specifically, Tyr69 with the backbone O of position 65 and Tyr105 with the backbone N-H of position 98). It should be clarified, however, that the subtle backbone movements between compressed and canonical conformations in loops D-E and F-G are somewhat obscured by the higher B-factors at these locations; therefore, our assessment is based on a combination of Cα dihedral measurements and visual inspection. Is it likely that high-energy intermediates exist along the loop and core rearrangements, creating an energy barrier that separates the canonical and compressed conformations into discrete states. It is this discrete character that allows them to be isolated by cryo-EM.

### Supplementary Text 6. Pocket structure in the (biarylamine-FT_2_-WT)_1_(C10A)_3_ TTR sample

When we look at the differences between bound and unbound pocket geometry in the (biarylamine-FT_2_-WT)_1_(C10A)_3_ TTR frayed state, we see the bound pocket is slightly larger due to a rotation of the surrounding β-sheets along the axis perpendicular to the strands (**Supplementary Fig. 20**), narrowing the pocket in one direction and widening on another, as shown by the pairwise distances between Gly22 in opposing chains. In the (biarylamine-FT_2_-WT)_1_(C10A)_3_ TTR compressed state, the pocket volume reduction in the bound site is mediated by the opposing sheets that form the pocket coming together near the opening of the pocket, as exemplified by the distances between Lys15 alpha carbon atoms from opposing chains, which is 16.1 Å in the unbound pocket, and 13.8 Å in the bound pocket (**Supplementary Fig. 20**). These motions are yet another variation of how the TTR protomers can slide against each other across the smaller tetramer interface, in addition to sheet compression and extension. The modes of deformation seen in the (biarylamine-FT_2_-WT)_1_(C10A)_3_ TTR structures are not entirely identical to those seen in the unliganded TTR or (Stilbene)_2_-TTR states (see **Supplementary Text 2**), and indicate the shape of the TTR pockets is remarkably flexible as a result of multiple degrees of freedom granted by the protomer sheet and inter-chain interface plasticity. This plasticity explains TTR’s ability to bind a host of different ligands.

## Supplementary Figures

**Supplementary Figure 1.**
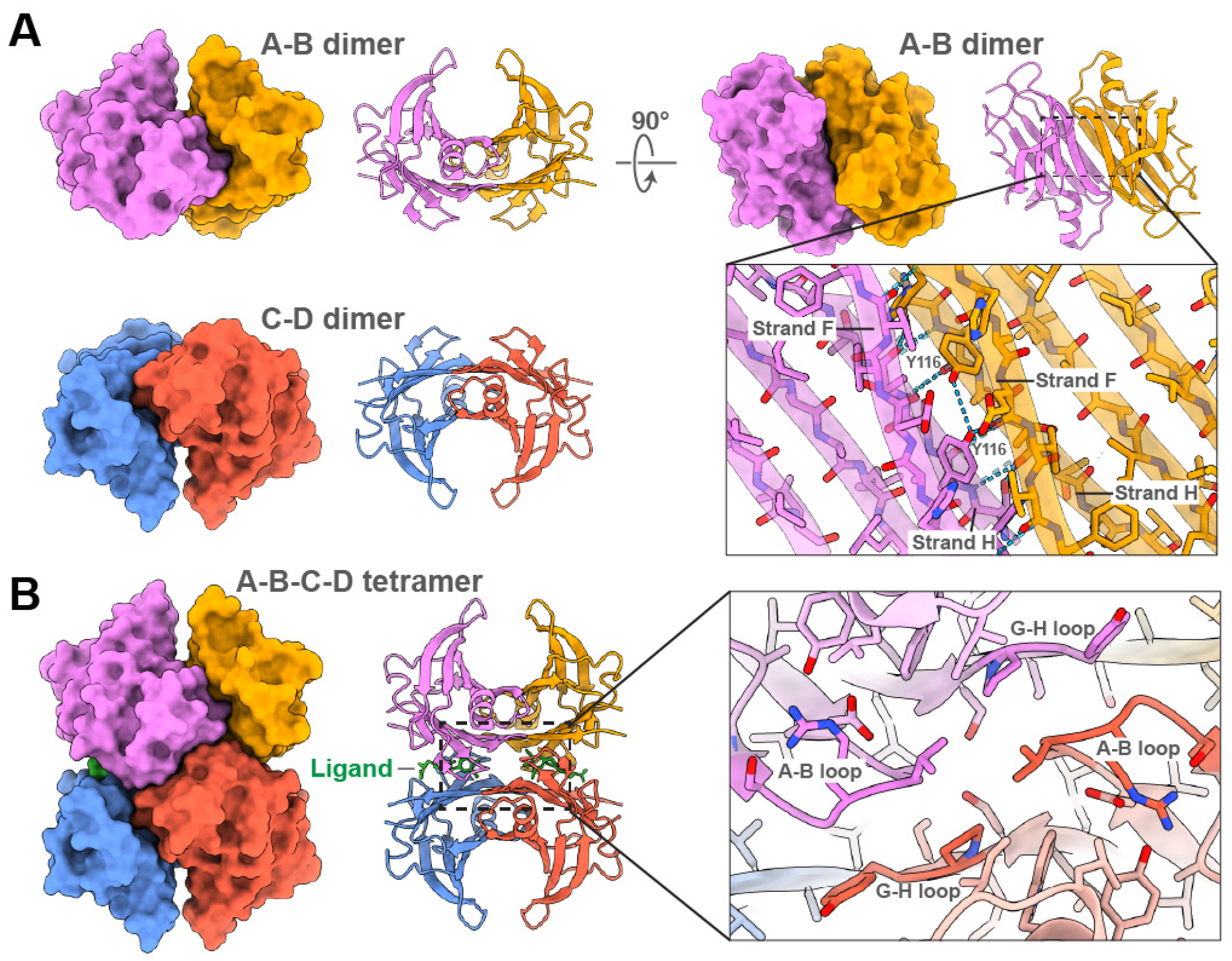
TTR quaternary structure, presented using a TTR crystal structure bound to thyroxine (PDB 1ICT). (**A**)TTR dimers, shown using molecular surface and ribbon representations, interact through an extensive hydrogen bond network between neighboring H- and F-β-strands from neighboring subunits. The inter-strand interactions are highlighted in the right panels, where the A-B dimer is viewed perpendicular to the H- and F-β-strands, and the hydrogen bond network is depicted with the atomic model overlaid with the ribbon representation. (**B**) Two TTR dimers assemble into a tetramer via interactions between the A-B and G-H loops from opposing subunits. On the left the TTR tetramer is shown as a molecular surface and ribbon representation. Thyroxine (colored green) is bound in each of the two binding pockets. On the right, a detailed view of the A-B loop interactions with the G-H loop of the opposing subunit is shown. For all images, each chain is colored as in **Fig. 1**.

**Supplementary Figure 2.**
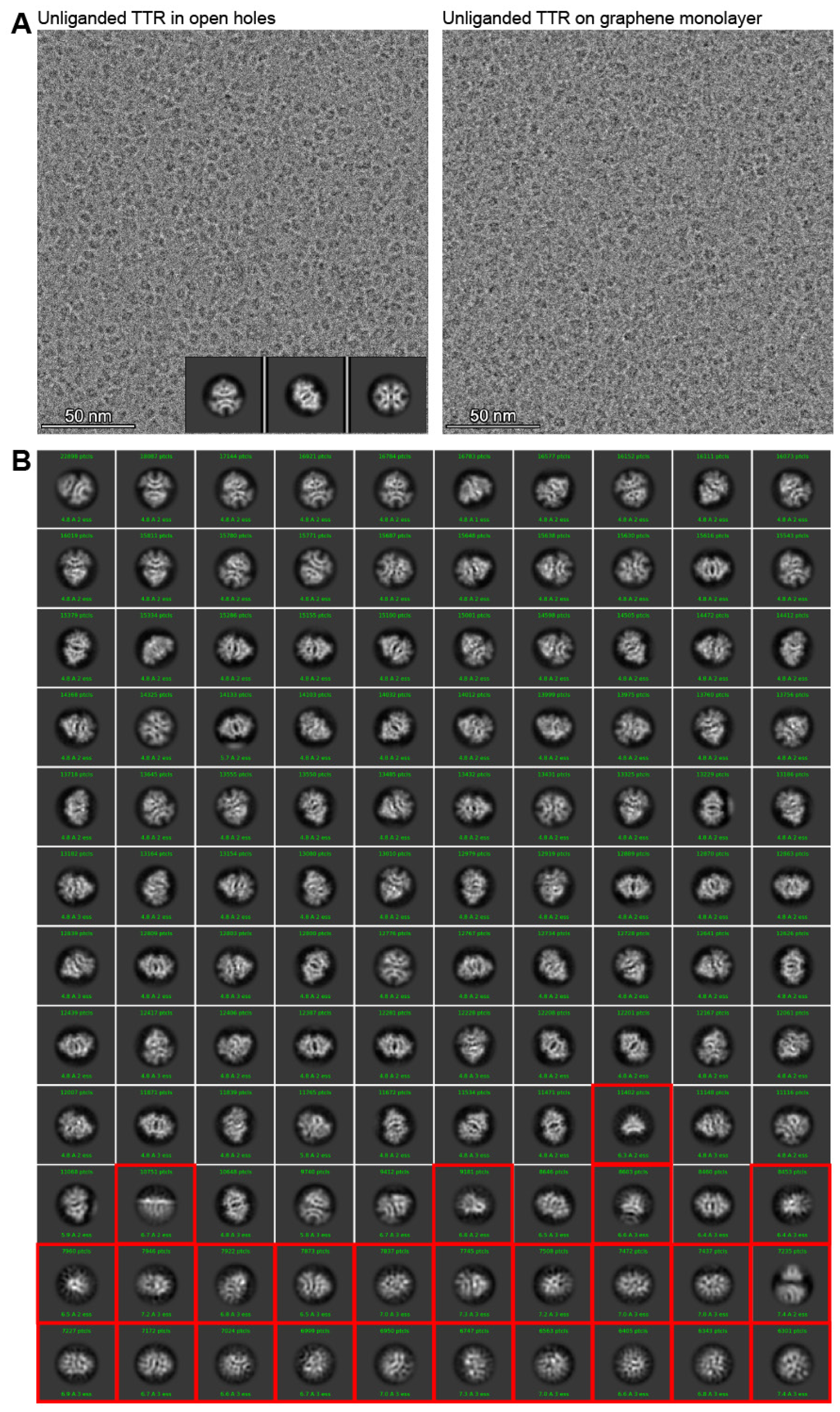
(previous page). Motion-corrected micrographs and 2D classification of unliganded TTR sample data analysis with and without graphene monolayer support. (**A**)The micrograph on the left shows unliganded TTR particles distributed in vitreous ice over an open hole in an EM grid. 2D class averages inset in the lower right show the three preferred views of TTR that were attainable over open holes. The limited views precluded ability to determine a high-resolution reconstruction. On the right is a micrograph of unliganded TTR particles acquired over a monolayer of graphene, which abolished the preferred orientation, as demonstrated by the range of views shown in 2D averages. (**B**) Final 2D classification of TTR, sorted by number of particles. Classes outlined with a red box were discarded from 3D reconstruction steps.

**Supplementary Figure 3.**
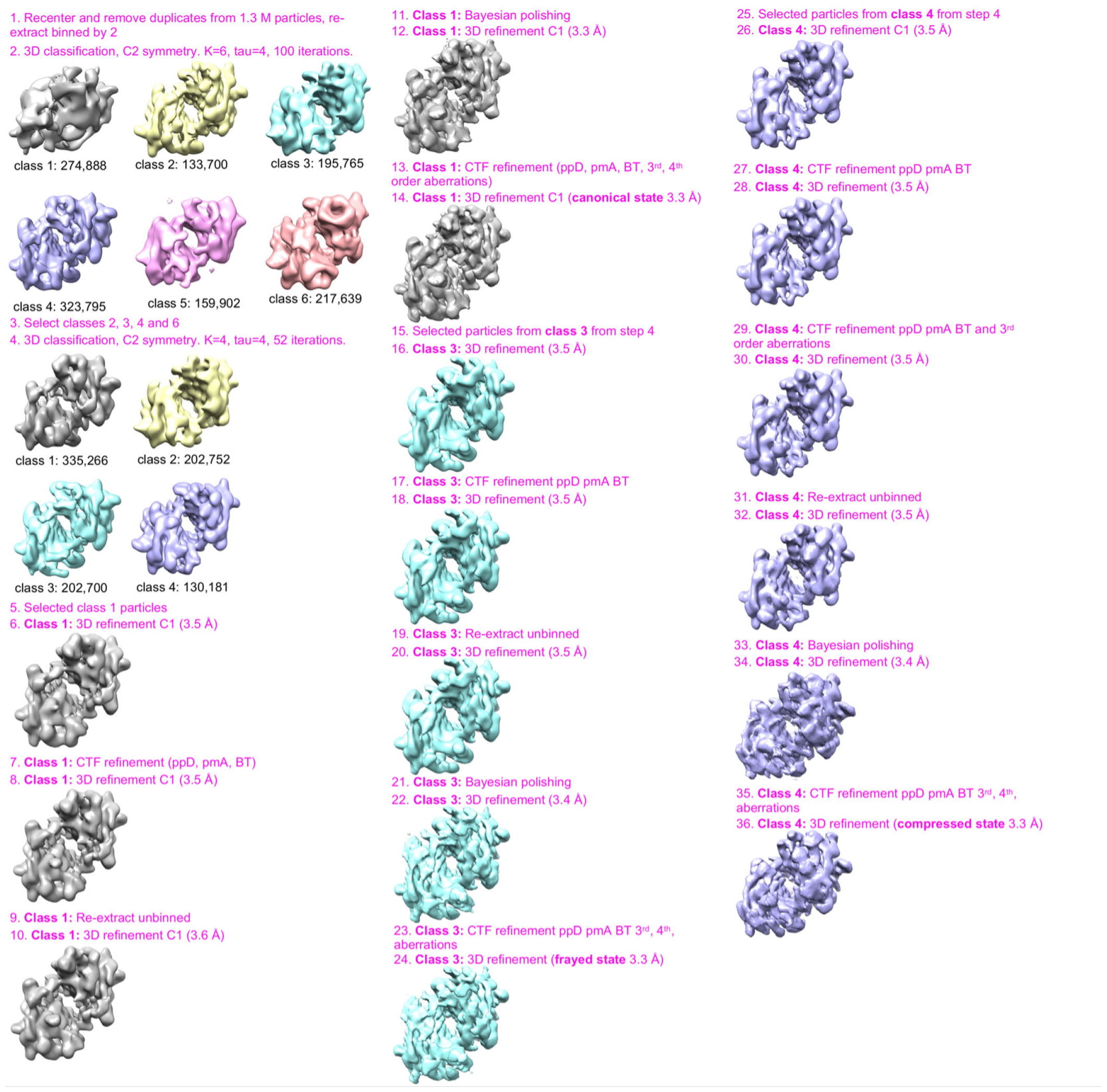
Data analysis workflow for the unliganded TTR sample, starting from import to RELION 3.1.

**Supplementary Figure 4.**
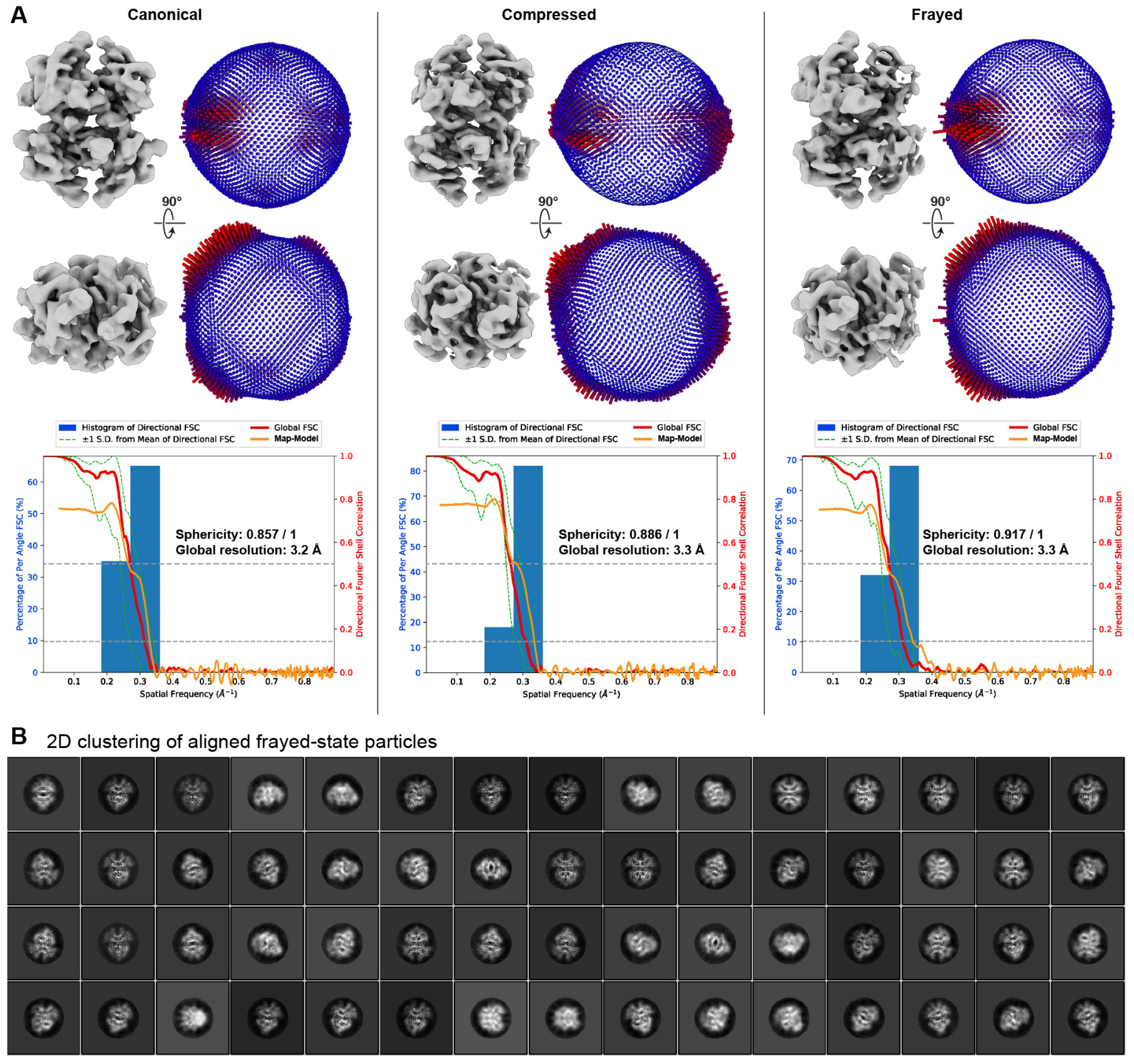
Angular distribution of views for the unliganded TTR reconstructions. (**A**)Two orthogonal views of the unfiltered reconstructions corresponding to the canonical, compressed, and frayed states of unliganded TTR are shown alongside 3D histogram of the angular distribution. Below, the directional Fourier Shell Correlation is shown, denoting the sphericity value and estimated global resolution. The map-to-model FSC is overlaid in orange, with the FSC at 0.5 shown as a dashed line. (**B**) 2D averages generated without alignment from particles from the unliganded frayed class, after 3D reconstruction.

**Supplementary Figure 5.**
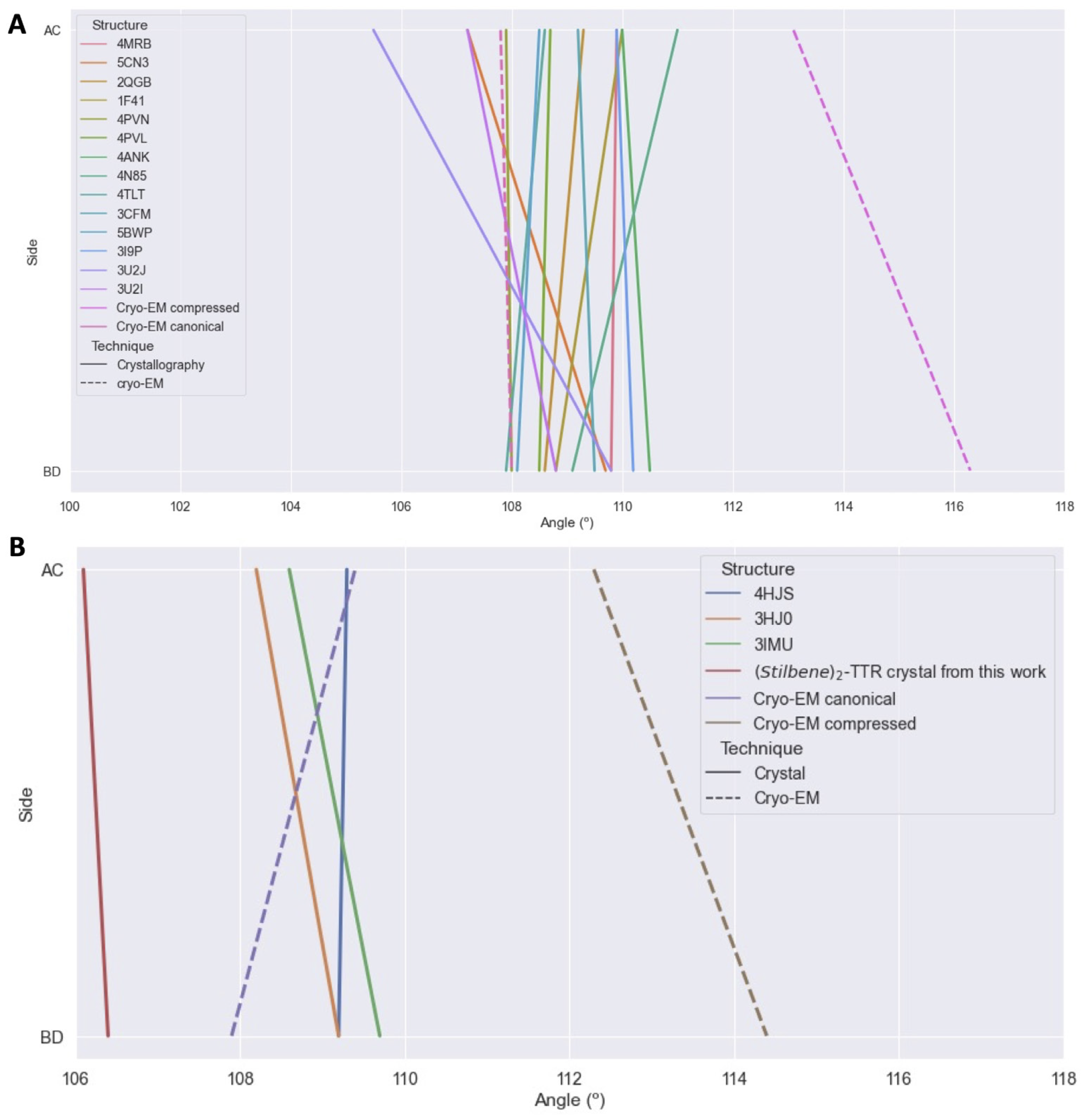
Comparison of H-strand cross angles for our cryo-EM structures and published crystal structures of unliganded and covalently liganded TTR. (**A**)The dihedral angles between opposing H β-strands, measured as shown in Fig. 1E, are plotted as a line with dihedral angle value of the A / C subunits as the upper point and the B / D subunit dihedral angle as the lower point. The PDB codes for previously characterized unliganded complexes are labeled in the figure inset. The majority of the crystal structures have values that are similar to the cryo-EM structure we refer to as the canonical state. (**B**) The same measurement of dihedral angles for our (Stilbene)_2_-TTR cryo-EM structure and published covalently liganded TTR crystal structures are plotted as in (A). The PDB codes for previously characterized complexes are labeled in the figure inset.

**Supplementary Figure 6.**
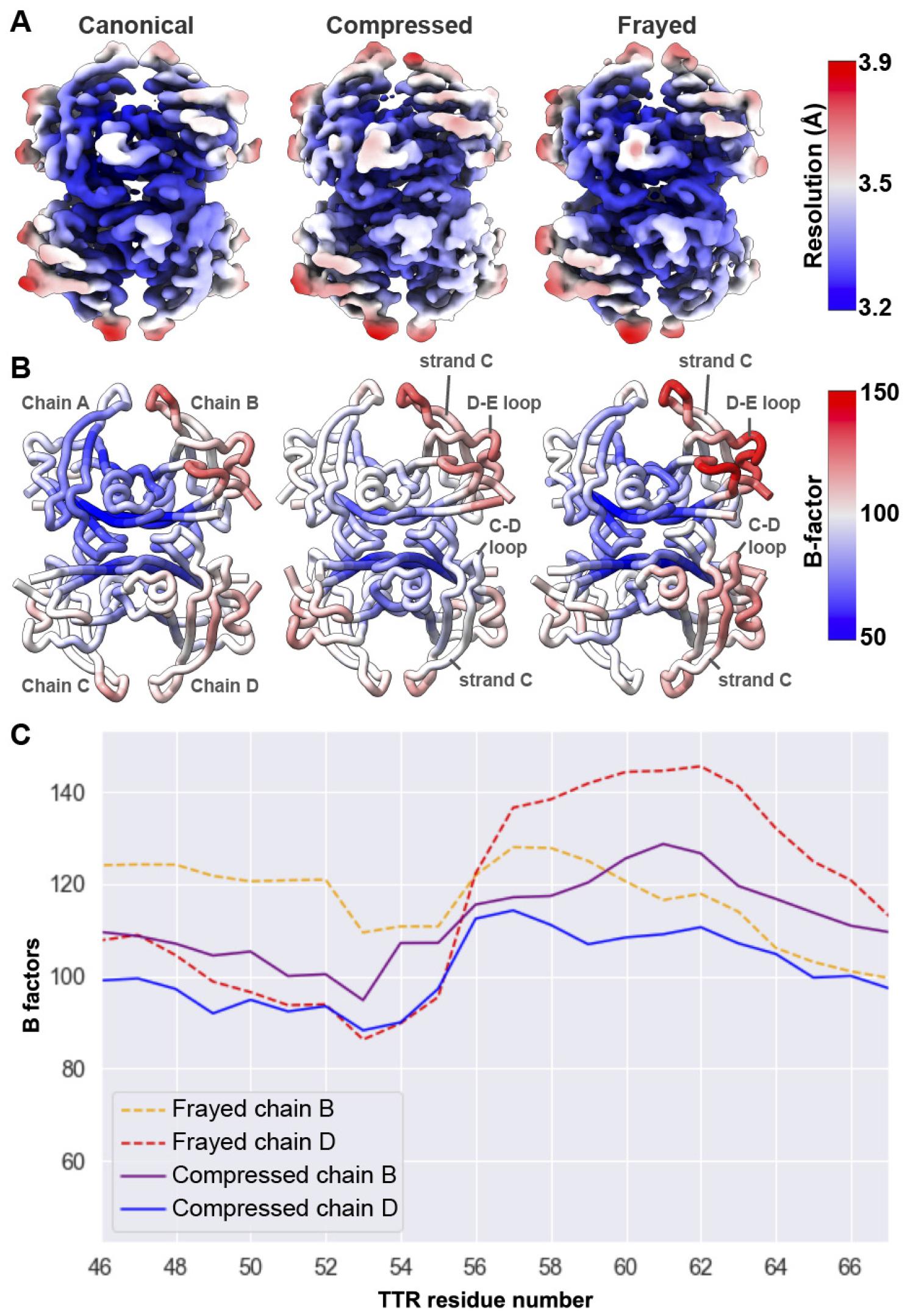
Local resolution and B-factors for the unliganded TTR conformations. (**A**)Reconstructions of the three unliganded TTR conformations are colored according to the estimated local resolution, as calculated by RELION. (**B**) Models for the unliganded TTR states shown as backbone “tube” cartoons, colored by B-factor using the same scale. Chains B in all structures have the highest B-factors in the structures, and these values are substantially higher in the frayed state. (**C**) B-factors are presented for residue numbers in areas that appear most disordered in the frayed state. B-factors for the same residues in the compressed state are presented for comparison.

**Supplementary Figure 7.**
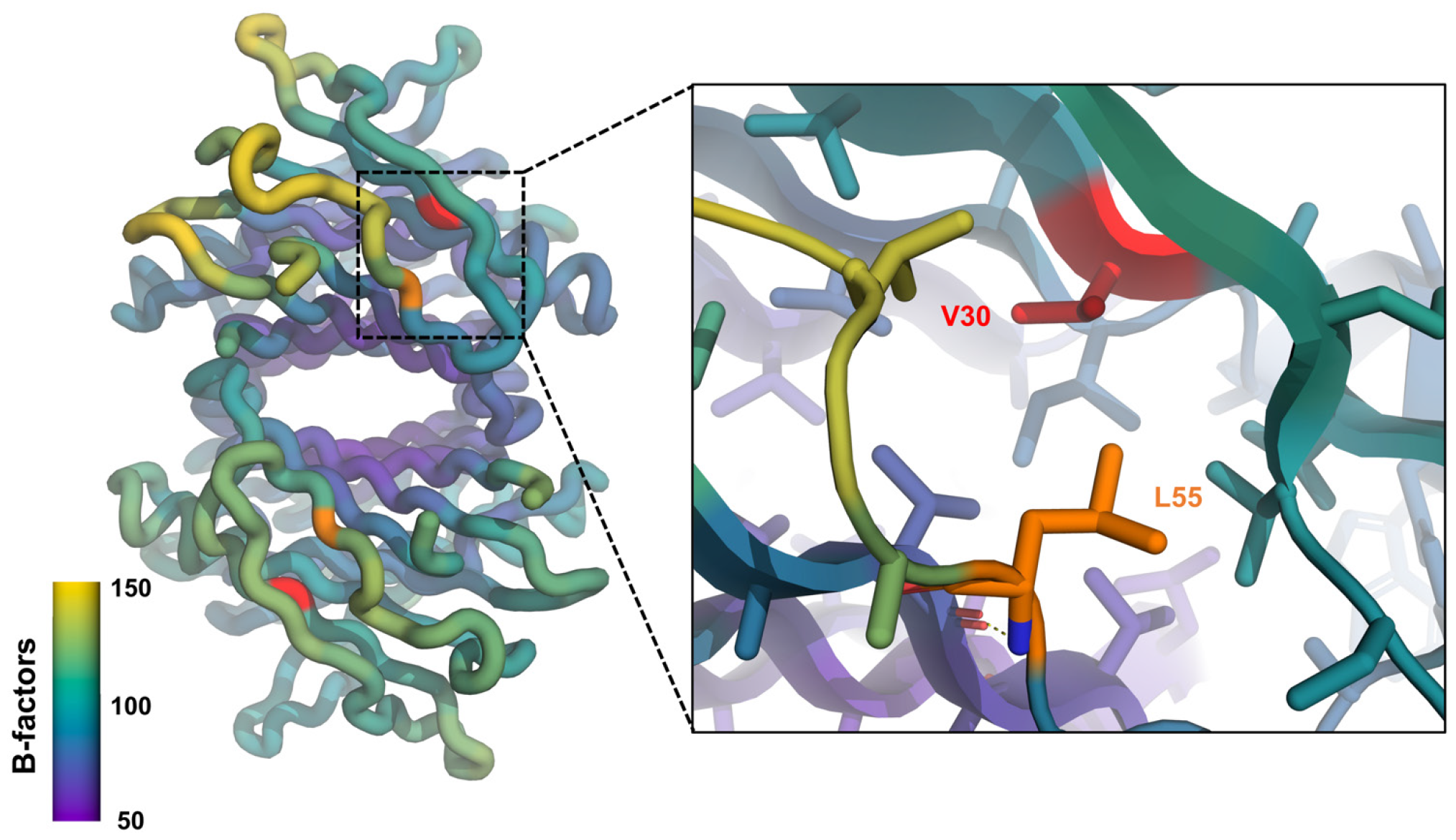
Location of common hATTR mutations (V30M and L55P) in the context of the unliganded TTR model based on the frayed state density. The backbone-only cartoon model (left) is colored by B-factor values, and an inset to the right highlights hATTR mutation sites V30 and L55 as sticks. The V30P mutation results in a larger residue in the core of TTR, near the frayed region. The L55P mutation abolishes a hydrogen bond (dashed lines in the inset) between position 55 and 14, which weakens the association of strand D and adjacent loops with the rest of the structure.

**Supplementary Figure 8.**
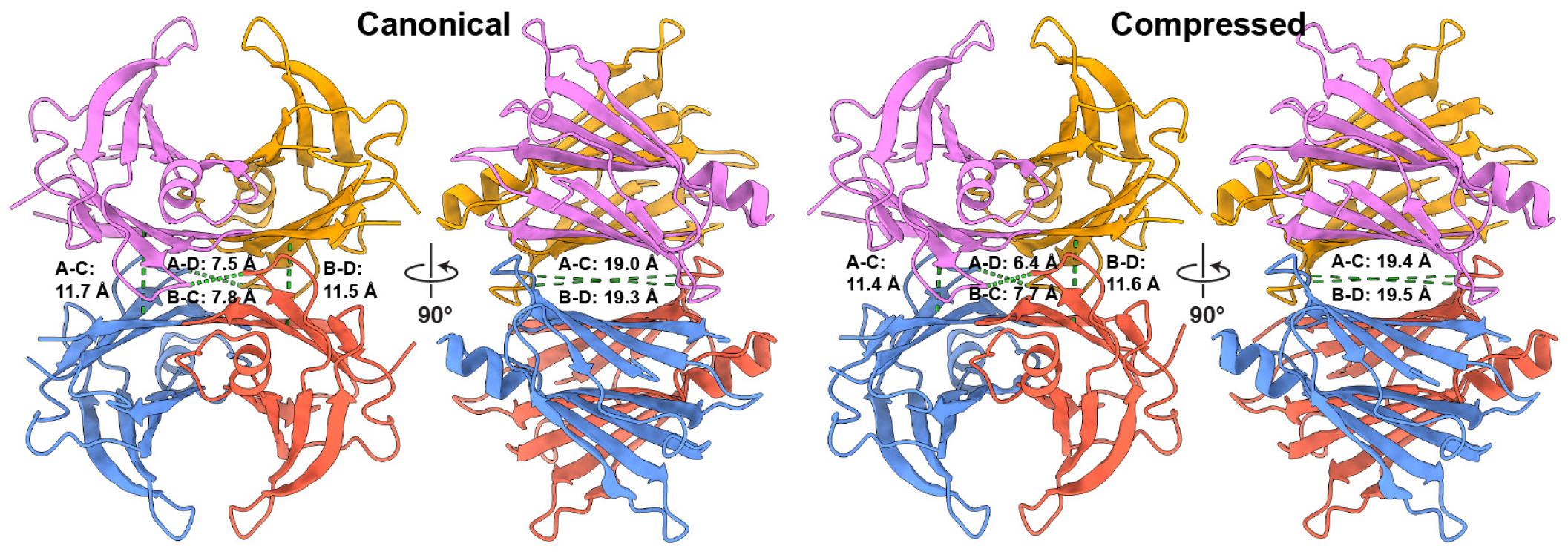
Differences in pocket geometry between canonical and compressed conformations of unliganded TTR. Ribbon representation of the unliganded TTR conformations, with subunits colored as in Fig. 1 and pairwise distances between Cα shown as dotted green lines. On the left, distances between Cα atoms of Val20 between the subunits A-D and B-C are shown at the center, and measured distances between the Cα atoms of Ala108 are shown at the periphery. On the right the distances between the between the Cα atoms of Ala19, oriented horizontally across the binding pockets, are shown.

**Supplementary Figure 9.**
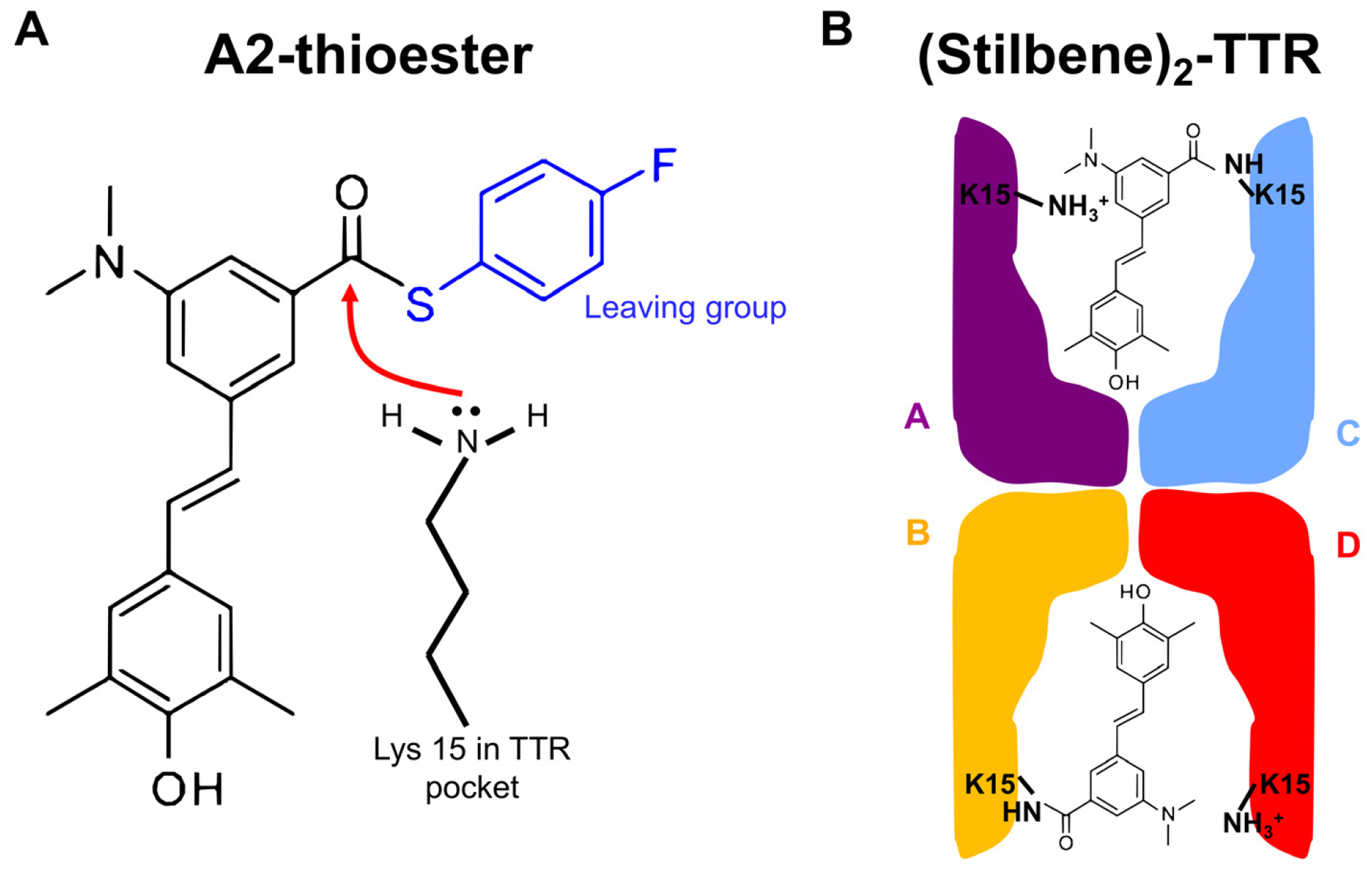
TTR reaction with A2 to afford the (Stilbene)_2_-TTR conjugate. (**A**)Line drawing of the reaction between the A2-thioester with the unprotonated lysine 15 e-amine of TTR acting as a nucleophile. The 4-fluorothiolphenolate serves as the leaving group (blue). (**B**) Cartoon depiction of the (Stilbene)_2_-TTR conjugate, covalently linked through subunits C and B. Another possible configuration would be the stilbene moiety covalently linked to subunits C and D or A and B.

**Supplementary Figure 10.**
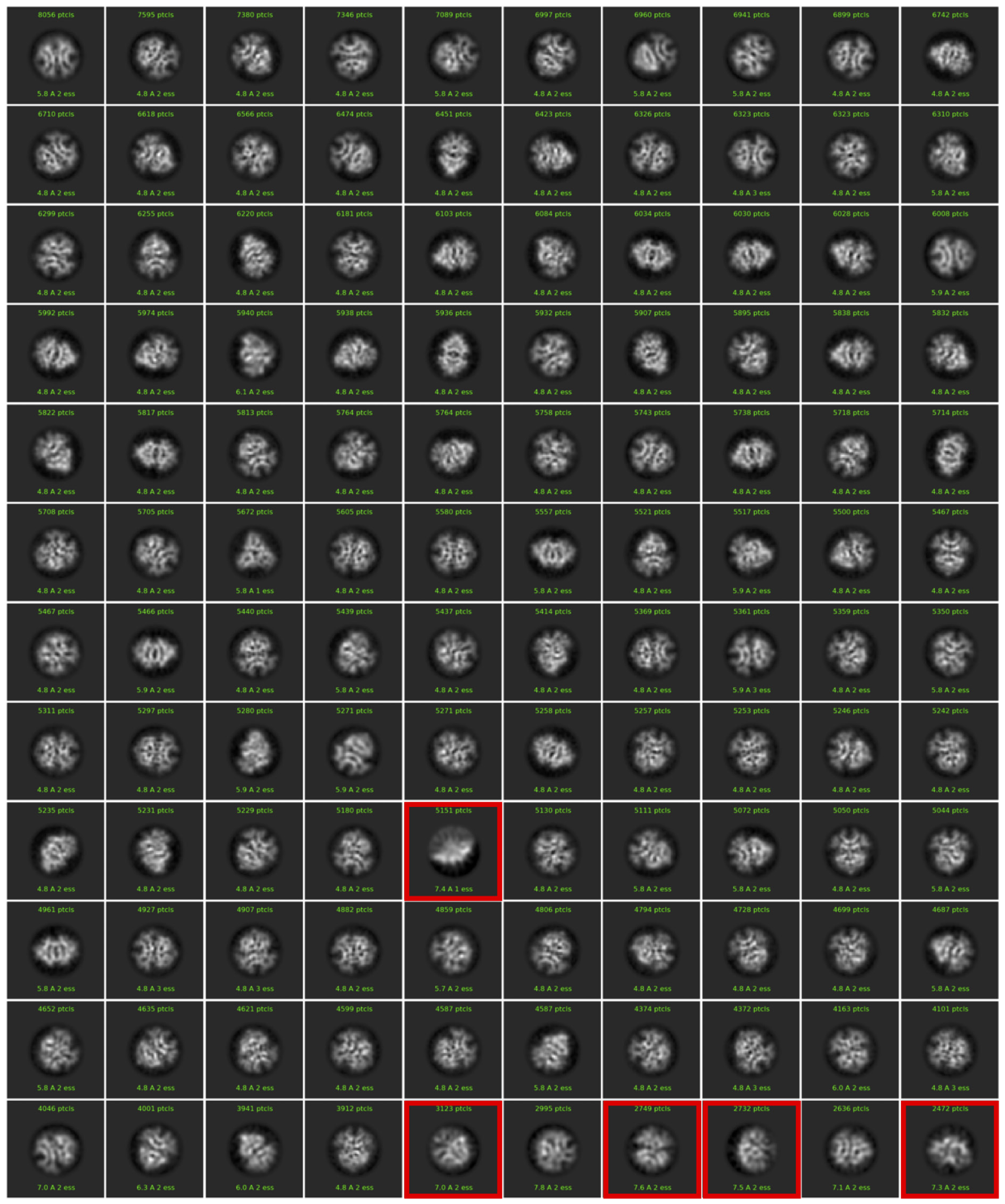
Final 2D classification step for the (Stilbene)_2_-TTR sample data analysis. Classes are sorted by number of particles. Classes in a red box were discarded.

**Supplementary Figure 11.**
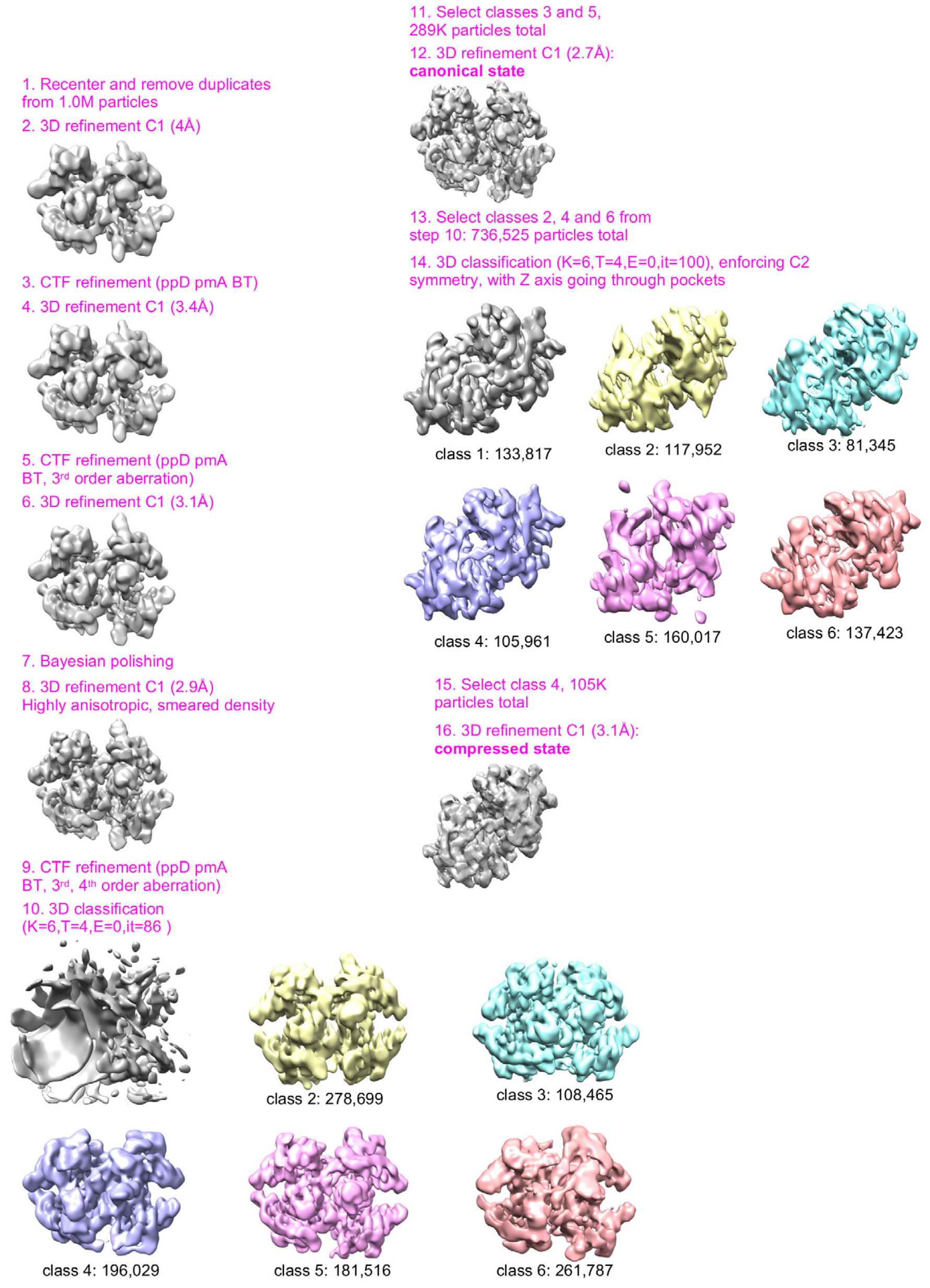
Data analysis workflow for the (Stilbene)_2_-TTR sample, starting from import to RELION 3.1.

**Supplementary Figure 12.**
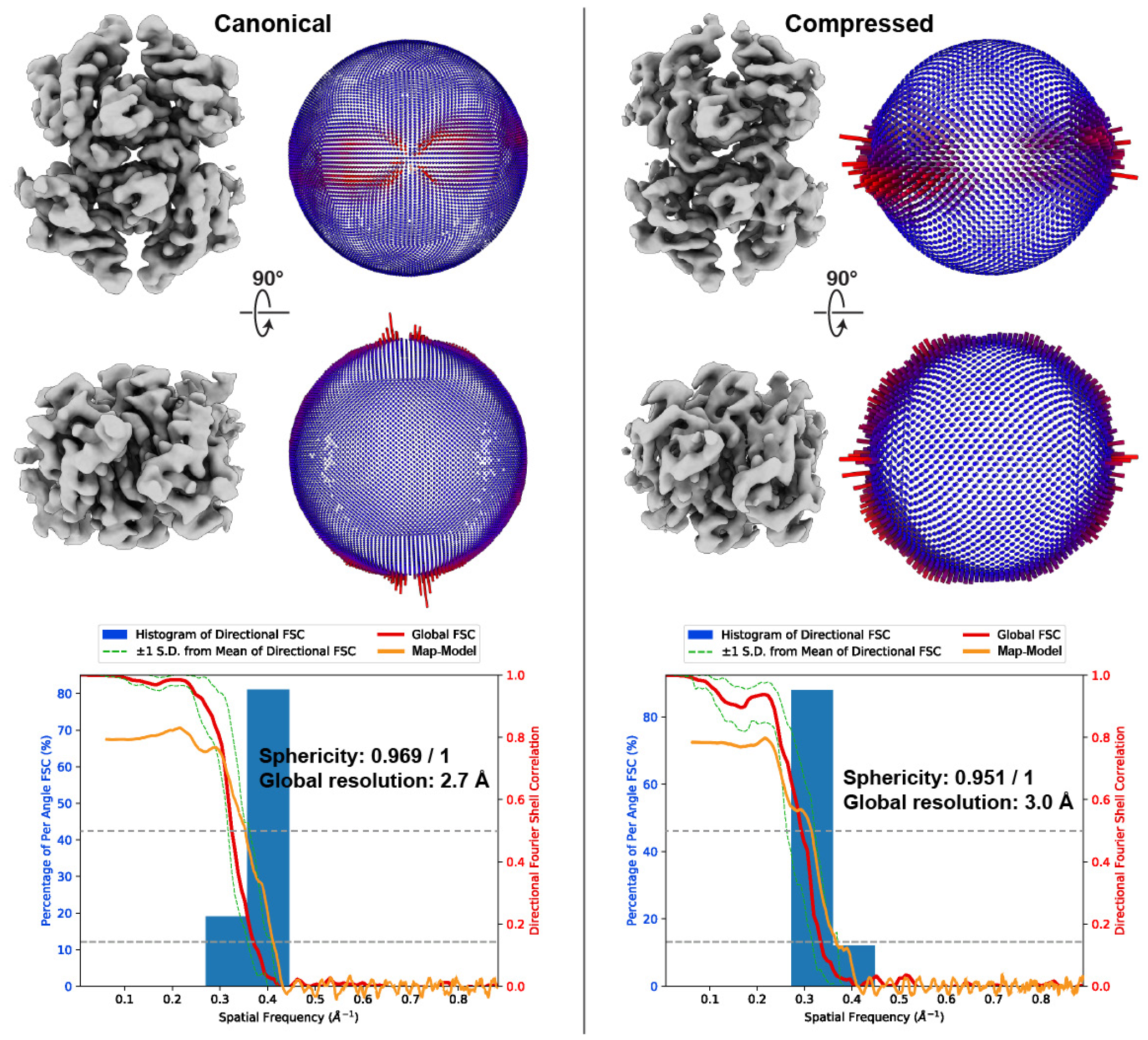
Angular view distribution and directional FSCs for the (Stilbene)_2_-TTR conjugate sample reconstructions. Two orthogonal views of the unfiltered reconstructions corresponding to the canonical and compressed states of the double-liganded TTR are shown alongside 3D histogram of the angular distribution. Below, the directional Fourier Shell Correlation is shown, denoting the sphericity value and estimated global resolution. The map-to-model FSC is overlaid in orange, with the FSC at 0.5 shown as a dashed line.

**Supplementary Figure 13.**
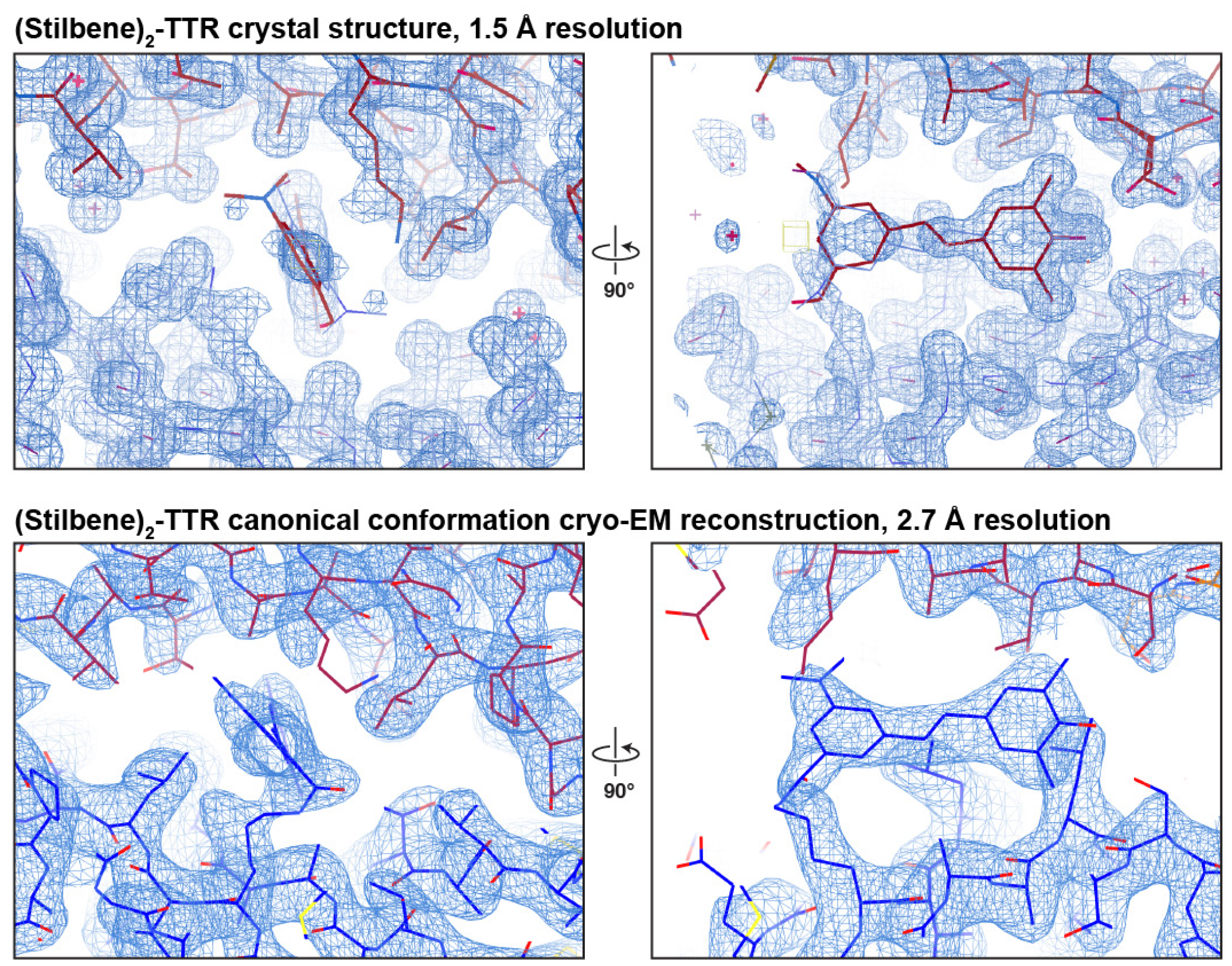
Stilbene conjugation site in the binding pocket formed by the A and C subunits determined by crystallography and cryo-EM. The top panels show electron density below noise level from a 1.5 Å resolution crystal structure. The stilbene moiety seats at a crystallographic symmetry axis, resulting in two-fold averaging of the density. The bottom panels show the same region in our cryo-EM canonical state reconstruction, which was generated without imposing symmetry, showing the asymmetric binding pose of the ligand.

**Supplementary Figure 14.**
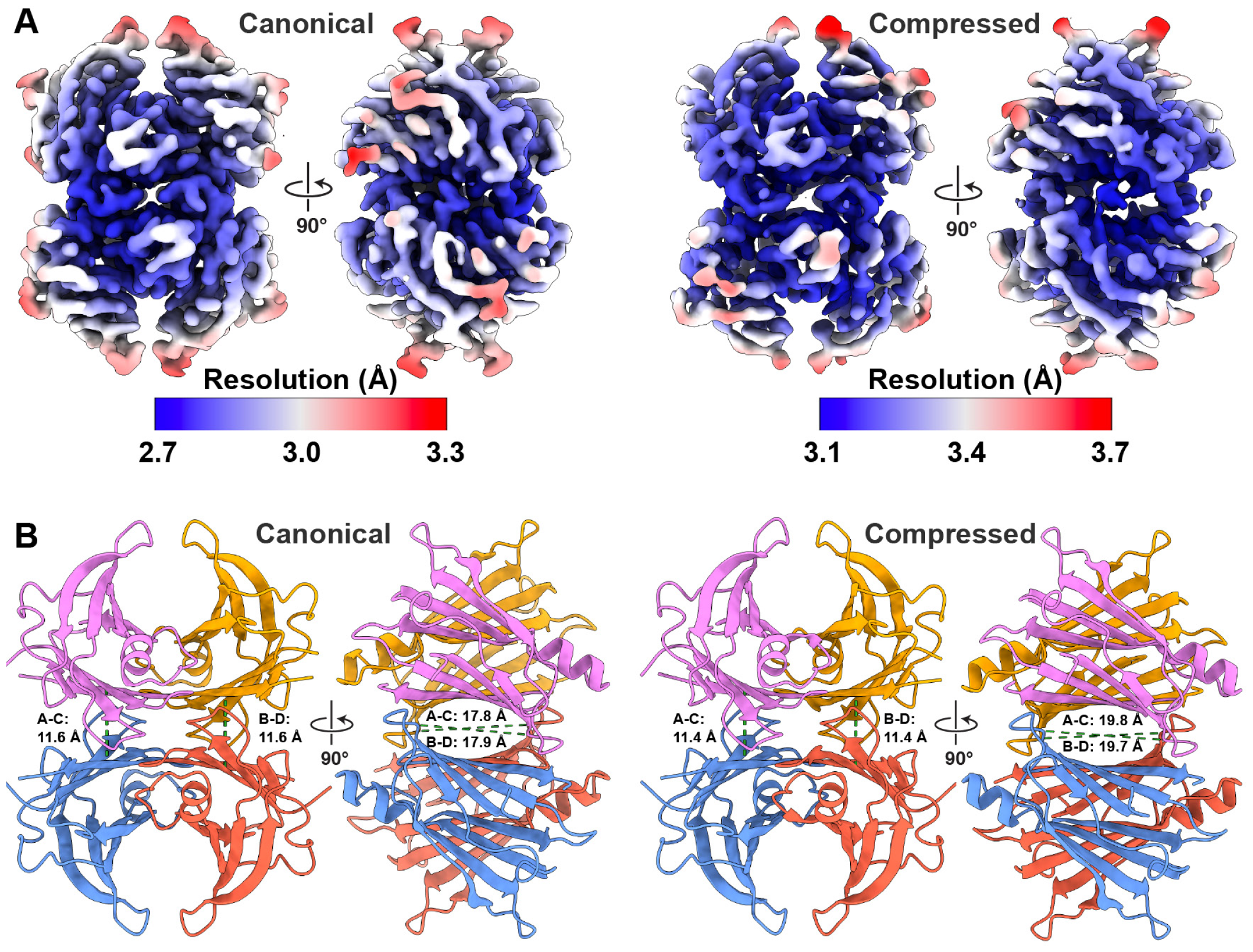
Local resolution and differences in pocket dimensions between canonical and compressed conformations of the (Stilbene)_2_-TTR conjugate. (**A**)Reconstructions of the canonical and compressed double-liganded TTR conformations are colored according to the estimated local resolution, as calculated by RELION. **B**. Ribbon representation of the double-liganded TTR conformations, with subunits colored as in **Fig. 1** and pairwise distances between Cα shown as dotted green lines. On the left the distances between the Cα atoms of Ala108, which are oriented vertically across the binding pockets, are measured and shown. On the right the distances between the between the Cα atoms of Ala19, oriented horizontally across the binding pockets, are shown.

**Supplementary Figure 15.**
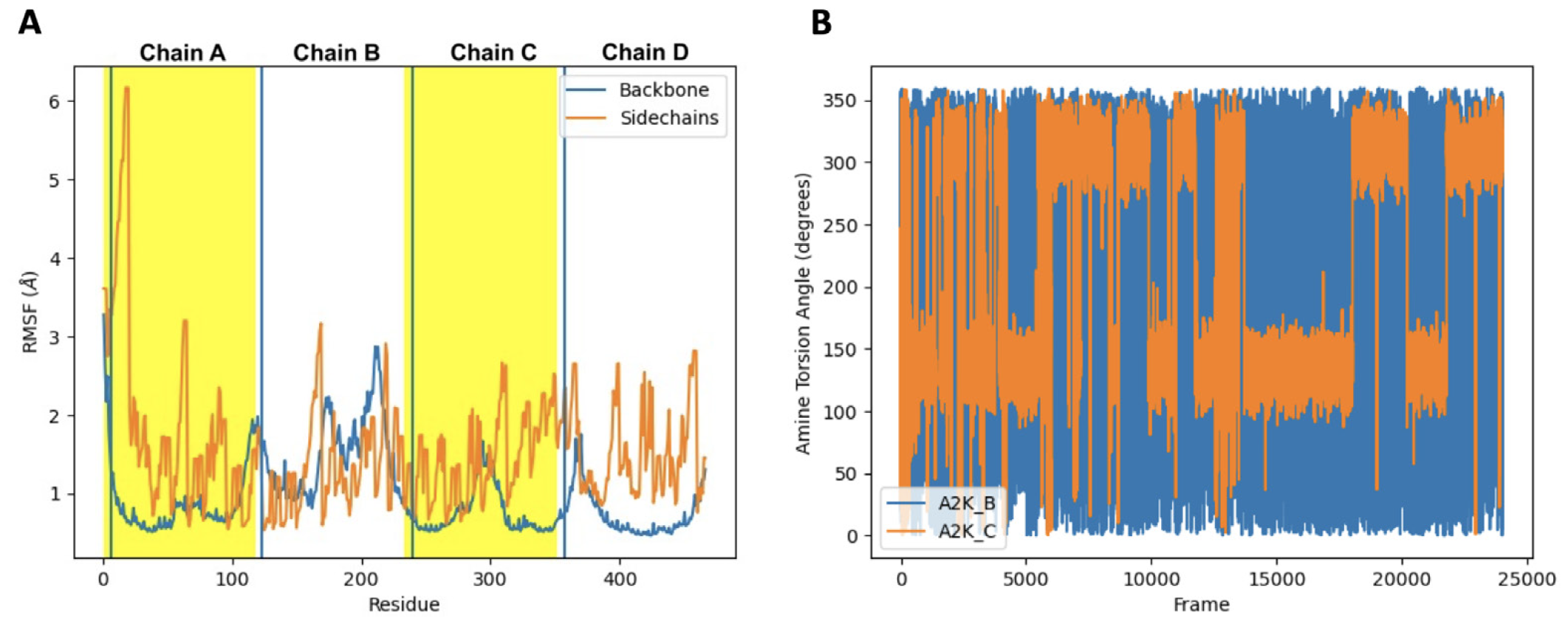
Backbone atoms around Stilbene-K15 and Stilbene amine group exhibit greater flexibility in the larger binding pocket. (**A**)Root mean squared fluctuation of backbones and side chain atoms in the molecular dynamics simulations for the canonical state of the (Stilbene)_2_-TTR covalent complex. Values corresponding to each chain are labeled with yellow or white background, as well as by chain name on the top. The position corresponding to residue 15 (either modified lysine or A2K stilbene-modified lysine) is labeled with arrows. The red arrow (left, chain B) corresponds to stilbene moiety residing in the larger pocket and the blue arrow (right, chain C) that in the smaller pocket. (**B**) Torsion angle of stilbene tertiary amine. A2K B (blue) corresponds to the stilbene moiety in the larger pocket and may rotate freely within it. A2K C (orange) resides in the smaller pocket and is mostly constrained to two torsions 180 ^*°*^ apart.

**Supplementary Figure 16.**
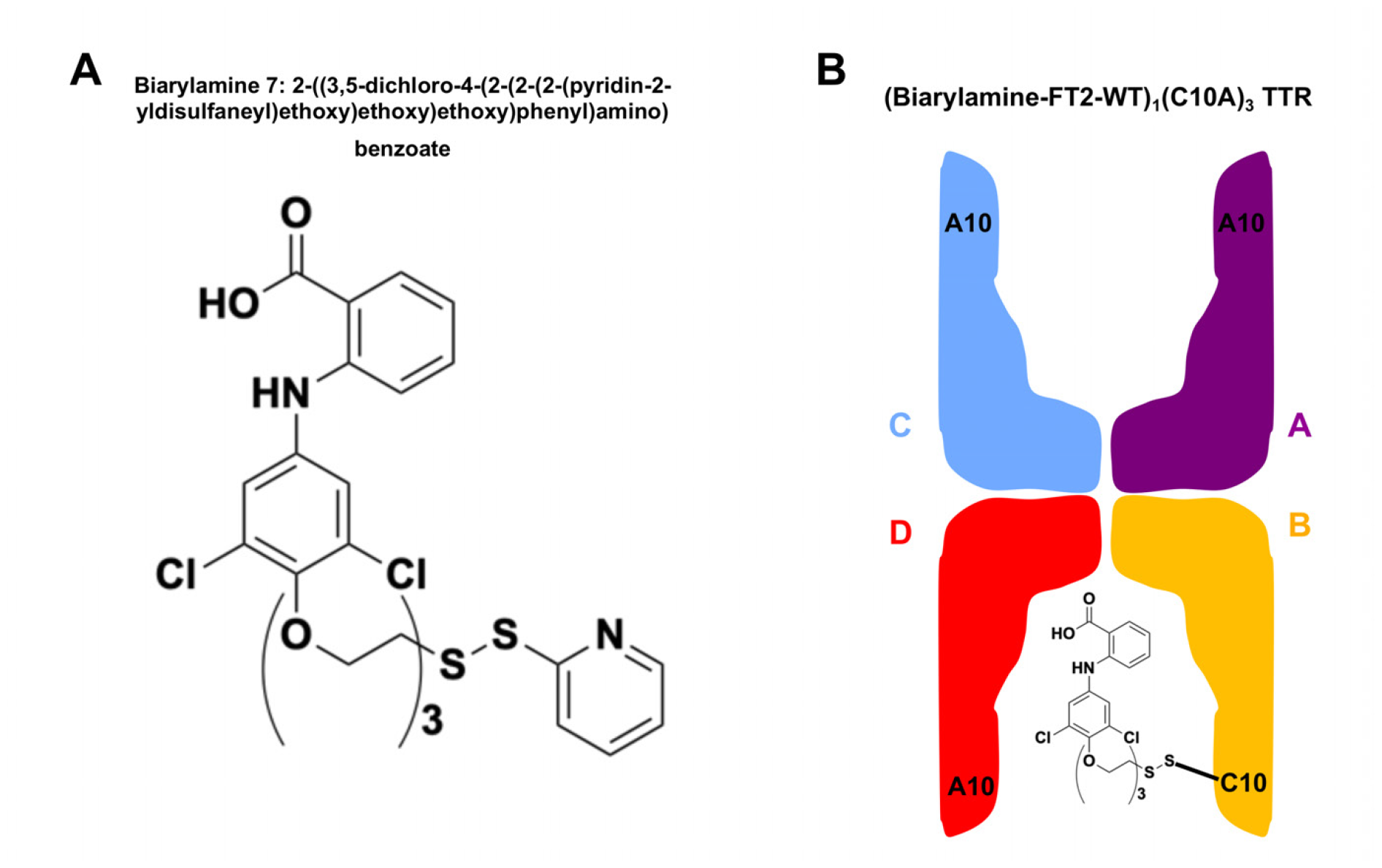
TTR reaction with Biarylamine 7 ligand (2-((3,5-dichloro-4-(2-(2-(2-(pyridin-2-yldisulfaneyl)ethoxy) ethoxy)ethoxy)phenyl)amino)benzoate) to afford the (biarylamine-FT_2_-WT)_1_(C10A)_3_ TTR conjugate. (**A**)Chemical structure of Biarylamine 7 ligand. (**B**) Cartoon depiction of the (biarylamine-FT_2_-WT)_1_(C10A)_3_ TTR adduct, covalently linked through subunit B.

**Supplementary Figure 17.**
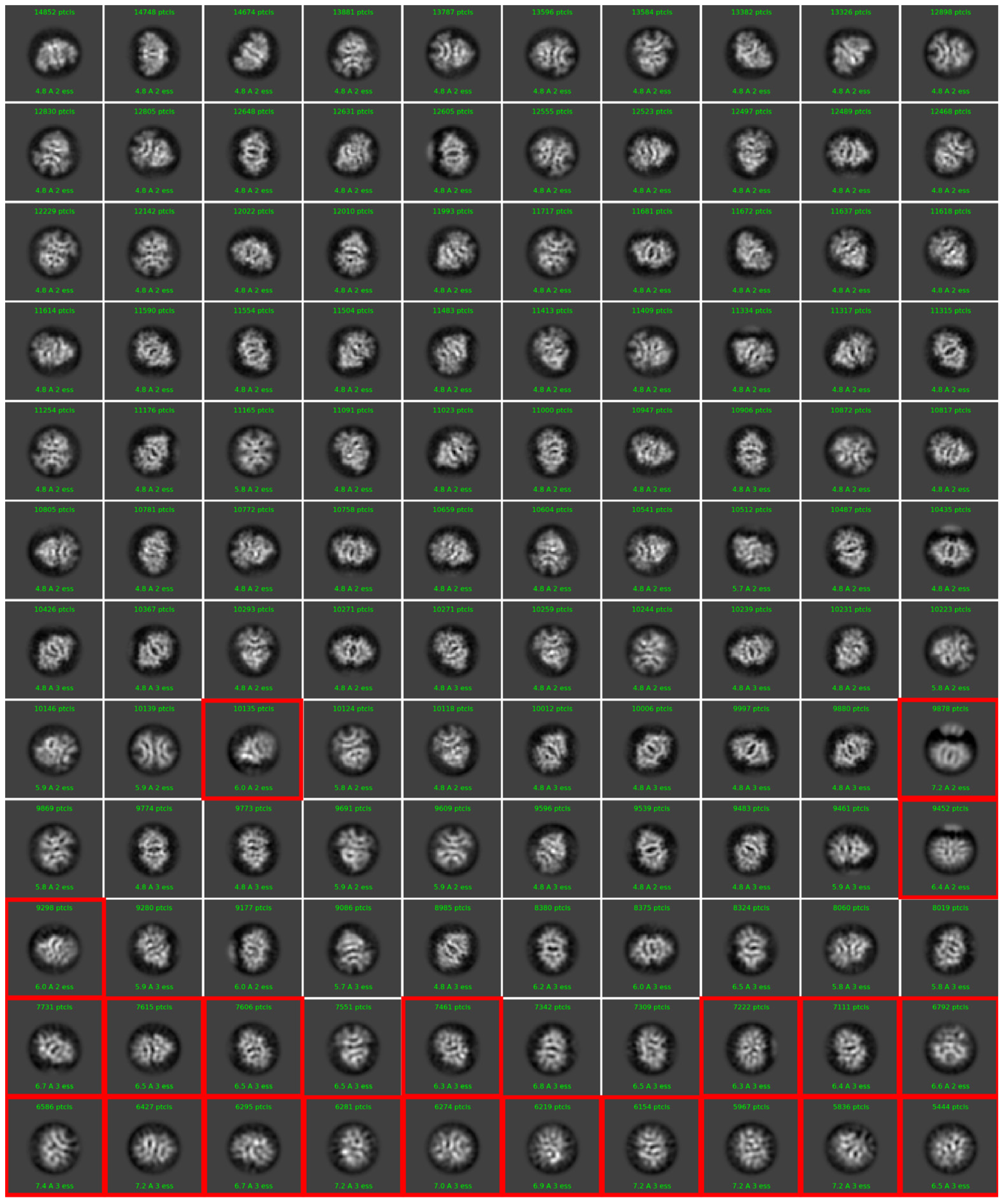
Final 2D classification step for the (biarylamine-FT_2_-WT)_1_(C10A)_3_ TTR sample data analysis. Classes are sorted by number of particles. Classes in a red box were discarded.

**Supplementary Figure 18.**
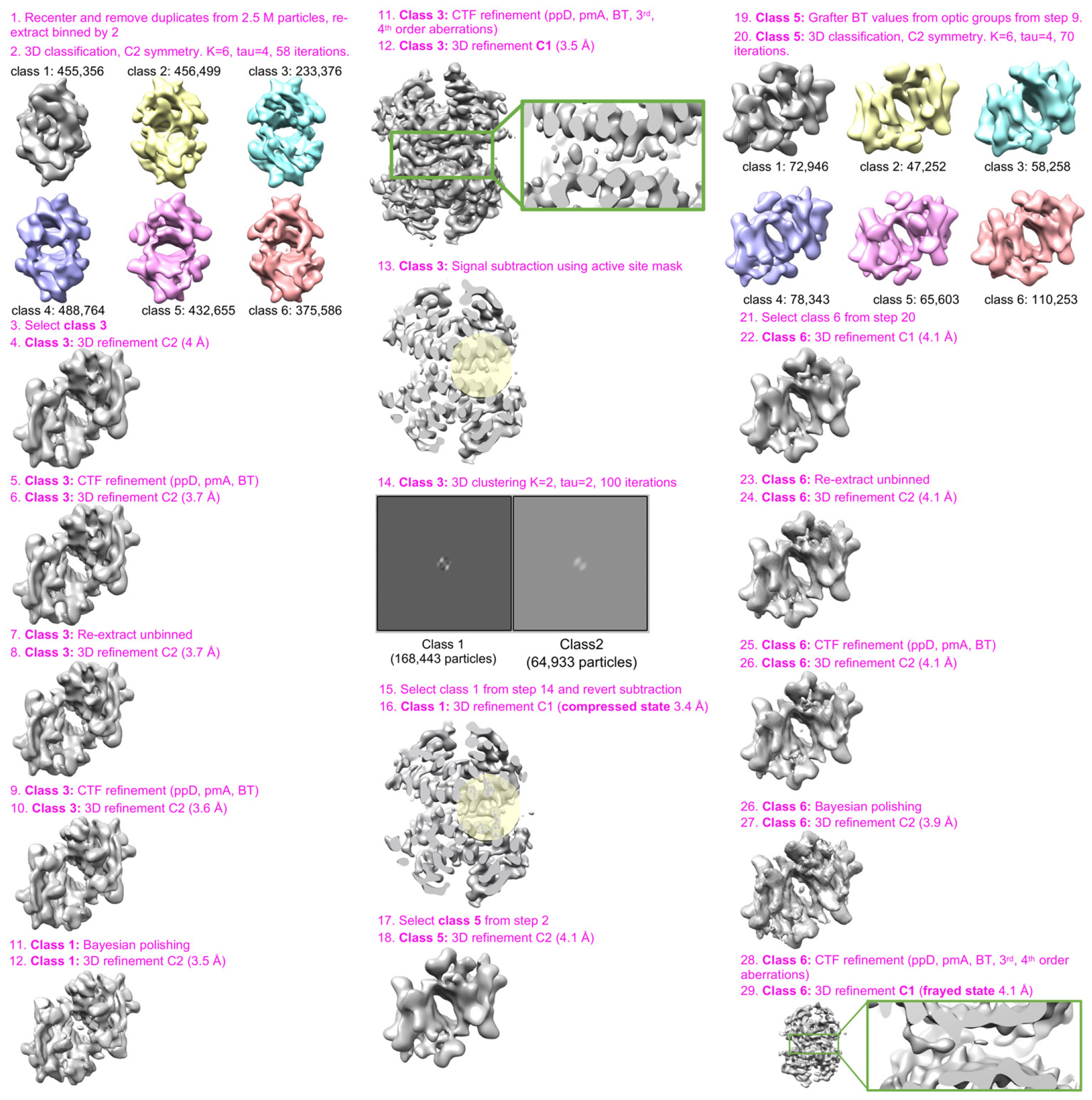
Data analysis workflow for the (biarylamine-FT_2_-WT)_1_(C10A)_3_TTR sample, starting from import to RELION 3.1.

**Supplementary Figure 19.**
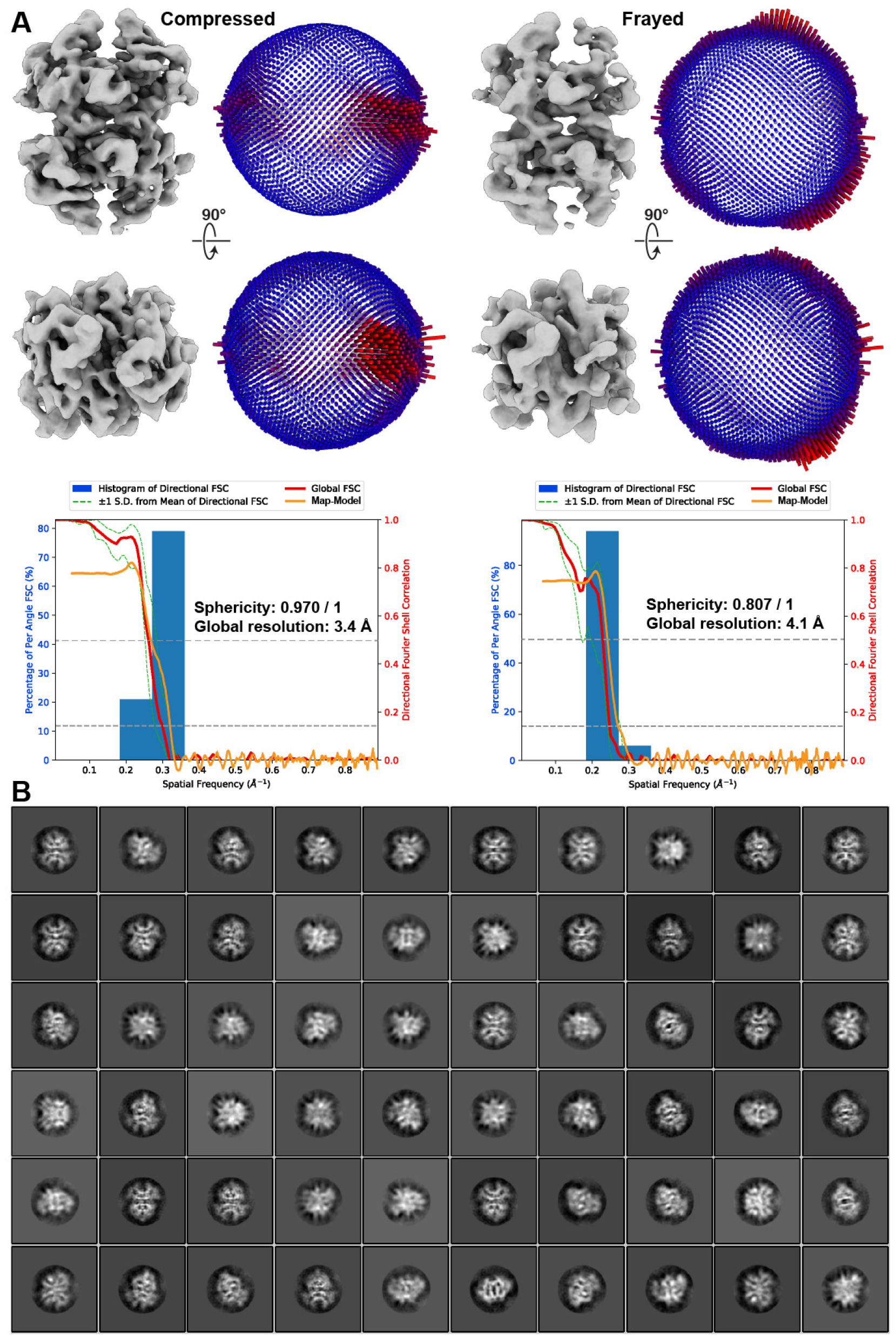
Angular distribution of views for the (biarylamine-FT_2_-WT)_1_(C10A)_3_TTR compressed and frayed states. (**A**)Two orthogonal views of the unfiltered reconstructions corresponding to the compressed and frayed states of single-liganded TTR are shown alongside 3D histogram of the angular distribution. Below, the directional Fourier Shell Correlation is shown, denoting the sphericity value and estimated global resolution. (**B**) 2D averages generated without alignment from particles from the (biarylamine-FT_2_-WT)_1_(C10A)_3_ TTR frayed class, after 3D reconstruction. The map-to-model FSC is overlaid in orange, with the FSC at 0.5 shown as a dashed line.

**Supplementary Figure 20.**
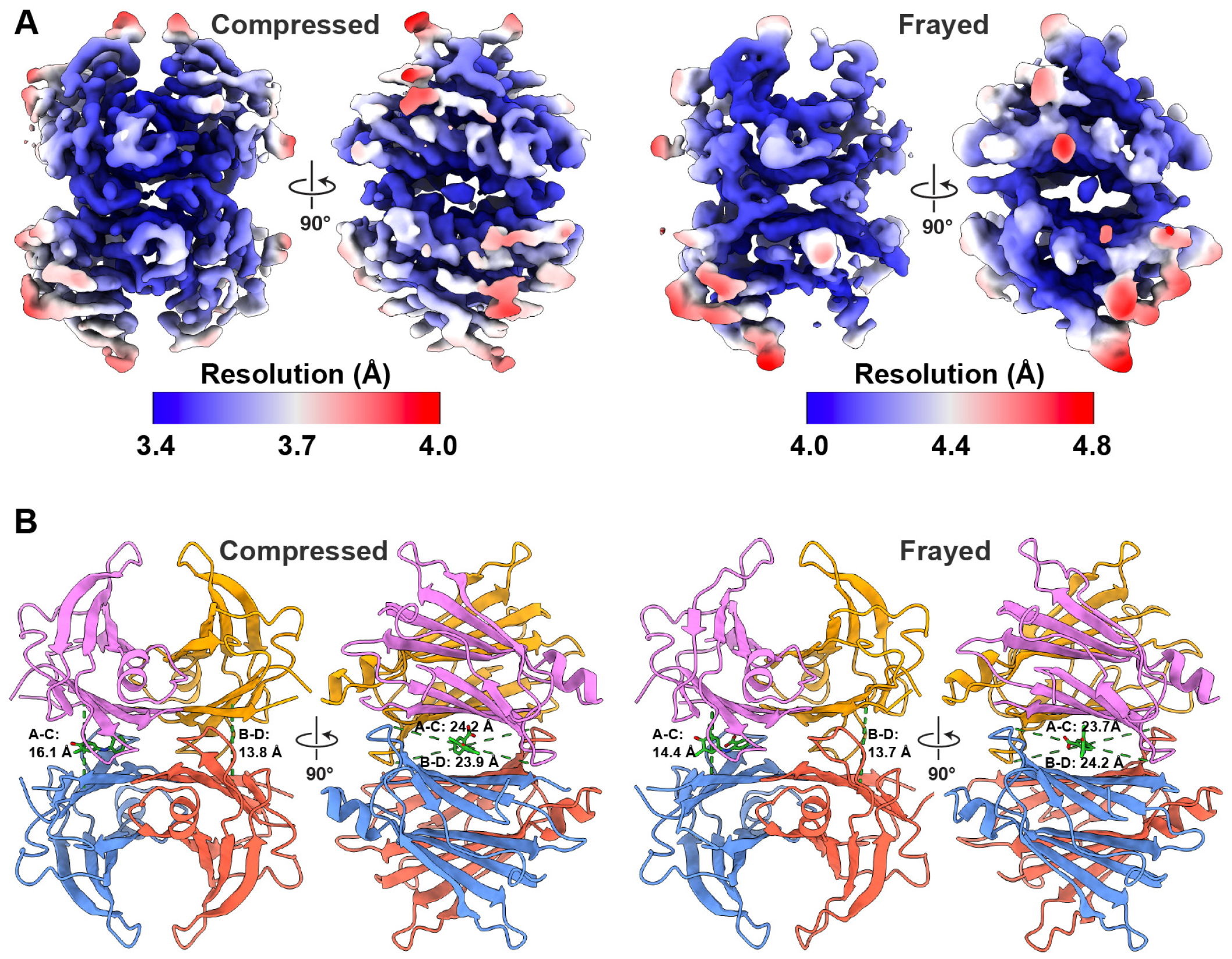
Local resolution and differences in pocket geometry between occupied and unoccupied binding sites in the (biarylamine-FT_2_-WT)_1_(C10A)_3_ TTR structures. (**A**)Reconstructions of the compressed and frayed single-liganded TTR colored according to the estimated local resolution, as calculated by RELION. (**B**) Ribbon representation of the single-liganded compressed and frayed TTR conformations, with subunits colored as in Fig. 1, and pairwise distances between Cα shown as dotted green lines. On the left, measured distances between Cα atoms of Lys15 between the subunits A-C and B-D are shown. On the right the distances between the between the Cα atoms of Gly22, oriented horizontally across the binding pockets, are shown.

**Supplementary Figure 21.**
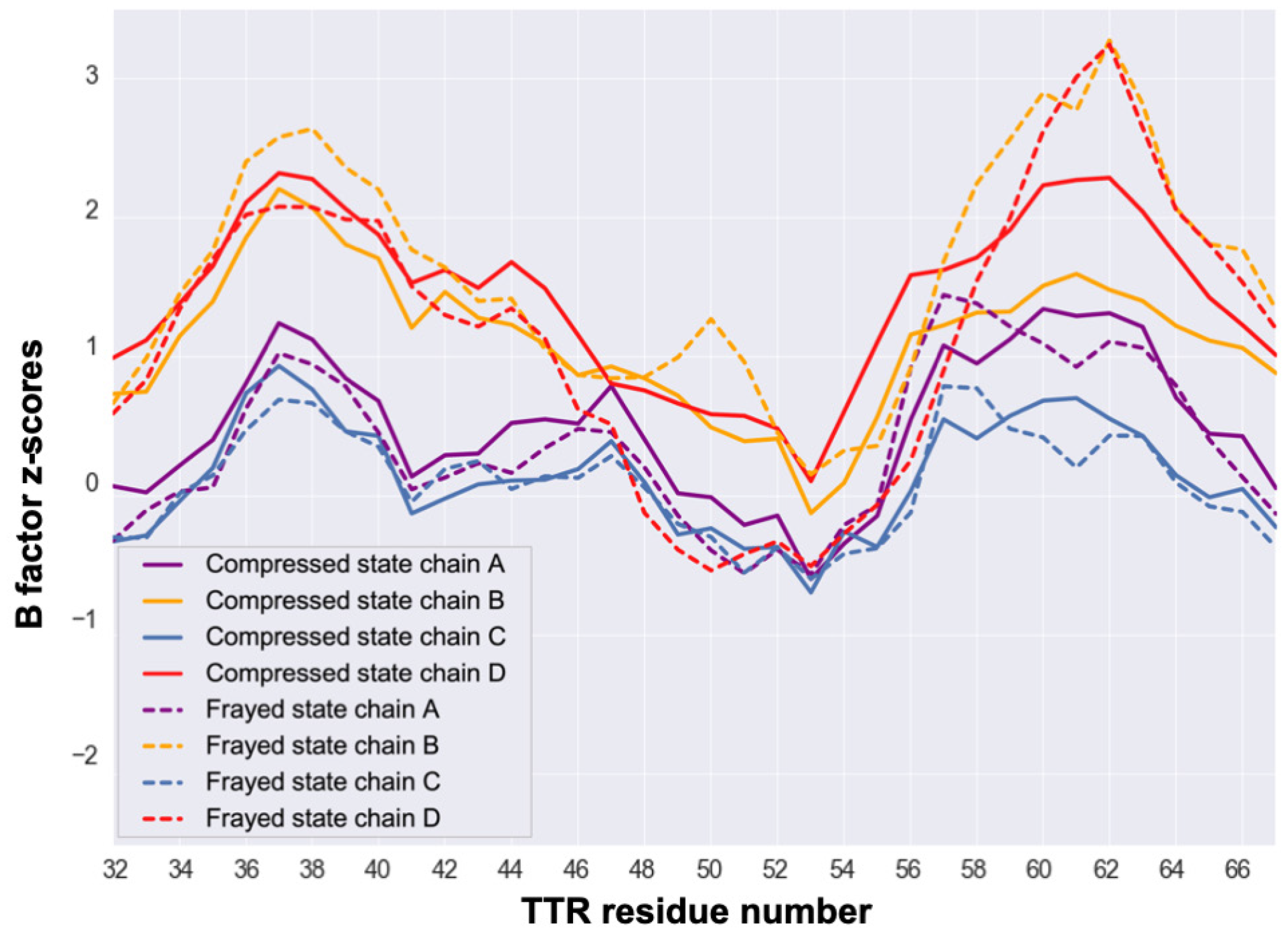
B-factor Z-scores for the compressed and frayed conformations of (biarylamine-FT_2_-WT)_1_(C10A)_3_ TTR. In both compressed and frayed states, chains B and D form the unoccupied binding pocket. The compressed and frayed states from (biarylamine-FT_2_-WT)_1_(C10A)_3_ TTR were resolved to 3.4 and 4.1 Å ; this precludes direct comparison of B factors, so instead we present B factor Z-scores, as suggested in^60^. Z-scores are only presented for residue numbers and chains in areas that appear disordered in the frayed state. Z-scores for the same residues in the compressed state are presented for comparison.

**Supplementary Figure 22.**
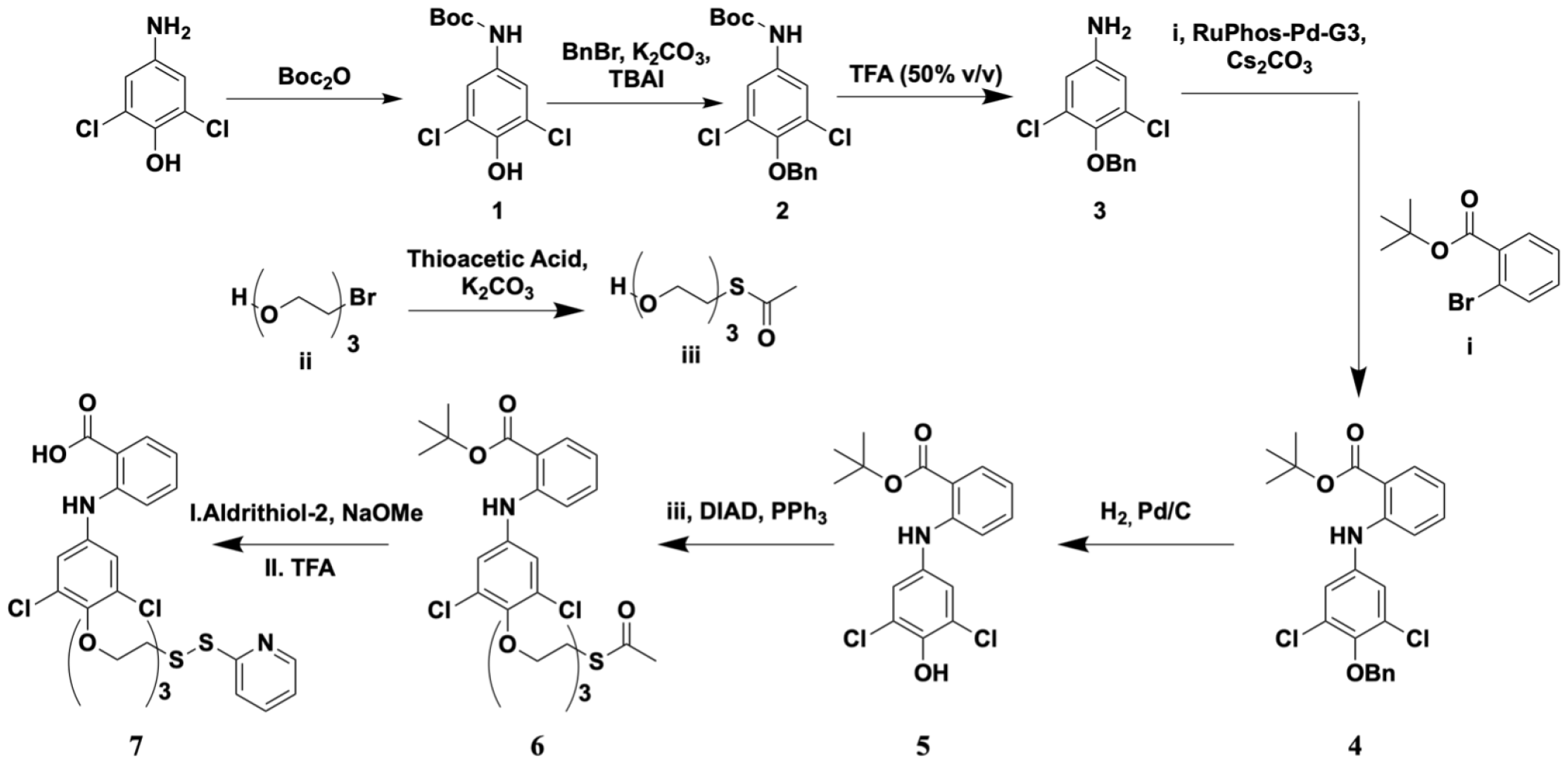
Synthetic scheme for 2-((3,5-dichloro-4-(2-(2-(2-(pyridin-2-yldisulfaneyl)ethoxy)ethoxy)ethoxy)phenyl) amino)benzoate.

**Supplementary Figure 23.**
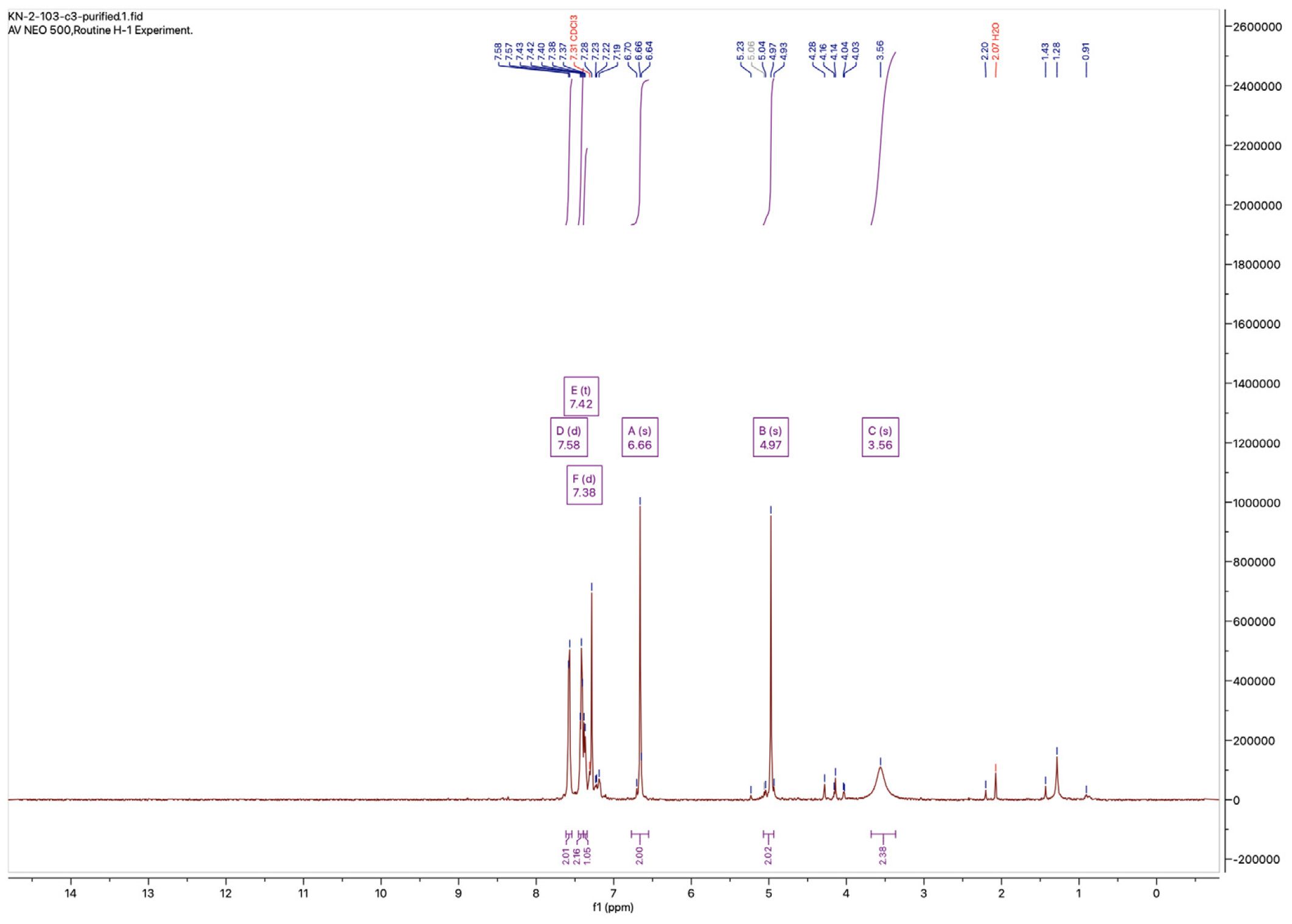
4-Benzyloxy-3,5-dichloroaniline NMR spectrum.

**Supplementary Figure 24.**
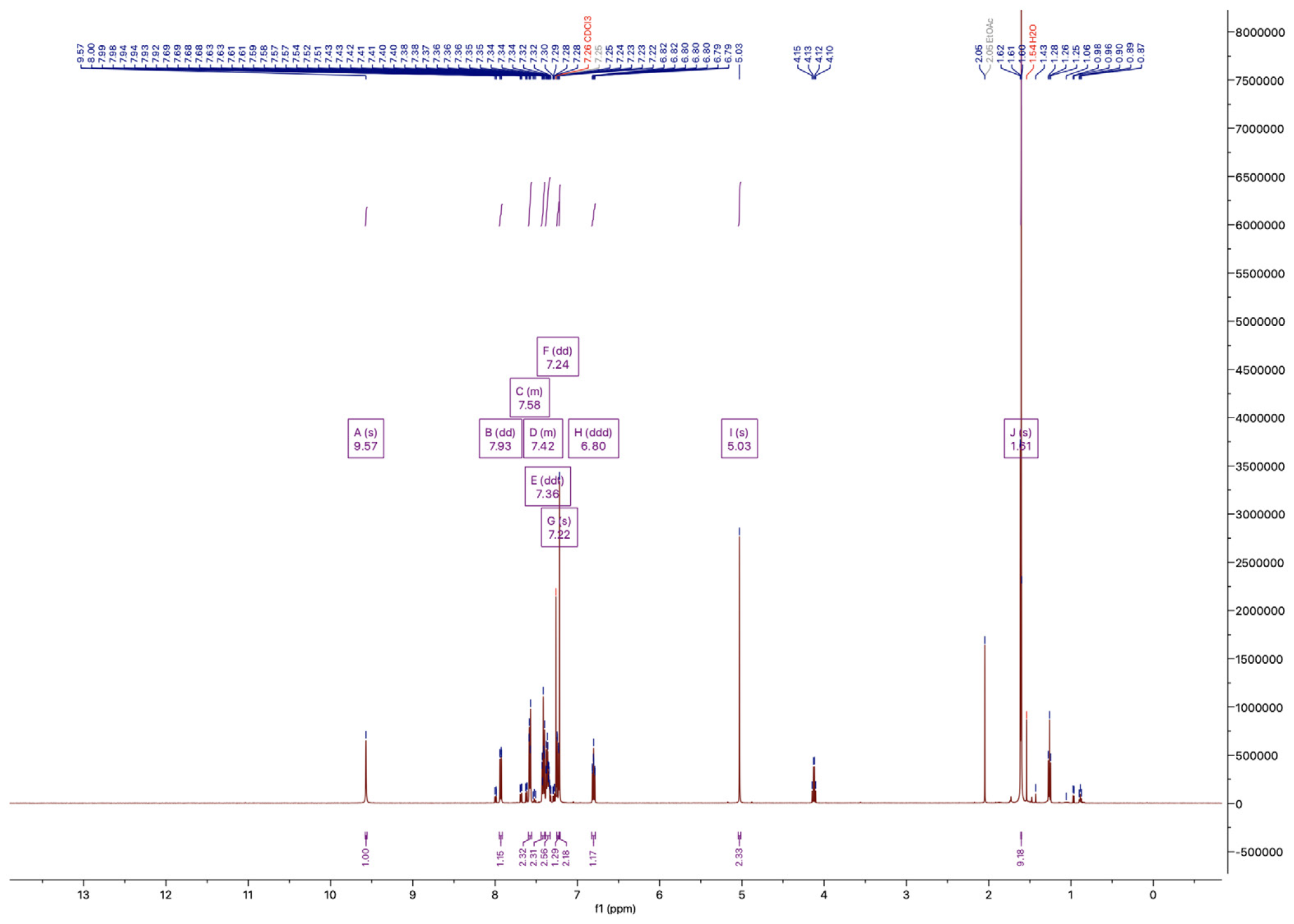
*tert*-Butyl 2-((4-(benzyloxy)-3,5-dichlorophenyl)amino)benzoate NMR spectrum.

**Supplementary Figure 25.**
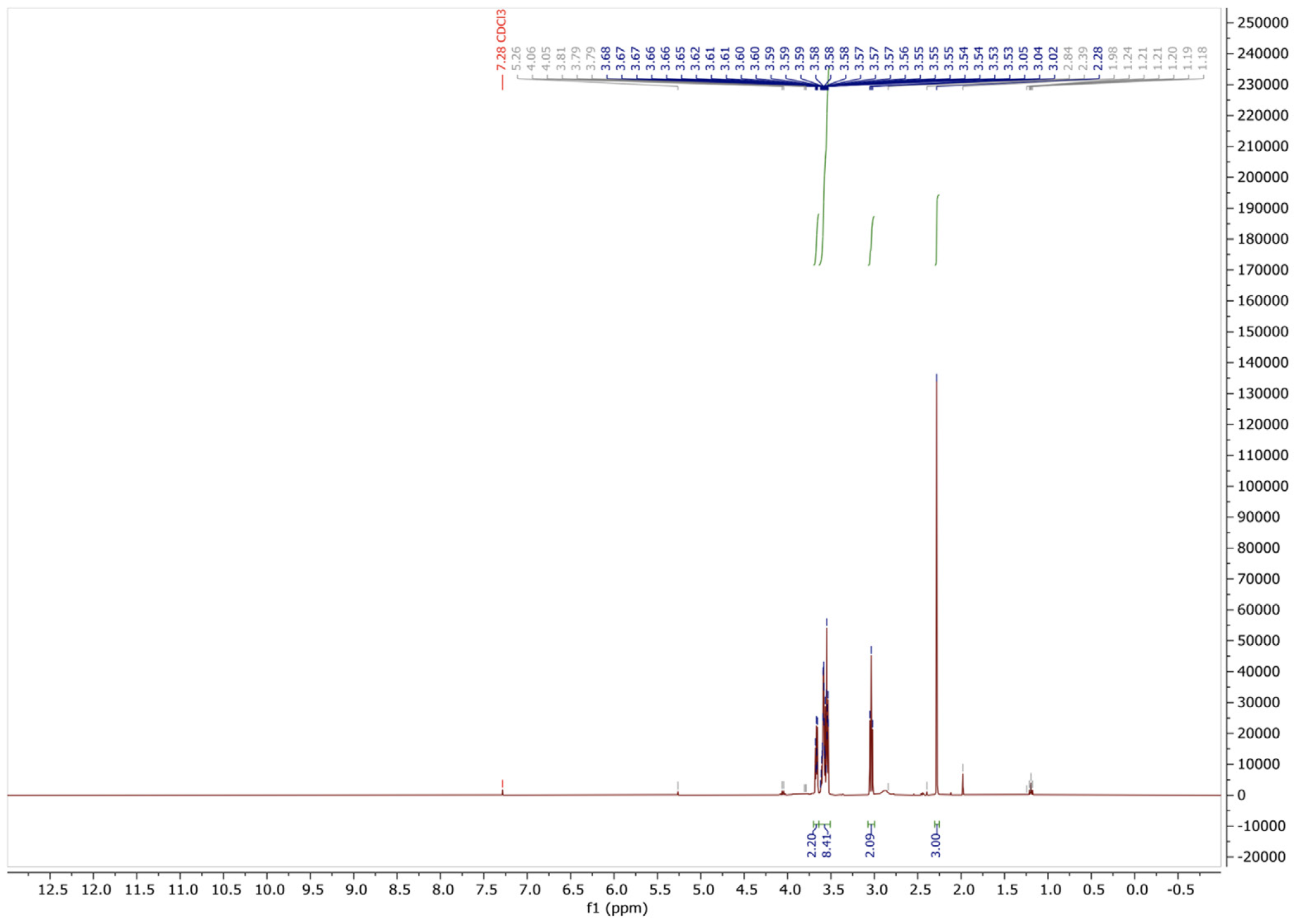
S-(2-(2-(2-hydroxyethoxy)ethoxy)ethyl) ethanethioate NMR spectrum.

**Supplementary Figure 26.**
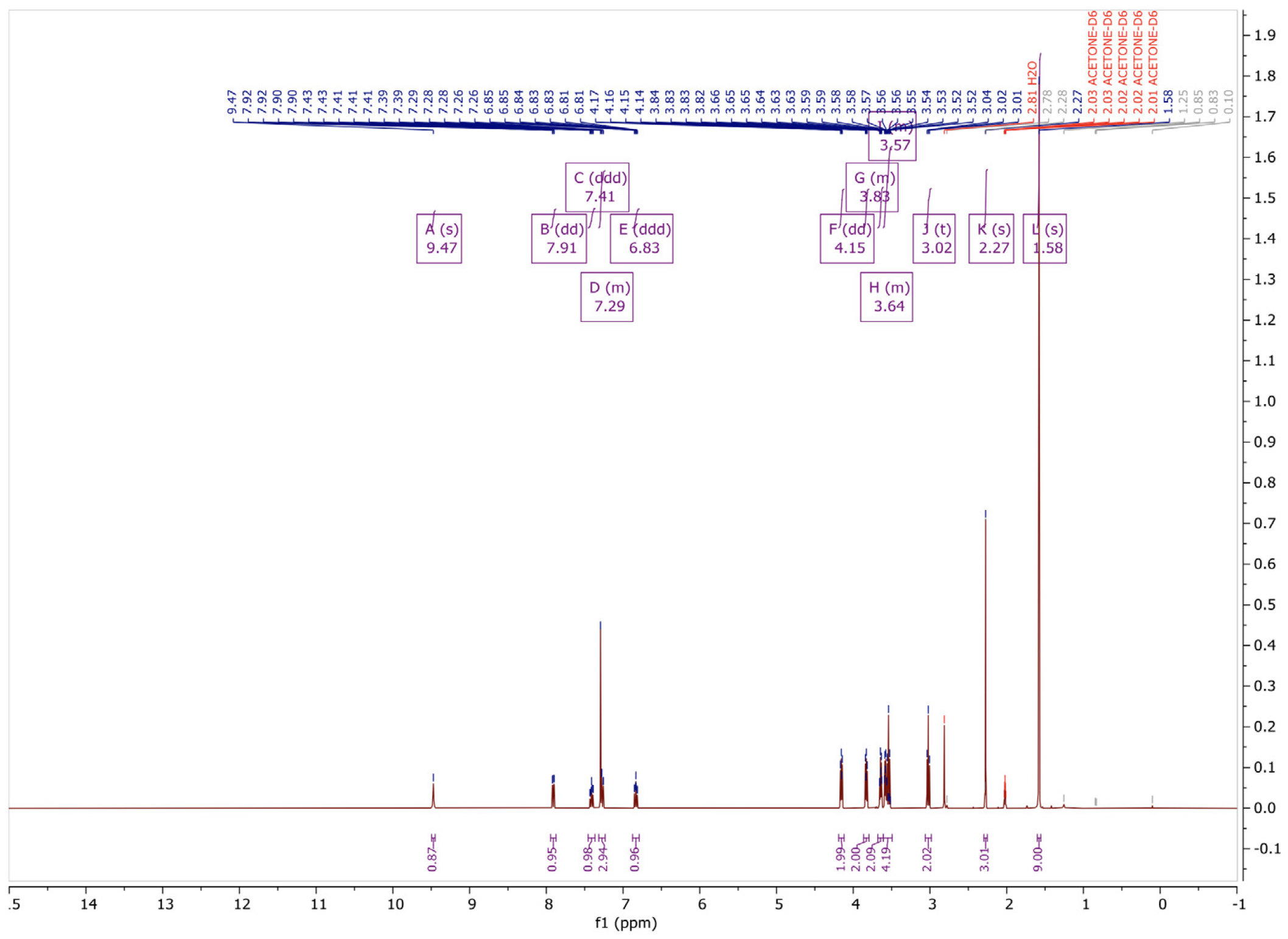
*tert*-butyl 2-((4-(2-(2-(2-(acetylthio)ethoxy)ethoxy)ethoxy)-3,5-dichlorophenyl)amino)benzoate NMR spectrum.

**Supplementary Figure 27.**
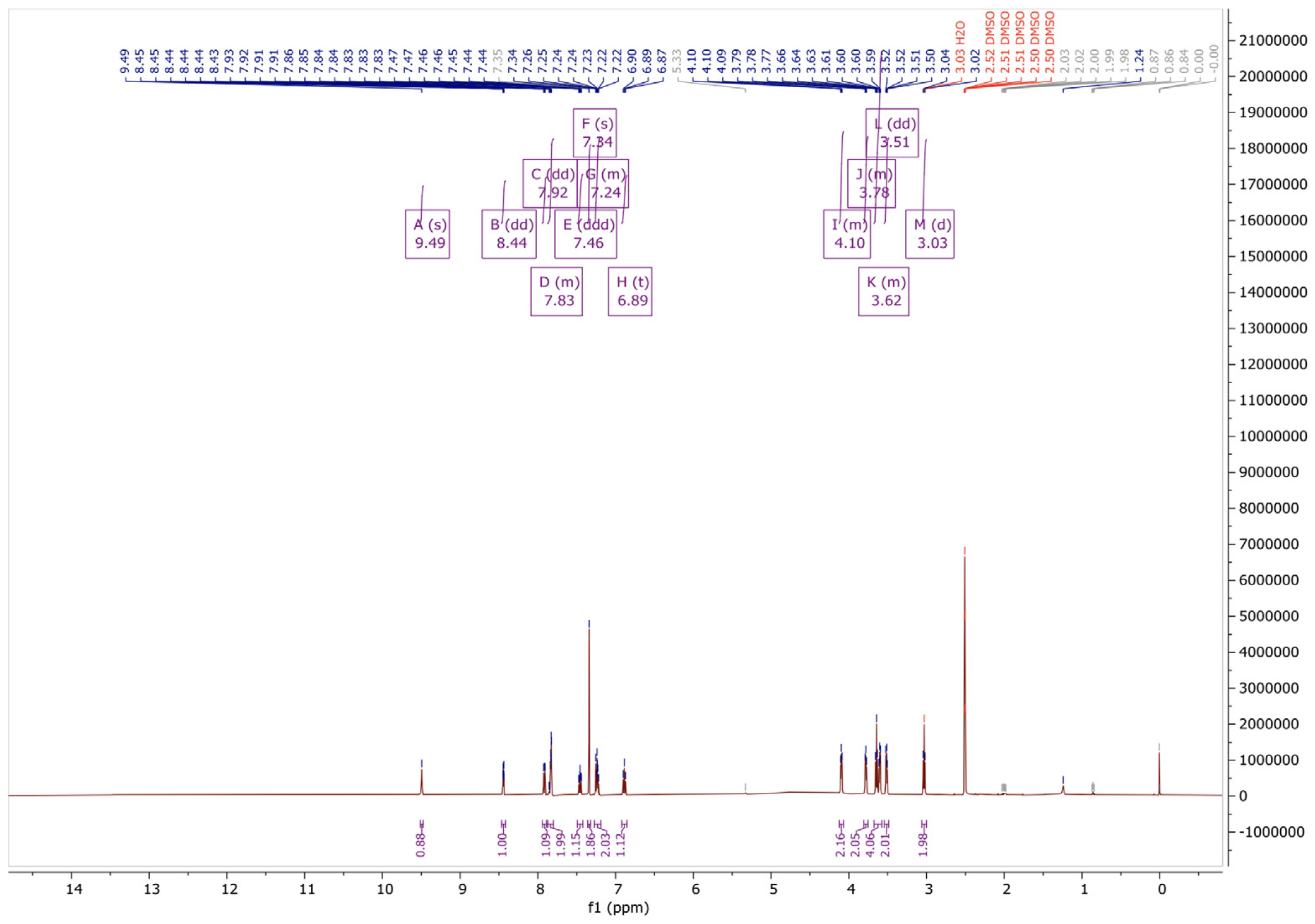
2-((3,5-dichloro-4-(2-(2-(2-(pyridin-2-yldisulfaneyl)ethoxy)ethoxy)ethoxy)phenyl)amino)benzoate (Biarylamine 7) NMR spectrum.

**Supplementary Figure 28.**
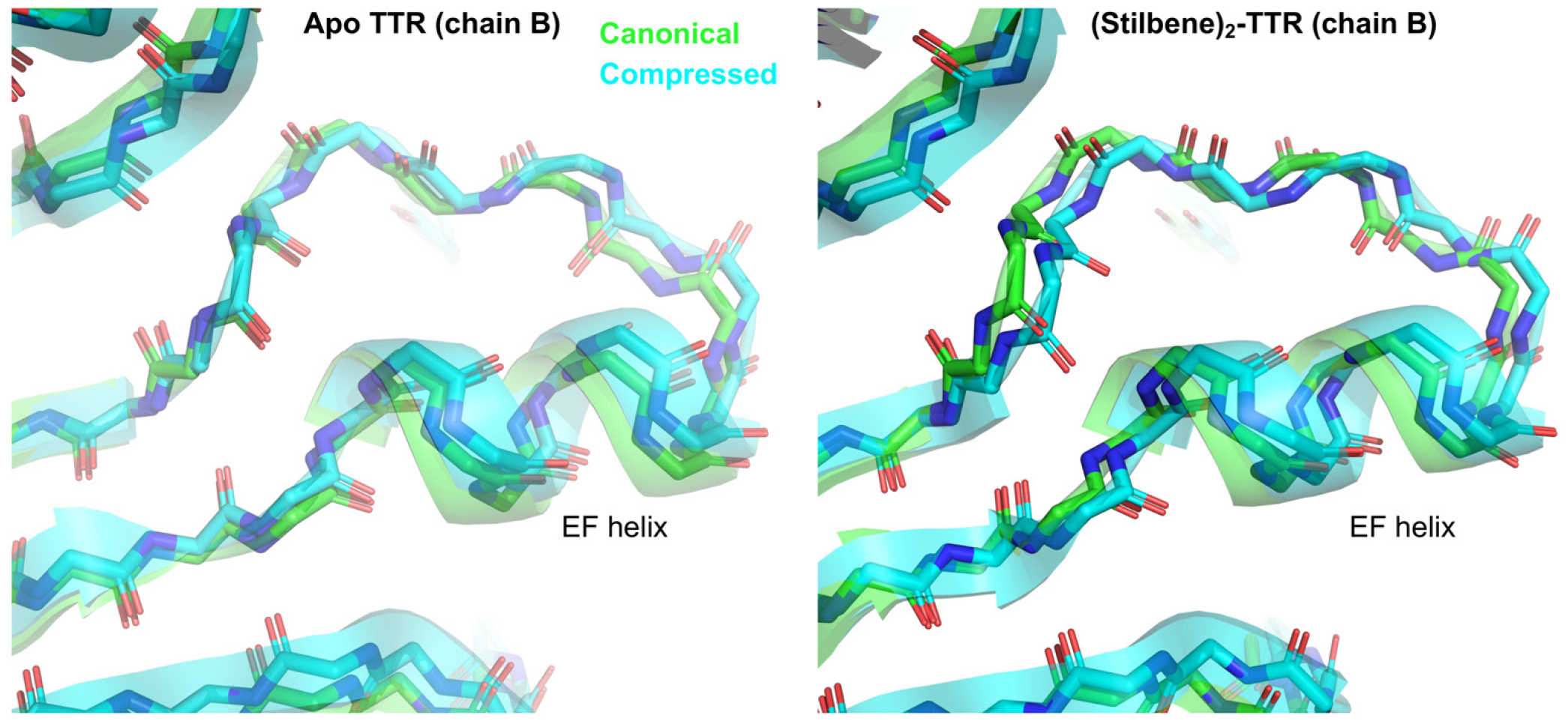
Elongation of the EF helix in the compressed conformation, compared to the canonical conformation. The EF helix is positioned further outward in the compressed conformation (cyan) than in the canonical conformation (green). This holds true whether ligand is present or not. Only chain B is displayed for simplicity, since all chains undergo similar changes in both samples.

**Supplementary Figure 29.**
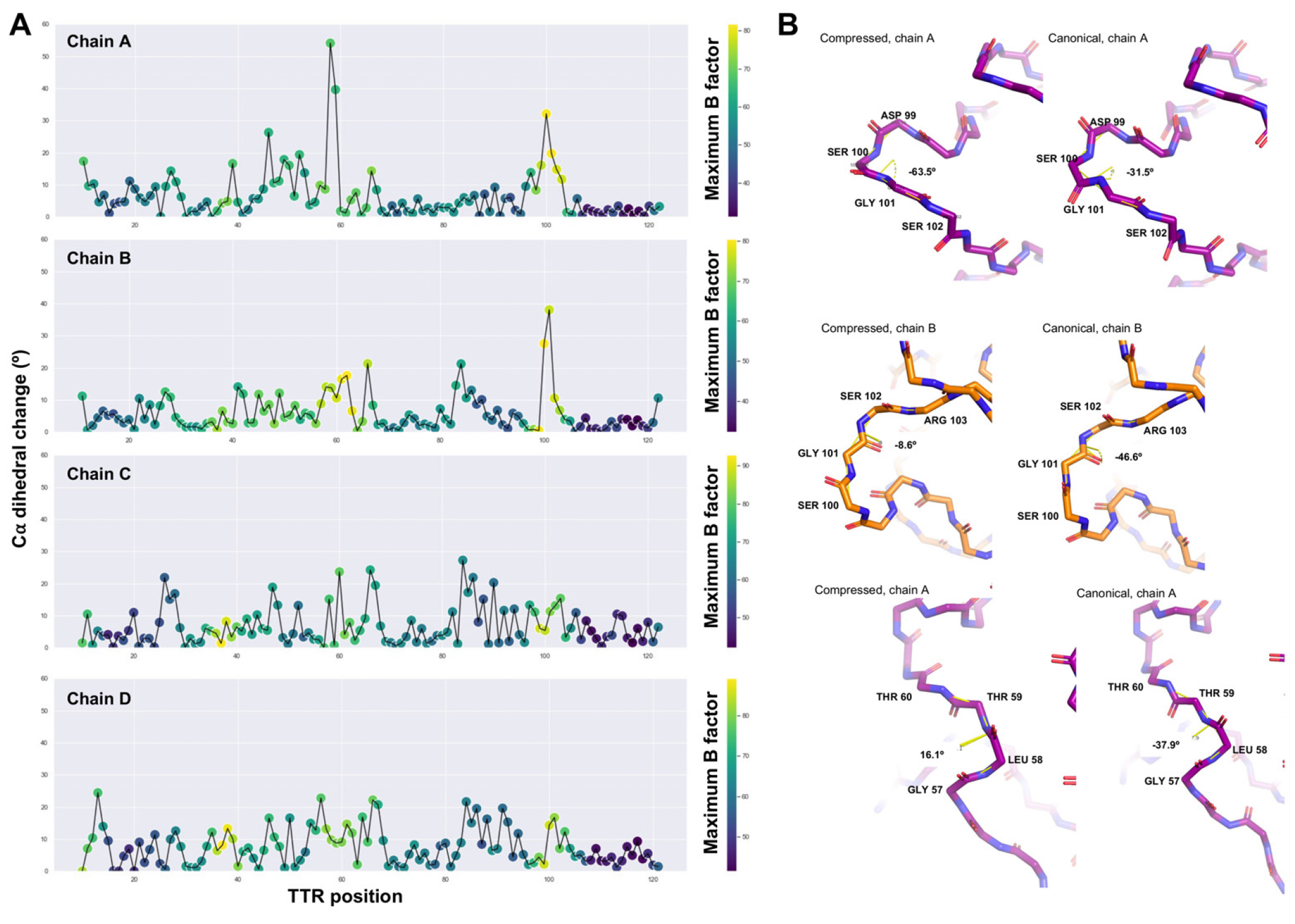
Backbone angle changes between the compressed and canonical (Stilbene)_2_-TTR conformations. (**A**)Per-position Cα dihedral changes, colored by the highest B-factor value between the two compared models. (**B**) Backbone-only representation of positions with the highest dihedral angle changes: Ser100 from chain A, Gly101 from chain B, and Leu58 from chain A. The Cα dihedral angle values are shown.

**Supplementary Figure 30.**
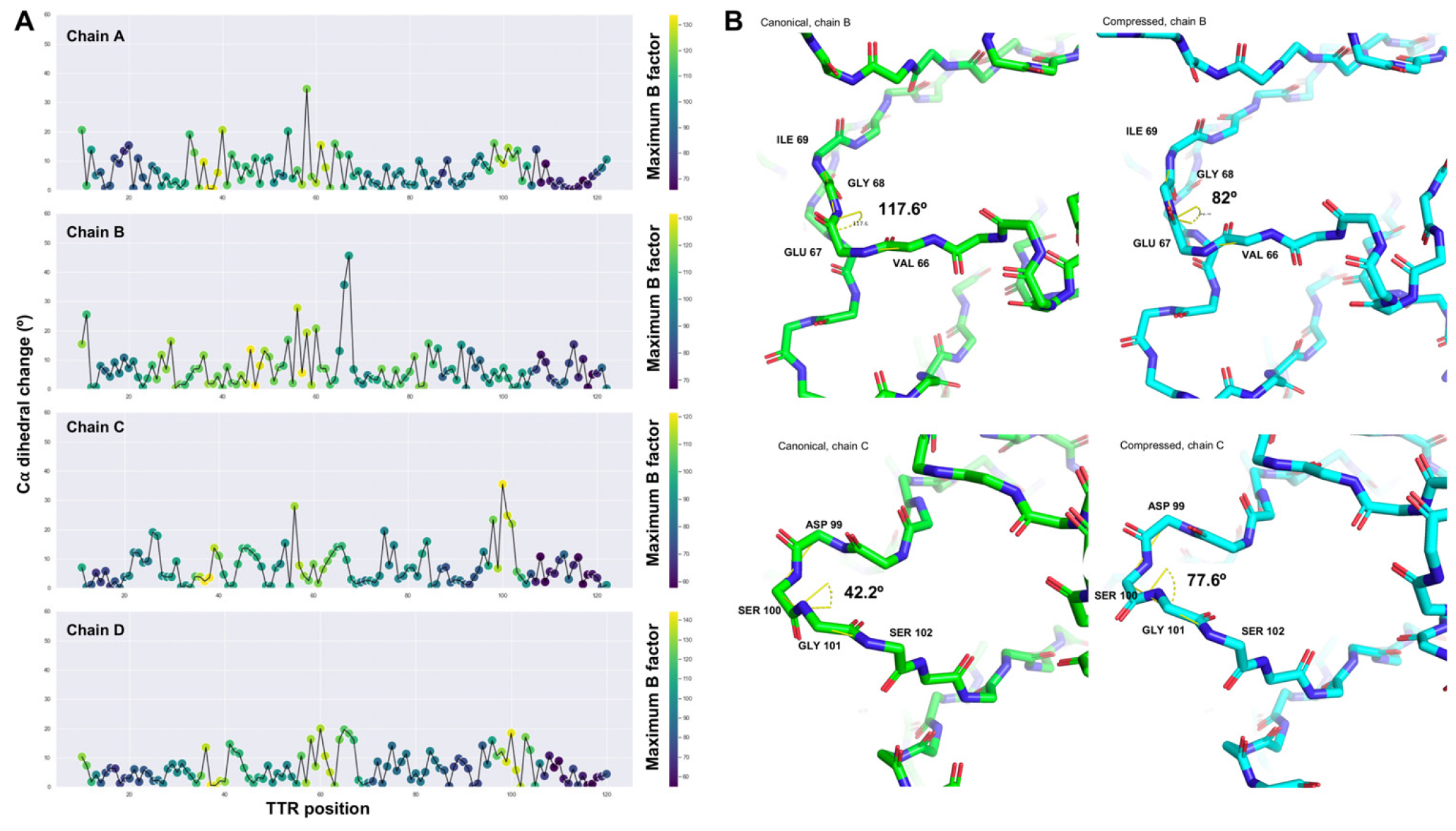
Backbone angle changes between the unliganded canonical and compressed TTR conformations. (**A**)Per-position Cα dihedral changes, colored by the highest B-factor value between the two compared models. (**B**) Backbone-only representation of positions with the highest dihedral angle changes: Glu67 from chain B and Ser100 from chain C. The Cα dihedral angle values are shown.

**Supplementary Figure 31.**
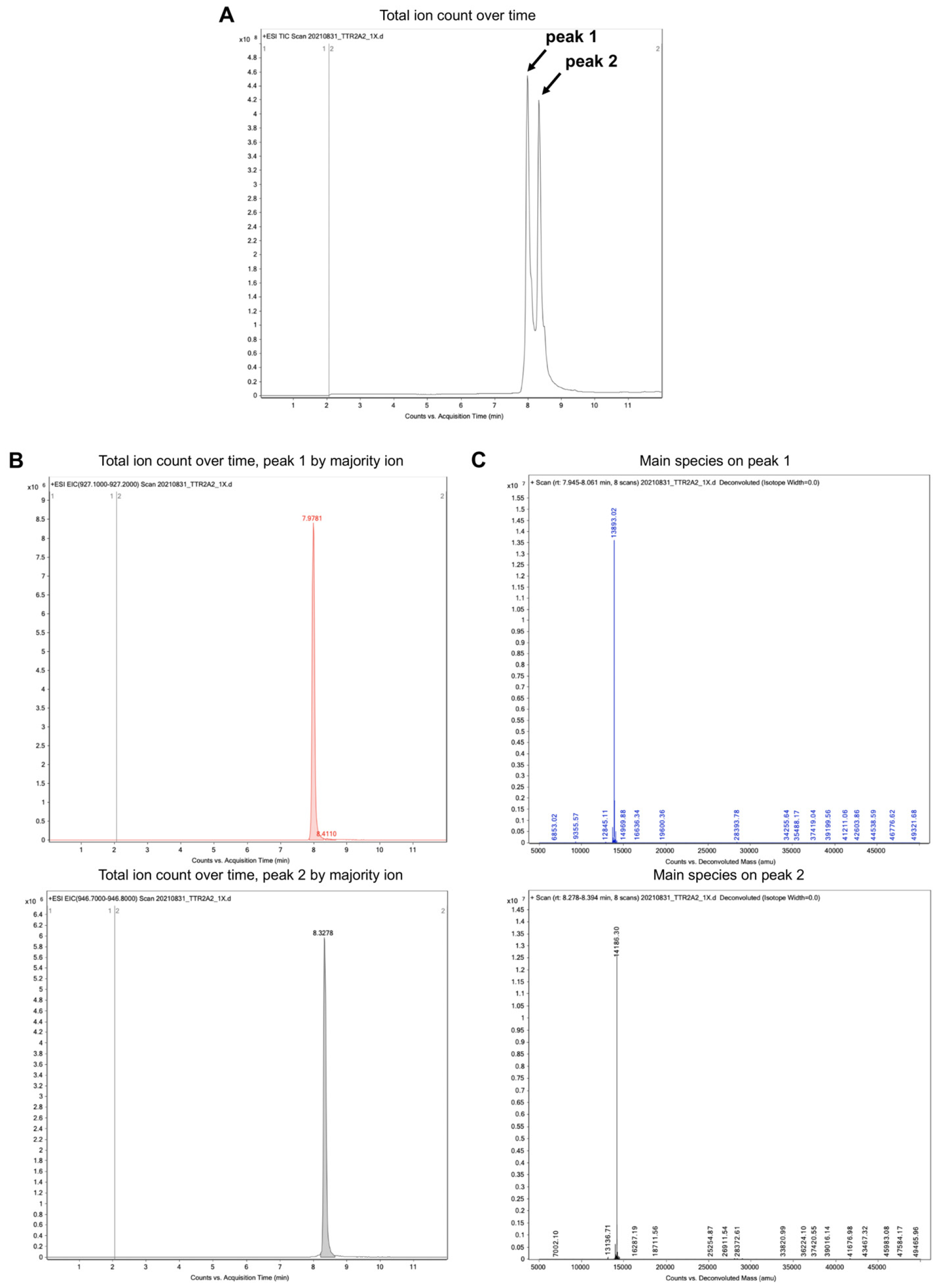
(previous page). Mass spectrometry analysis of (Stilbene)_2_-TTR conjugate. (**A**)Two peaks are clearly distinguishable in the total ion count as a function of elution time plot. (**B**) The relative area under the curve for majority ion peak 1 is 43099823 and for majority ion peak 2 is 32270904. (**C**) Main species in peak 1 and 2. The expected mass for unlabeled TTR monomers (peak 1) is 13,883 Da (observed 13,893), for stilbene-conjugated TTR monomers (peak 2) is 14,185 Da (observed 14,186).

**Supplementary Figure 32.**
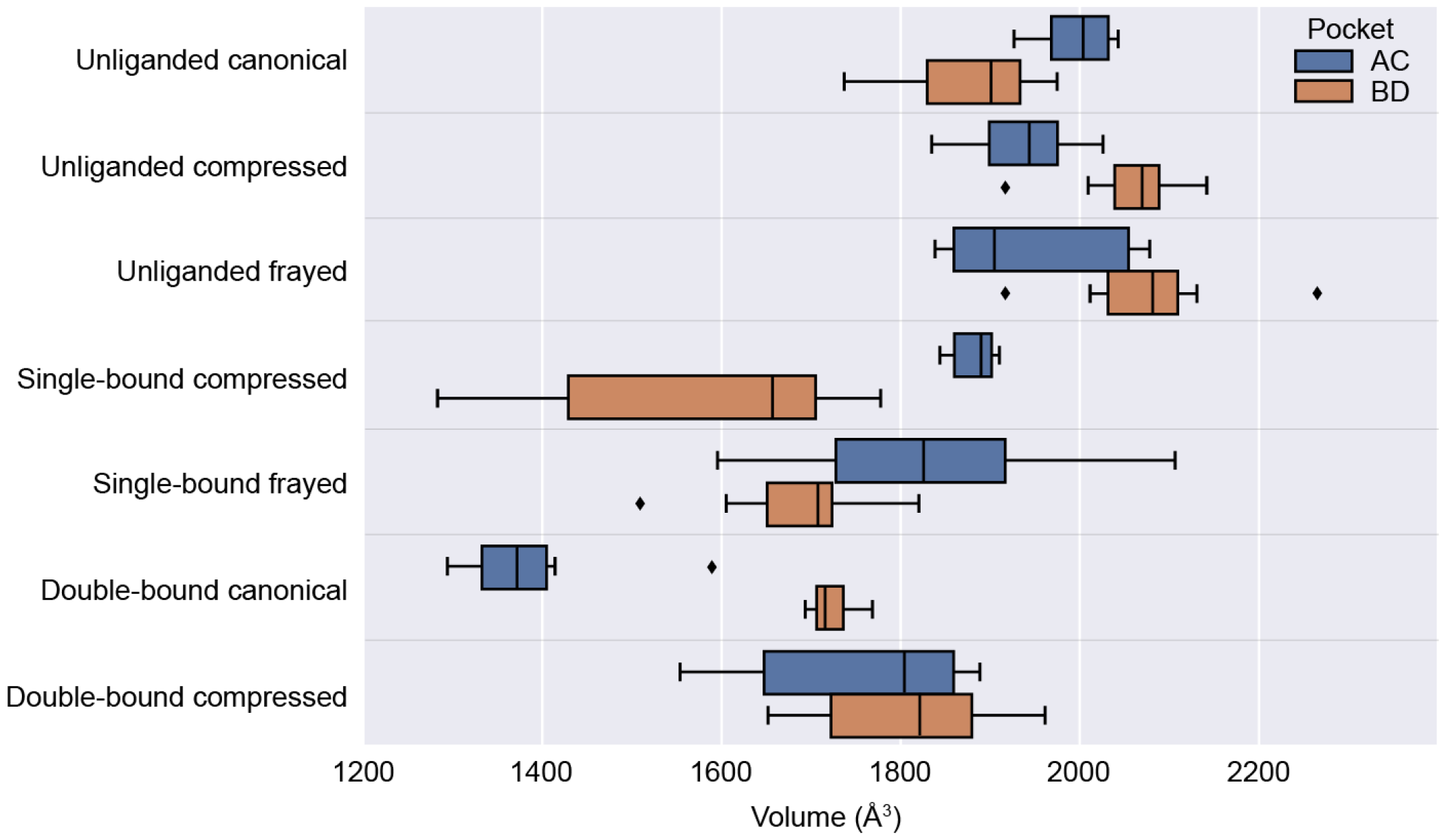
Fpocket analysis of TTR binding pockets. A box and whisker plot is shown for each of the binding pockets for all of the conformational states of TTR reported in this study. Ten atomic models were generated for each cryo-EM reconstruction using Rosetta to model any missing side-chains, and Fpocket was used to measure the volume of each binding site (see Methods). The central line in the box plot is the median and the rectangles extend to the central quartiles. The whiskers extend to contain the rest of the data, except for points that are further than 1.5 times the inter-quartile range from the closest hinge, which are considered outliers (denoted by diamonds). The mean and standard deviation of each binding pocket for the conformational states are as follows (AC, BD pocket, respectively in Å ^3^): unliganded canonical: 1997 +/-36, 1880 +/-69; unliganded compressed: 1935 +/-59, 2061 +/-59; unliganded frayed: 1945 +/-91, 2080 +/-85; single-bound compressed: 1883 +/-22, 1582 +/-170; single-bound frayed: 1827 +/-139, 1691 +/-84; double-bound canonical: 1384 +/-79, 1723 +/-21; double-bound compressed: 1681 +/-311, 1815 +/-91.

**Supplementary Figure 33.**
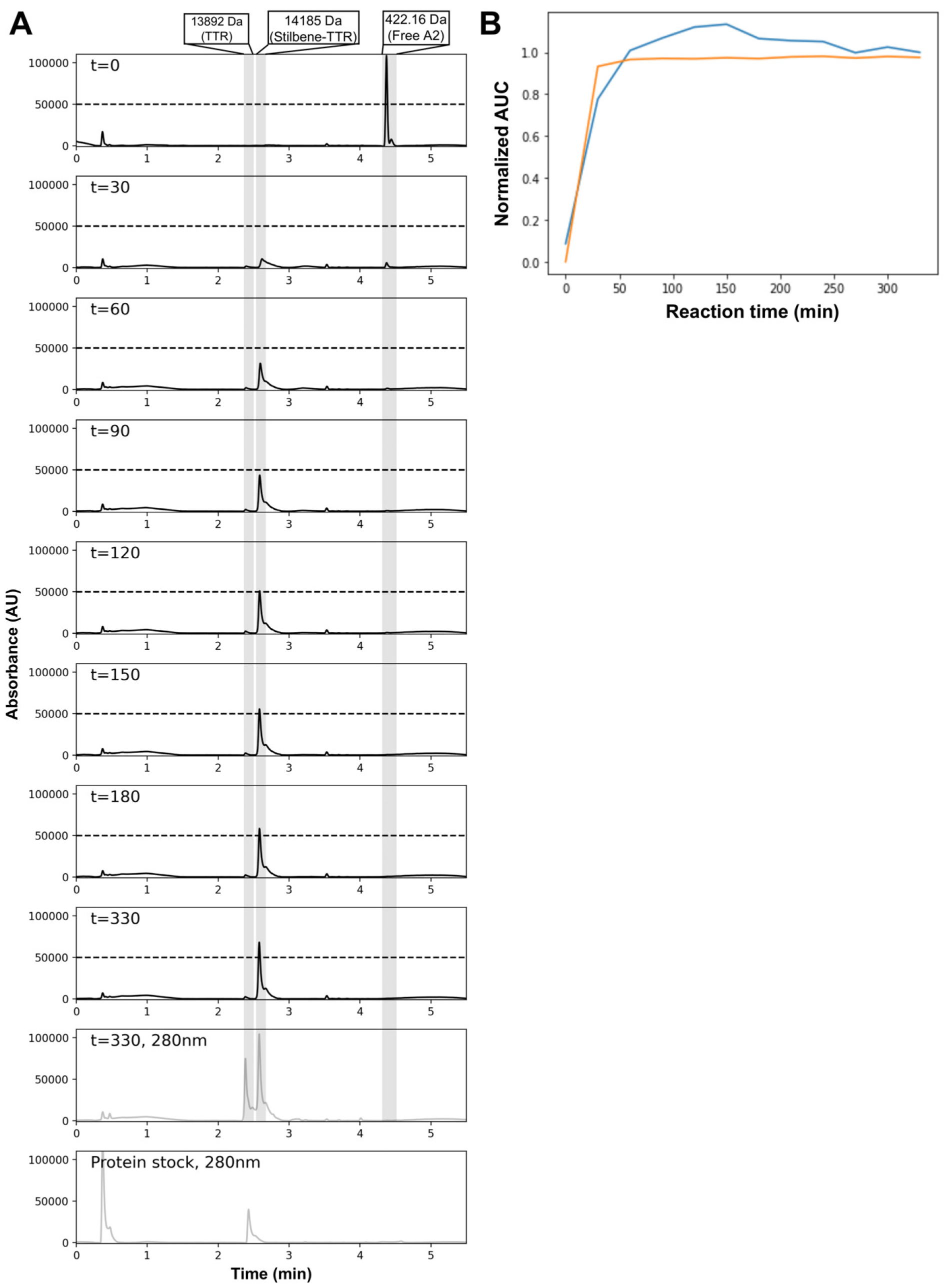
(previous page). Time course of A2 reaction with TTR affording (Stilbene)_2_-TTR conjugate, followed by LC-MS. (**A**)The top eight panels are a 5 Å L reaction aliquot chromatography, followed by UV absorbance at 330 nm. The ninth panel corresponds to the last timepoint (330 minutes), but protein absorbance at 280 nm is plotted. The bottom panel is the protein stock, without A2. The main species mass detected by MS is labeled at the top and highlighted in all panels by a grey transparent bar. (**B**) Blue: Area under the curve for the (Stilbene)_2_-TTR peak as a function of reaction time, normalized by the (Stilbene)_2_-TTR peak intensity maximum. Orange: 1-AUC for the A2 peak as a function of reaction time, normalized by A2 peak intensity maximum.

**Supplementary Figure 34.**
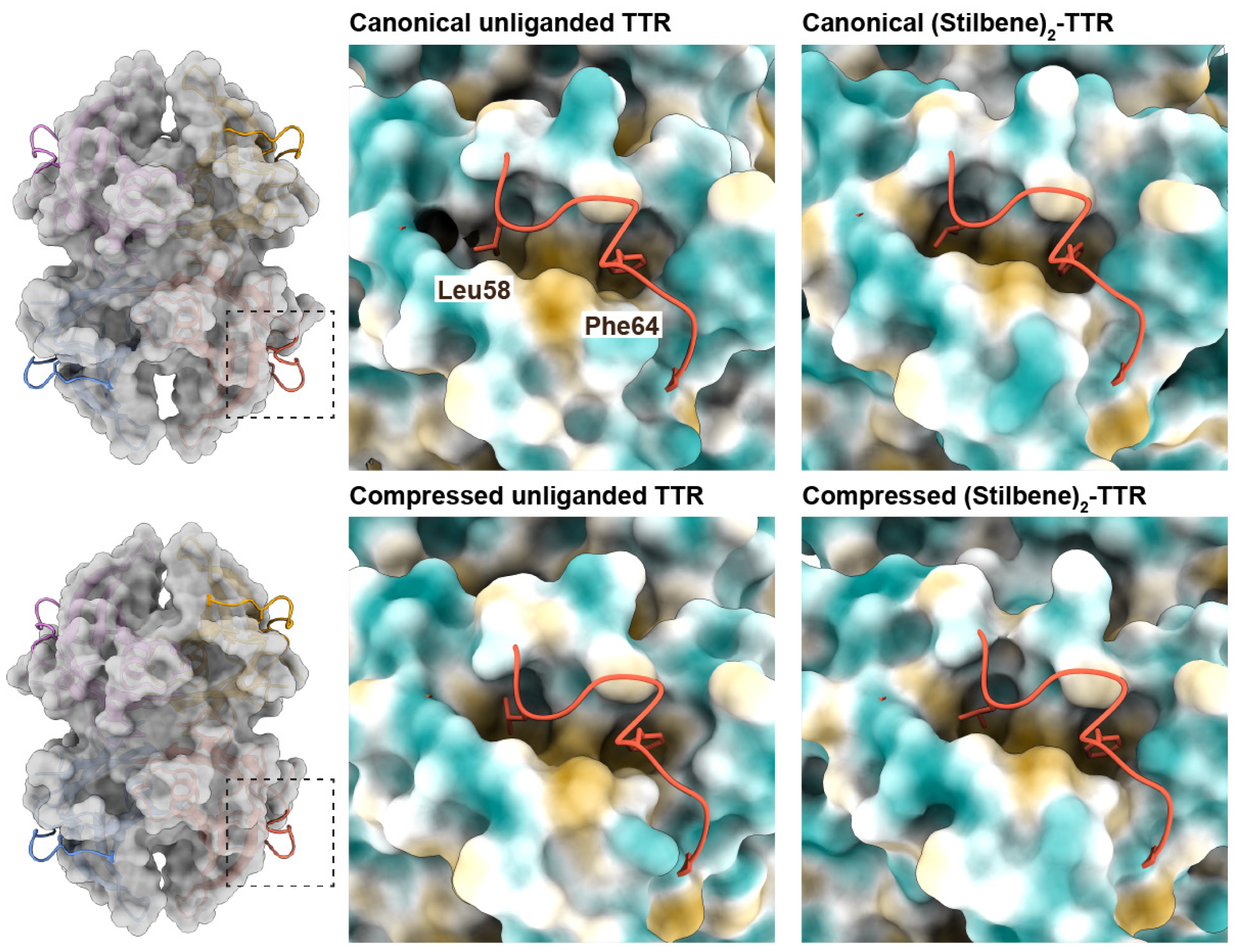
Conformational changes around Phe64 between the canonical and compressed TTR conformations. On the left, a transparent surface representation of the unliganded canonical (above) and compressed (below) conformations of TTR are shown with transparency over the the ribbon model. The surface corresponding to the D-E loop has been removed to highlight its location. On the right, the surface representation of TTR is colored according to hydrophobicity according to the Kyte-Doolittle scale, with brown representing hydrophobic surfaces and blue as hydrophilic. The D-E loop is shown as a ribbon with residues Phe64 and Leu58 shown as sticks, highlighting how these residues anchor the D-E loops to the rest of the structure by interacting with the hydrophobic core of each subunit. When TTR transitions from the canonical to compressed conformations, the hydrophobic pockets with which these residues interact become wider. We hypothesize this decreases the stability of the D-E loop interaction relative to the rest of the structure, enabling sampling of the frayed state.

**Supplementary Table 1.** (Stilbene)_2_-TTR conjugate crystallographic data collection and modeling statistics. Data collection and refinement statistics for the (Stilbene)_2_-TTR crystal structure. The structure was determined from one crystal. Values for the highest resolution shell are given in parentheses.

| **(Stilbene)<sub>2</sub>-TTR** |  |
| --- | --- |
| PDB ID | 8U52 |
| **Data Collection** |  |
| Beamline | ALS 5.0.1 |
| Space group | P2<sub>1</sub>2<sub>1</sub>2 |
| (a, b, c) (Å) | 42.96, 85.20, 63.75 |
| (α, β, γ) (°) | 90, 90, 90 |
| Resolution range (Å) | 42.60 – 1.50 (1.58 – 1.50) |
| Unique reflections | 36,591 (4,326) |
| Completeness (%) | 95.7 (79.8) |
| R<sub>sym</sub> | 0.029 (0.411) |
| R<sub>pim</sub> | 0.008 (0.136) |
| CC(1/2) | 1.00 (0.94) |
| I/σ (I) | 42.8 (4.9) |
| Redundancy | 12.4 (9.6) |
| **Refinement** |  |
| Resolution range (Å) | 42.64 – 1.50 |
| No. reflections - work | 34,712 |
| No. reflections - free | 1,672 |
| R<sub>work</sub> | 0.165 |
| R<sub>free</sub> | 0.194 |
| RMS bond length (Å) | 0.050 |
| RMS bond angle (°) | 2.89 |
| Mean B value (Å<sup>2</sup>) |  |
| overall | 25 |
| protein | 24 |
| ligand | 31 |
| water | 36 |
| Ramachandran favored (%) | 97.7 |
| Ramachandran allowed (%) | 100.0 |
| Clashscore | 4.9 |
| No. atoms |  |
| total | 2,040 |
| protein | 1,832 |
| ligand | 44 |
| water | 164 |

**Supplementary Table 2.** Unliganded TTR canonical states cryo-EM data collection, analysis, and modeling statistics.

|  | Canonical | Compressed | Frayed |
| --- | --- | --- | --- |
| PDB ID | 8VE2 | 8VE3 | 8VE4 |
| EMDB ID | EMD-43162 | EMD-43163 | EMD-43164 |
| Data Collection |  |  |  |
| Microscope | Talos Arctica |  |  |
| Camera | K2 Summit (counting) |  |  |
| Magnification (nominal/at detector) | 73,000/88,968 |  |  |
| Voltage (kV) | 200 |  |  |
| Data acquisition software | Leginon |  |  |
| Exposure navigation | image shift to 16 |  |  |
| Total electron exposure (e<sup>-</sup>/Å<sup>2</sup>) | 55 |  |  |
| Exposure rate (e<sup>-</sup>/pixel/sec) | 2.3 |  |  |
| Frame length (ms) | 150 |  |  |
| Number of frames | 50 |  |  |
| Pixel size (Å/pixel)* | 0.562 |  |  |
| Defocus range (μm) | -0.8 to -1.5 |  |  |
| Micrographs collected (no.) | 3,739 |  |  |
| Data analysis |  |  |  |
| Total extracted picks (no.) | 1,950,142 |  |  |
| Particles used for 3D (no.) | 1,301,827 |  |  |
| Final Particles (no.) | 335,266 | 130,181 | 202,700 |
| Symmetry | C1 | C1 | C1 |
| Resolution (global) (Å) |  |  |  |
| FSC 0.5 (unmasked / masked) | 4.1 / 3.5 | 4.1 / 3.5 | 4.1 / 3.5 |
| FSC 0.143 (unmasked / masked) | 3.7 / 3.3 | 3.7 / 3.3 | 3.7 / 3.3 |
| Local resolution range (Å) | 3.2 - 3.9 | 3.2 - 3.9 | 3.2 - 3.9 |
| 3DFSC Sphericity (%) | 85.7 | 88.6 | 91.7 |
| Accuracy of rotations / translations | 3.17 / 0.72 | 2.81 / 0.60 | 3.02 / 0.66 |
| Map sharpening *B*-factor (Å<sup>2</sup>) | -60 | -55 | -55 |
| Model composition |  |  |  |
| Chains | 4 | 4 | 4 |
| Non-hydrogen atoms | 3,105 | 3,210 | 3,148 |
| Protein residues | 465 | 466 | 465 |
| Model Refinement* |  |  |  |
| Refinement package | Phenix | Phenix | Phenix |
| MapCC (volume/mask) | 0.75 / 0.77 | 0.77 / 0.78 | 0.74 / 0.75 |
| B-factors (Å<sup>2</sup>) |  |  |  |
| Protein residues | 95.0 | 91.4 | 89.6 |
| R.m.s. deviations |  |  |  |
| Bond lengths (Å) | 0.00 | 0.00 | 0.00 |
| Bond angles (°) | 0.46 | 0.48 | 0.47 |
| Validation |  |  |  |
| Map-to-model FSC 0.5 | 3.7 | 3.5 | 3.8 |
| Ramachandran plot |  |  |  |
| Favored (%) | 98.91 | 96.29 | 98.69 |
| Allowed (%) | 1.09 | 3.71 | 1.31 |
| Outliers (%) | 0.0 | 0.0 | 0.0 |
| MolProbity score | 1.27 | 1.83 | 1.38 |
| Clashscore | 5.05 | 11.69 | 6.96 |
| Poor rotamers (%) | 0.00 | 0.00 | 0.00 |
| CaBLAM outliers (%) | 1.78 | 1.78 | 1.34 |
| EMRinger score<sup>64</sup> | 4.10 | 5.02 | 4.32 |
| Qscore | 0.63 | 0.65 | 0.62 |

**Supplementary Table 3.** Double-bound TTR conjugate states cryo-EM data collection, analysis, and modeling statistics.

|  | Canonical | Compressed |
| --- | --- | --- |
| PDB ID | 8VE1 | 8VE0 |
| EMDB ID | EMD-43161 | EMD-43160 |
| Data Collection |  |  |
| Microscope | Talos Arctica |  |
| Camera | K2 Summit (counting) |  |
| Magnification (nominal/at detector) | 73,000/88,968 |  |
| Voltage (kV) | 200 |  |
| Data acquisition software | Leginon |  |
| Exposure navigation | image shift to 16 |  |
| Total electron exposure (e<sup>-</sup>/Å<sup>2</sup>) | 55 |  |
| Exposure rate (e<sup>-</sup>/pixel/sec) | 2.3 |  |
| Frame length (ms) | 150 |  |
| Number of frames | 50 |  |
| Pixel size (Å/pixel)* | 0.562 |  |
| Defocus range (μm) | -0.8 to -1.5 |  |
| Micrographs collected (no.) | 4,536 |  |
| Data analysis |  |  |
| Total extracted picks (no.) | 1,085,771 |  |
| Particles used for 3D (no.) | 1,030,175 |  |
| Final Particles (no.) | 289,981 | 105,961 |
| Symmetry | C1 | C1 |
| Resolution (global) (Å) |  |  |
| FSC 0.5 (unmasked / masked) | 3.8 / 3.2 | 4.1 / 3.4 |
| FSC 0.143 (unmasked / masked) | 3.0 / 2.7 | 3.3 / 3.1 |
| Local resolution range (Å) | 2.7 - 3.3 | 3.1 - 3.8 |
| 3DFSC Sphericity (%) | 96.9 | 95.1 |
| Accuracy of rotations / translations | 2.17 / 0.49 | 2.78 / 0.64 |
| Map sharpening *B*-factor (Å<sup>2</sup>) | -60 | -60 |
| Model composition |  |  |
| Chains | 5 | 5 |
| Non-hydrogen atoms | 3,656 | 3,529 |
| Protein residues | 464 | 464 |
| Water | 27 | 0 |
| Ligands | A2K: 2 | A2K: 2 |
| Model Refinement* |  |  |
| Refinement package | Phenix | Phenix |
| MapCC (volume/mask) | 0.83 / 0.85 | 0.77 / 0.79 |
| B-factors (Å<sup>2</sup>) |  |  |
| Protein residues | 42.1 | 61.2 |
| Ligand | 30.67 | 47.79 |
| Water | 32.67 | - |
| R.m.s. deviations |  |  |
| Bond lengths (Å) | 0.00 | 0.00 |
| Bond angles (°) | 0.47 | 0.47 |
| Validation |  |  |
| Map-to-model FSC 0.5 | 2.7 | 3.1 |
| Ramachandran plot |  |  |
| Favored (%) | 97.79 | 99.78 |
| Allowed (%) | 2.21 | 0.22 |
| Outliers (%) | 0.0 | 0.0 |
| MolProbity score | 1.1 | 1.35 |
| Clashscore | 2.7 | 6.0 |
| Poor rotamers (%) | 0.00 | 0.00 |
| CaBLAM outliers (%) | 1.78 | 0.67 |
| EMRinger score | 5.84 | 4.24 |
| Qscore | 0.76 | 0.68 |

**Supplementary Table 4.** Single-bound (biarylamine-FT_2_-WT)_1_(C10A)_3_ TTR states cryo-EM data collection, analysis, and modeling statistics.

|  | Compressed | Frayed |
| --- | --- | --- |
| PDB ID | 8VE5 | 8VE6 |
| EMDB ID | EMD-43165 | EMD-43166 |
| Data Collection |  |  |
| Microscope | Talos Arctica |  |
| Camera | K2 Summit (counting) |  |
| Magnification (nominal/at detector) | 73,000/88,968 |  |
| Voltage (kV) | 200 |  |
| Data acquisition software | Leginon |  |
| Exposure navigation | image shift to 64 |  |
| Total electron exposure (e<sup>-</sup>/Å<sup>2</sup>) | 55 |  |
| Exposure rate (e<sup>-</sup>/pixel/sec) | 2.3 |  |
| Frame length (ms) | 150 |  |
| Number of frames | 50 |  |
| Pixel size (Å/pixel)* | 0.562 |  |
| Defocus range (μm) | -0.8 to -1.5 |  |
| Micrographs collected (no.) | 7,939 |  |
| Data analysis |  |  |
| Total extracted picks (no.) | 2,650,295 |  |
| Particles used for 3D (no.) | 2,432,152 |  |
| Final Particles (no.) | 168,443 | 110,253 |
| Symmetry | C1 | C1 |
| Resolution (global) (Å) |  |  |
| FSC 0.5 (unmasked / masked) | 4.6 / 4.2 | 5.2 / 4.5 |
| FSC 0.143 (unmasked / masked) | 3.8 / 3.4 | 4.4 / 4.1 |
| Local resolution range (Å) | 3.4 - 4.0 | 4.0 - 4.8 |
| 3DFSC Sphericity (%) | 97.0 | 81.0 |
| Accuracy of rotations / translations | 2.82 / 0.67 | 3.31 / 0.85 |
| Map sharpening *B*-factor (Å<sup>2</sup>) | -70 | -80 |
| Model composition |  |  |
| Chains | 5 | 5 |
| Non-hydrogen atoms | 3,386 | 3,181 |
| Protein residues | 467 | 465 |
| Ligands | P2C: 1 | P2C: 1 |
| Model Refinement* |  |  |
| Refinement package | Phenix | Phenix |
| MapCC (volume/mask) | 0.79 / 0.81 | 0.79 / 0.81 |
| B-factors (Å<sup>2</sup>) |  |  |
| Protein residues | 95.9 | 95.9 |
| Ligand | 83.2 | 68.5 |
| R.m.s. deviations |  |  |
| Bond lengths (Å) | 0.00 | 0.00 |
| Bond angles (°) | 0.50 | 0.49 |
| Validation |  |  |
| Map-to-model FSC 0.5 | 3.8 | 4.3 |
| Ramachandran plot |  |  |
| Favored (%) | 97.82 | 96.28 |
| Allowed (%) | 2.18 | 3.72 |
| Outliers (%) | 0.0 | 0.0 |
| MolProbity score | 1.45 | 1.76 |
| Clashscore | 7.4 | 9.8 |
| Poor rotamers (%) | 0.00 | 0.00 |
| CaBLAM outliers (%) | 2.00 | 3.72 |
| EMRinger score | 4.97 | 4.64 |
| Qscore | 0.64 | 0.60 |

